# Deep-learning-based design of an orthogonal self-labeling protein from K-Ras(G12C)

**DOI:** 10.64898/2026.09.27.754867

**Authors:** Jody Mou, Jing Wesley Leong, Sean Gao, Kaia Slaw, Michael McGill, Katherine A. Donovan, Rebecca J. Metivier, Franziska L. Sendker, Qi Kong, Diangen Lin, Azim M. Dharani, Xi Dawn Chen, Kevin M. Haigis, Eric S. Fischer, Fei Chen, Nicholas F. Polizzi

## Abstract

Self-labeling protein (SLP) tags enable versatile labeling of proteins in live cells, yet only two SLPs are commonly used, limiting multiplexing. Here, we repurposed the oncoprotein K-Ras(G12C) and its covalent inhibitors to create a third, orthogonal SLP system. We used a deep-learning-based approach to radically redesign the sequence of K-Ras(G12C) while preserving its covalent-inhibitor binding pocket. Our top design, LUCI-tag, is a 19 kDa, monomeric, thermostable SLP that rapidly and covalently reacts with commercially available K-Ras(G12C) inhibitors bearing diverse payloads. Unlike existing SLPs, LUCI-tag exhibited payload-agnostic labeling kinetics, outperforming HaloTag7 and SNAP-tag for a negatively charged payload. X-ray crystal structures of drug-bound and drug-free LUCI-tag showed the design was structurally accurate and contained a preorganized inhibitor-binding pocket that could explain its rapid labeling kinetics. Proteomics and cell-signaling experiments confirmed LUCI-tag is biologically inert. LUCI-tag enabled rapid, wash-free, live-cell imaging and simultaneous three-color multiplexed experiments with HaloTag7 and SNAP-tag. This work establishes LUCI-tag as an immediately useful orthogonal SLP and demonstrates that covalent drug–target pairs can be repurposed into a broadly applicable platform for protein labeling.

## Main text

Self-labeling protein (SLP) tags are powerful and accessible tools for visualizing and manipulating proteins in living systems (Fig. 1a). An SLP system consists of a genetically encoded protein tag and a covalently attached small-molecule probe carrying molecular cargo^1^. The high brightness and modular spectra of synthetic fluorophores make SLPs an attractive live-cell imaging strategy compared to fluorescent protein tags. In contrast to the broad palette of available fluorescent proteins, HaloTag^2–4^ and SNAP-tag^5–7^ are the only two commonly used orthogonal tags, limiting the number of targets that can be labeled simultaneously (Figs. 1b and S1). A reliable and orthogonal third SLP would enable greater multiplexing in contexts such as multi-color imaging, pulse-chase, and single-molecule experiments. However, attempts at creating new SLPs have struggled to overcome hurdles of slow labeling, limited probe permeability, off-target activity, and probe availability, limiting their immediate utility^8–10^.

**Figure 1.**
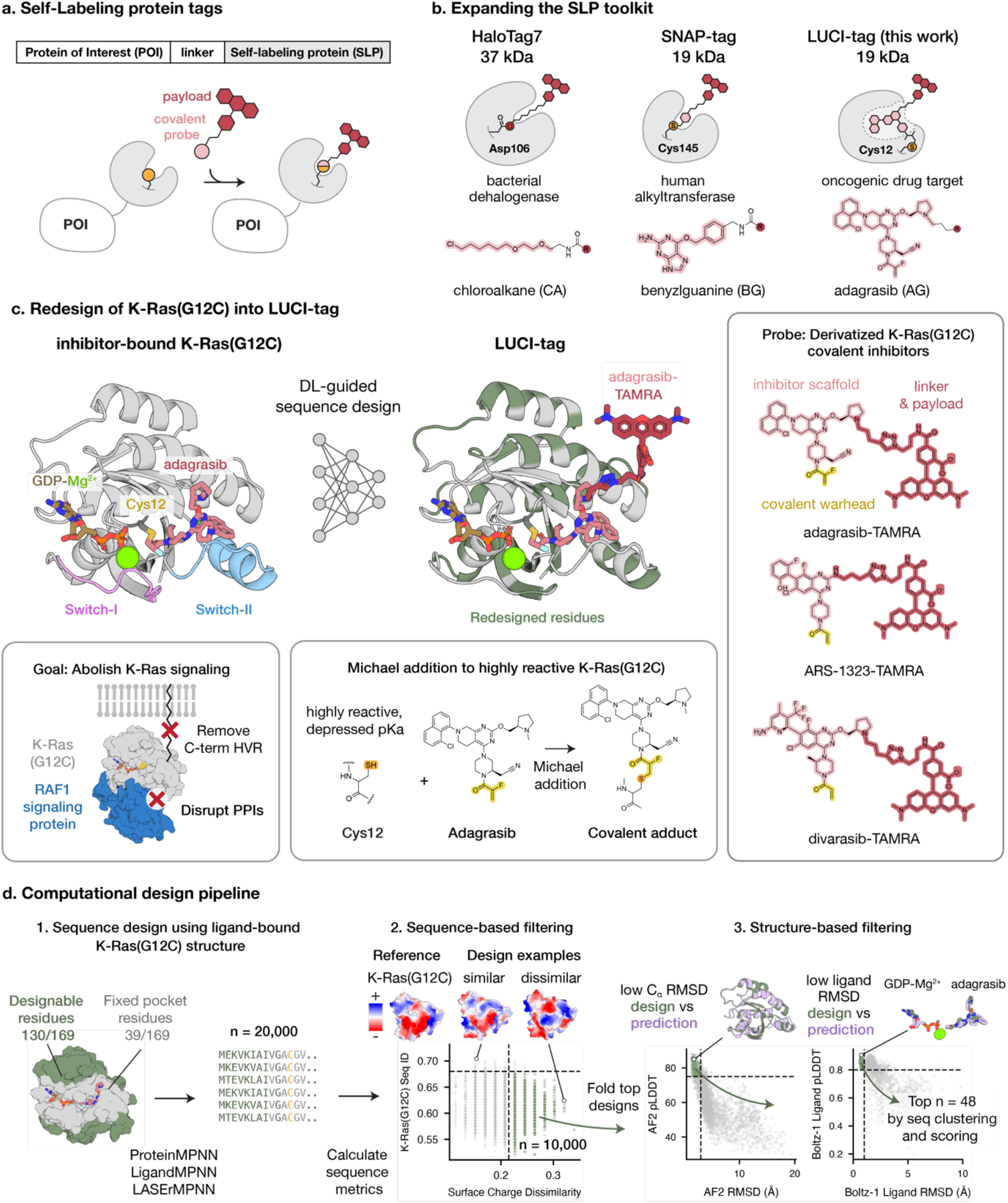
Using deep learning to convert K-Ras(G12C) into an orthogonal SLP. **a**, Schematic of a SLP tag. **b**, Two popular SLPs with their covalent probes, along with our newly designed LUCI-tag SLP. LUCI-tag targets a cryptic pocket in K-Ras(G12C) and a synthetic covalent inhibitor, whereas HaloTag7 and SNAP-tag repurpose enzyme orthosteric sites and their substrates. **c**, Unlike HaloTag7 and SNAP-tag, LUCI-tag is an extensive overhaul of its parent sequence, here K-Ras(G12C). The design goal, using structure-based sequence design, is to preserve (or enhance) the inhibitor binding of K-Ras(G12C) but eliminate all biological function of the enzyme, including protein– protein interactions (PPIs) with evolved binding partners. Covalent inhibitor binding is accelerated by the inherent reactivity of Cys12 and a med-chem optimized suite of highly specific K-Ras(G12C) covalent inhibitors, each of which has an appropriately positioned exit vector for attaching various payloads. **d**, Deep-learning-based design pipeline for LUCI-tag starting from an inhibitor-bound structure of K-Ras(G12C). Out of a total of 169 residues, 130 are designable (green). GDP-Mg^2+^ cofactor and inhibitor-contacting residues (gray) are held constant.

We wondered if a new orthogonal SLP could be developed using previously established allosteric covalent inhibitors against a human drug target. We aimed to take advantage of previous medicinal chemistry efforts to increase bioavailability, target specificity, reactivity, and cell permeability, while achieving orthogonality to HaloTag and SNAP-tag. At the same time, an ideal SLP protein should be small, monomeric, and biologically inert, with a solvent-accessible binding pocket for a probe that can accommodate bulky payloads. Considering these properties, we identified K-Ras(G12C) and its covalent inhibitors as an ideal starting point for developing a new SLP platform (Fig. 1c).

K-Ras(G12C) is a 21 kDa mutant oncoprotein that drives aberrant cell growth through membrane-localized GTP-dependent signaling^11^. The K-Ras protein structure is a compact α/β fold with flexible loops called switch I and switch II that undergo nucleotide-dependent conformational change and participate in GTP hydrolysis and effector interactions (Fig. 1c, left). Its membrane localization is driven by farnesylation of the basic C-terminal hypervariable region (HVR). Mutant K-Ras has been the target of several recent breakthroughs in precision oncology^12^, including the development of covalent inhibitors that irreversibly label the Cys12 mutation^13,14^. Unlike HaloTag and SNAP-tag ligands, which target enzyme orthosteric sites using substrate mimics, these synthetic inhibitors target the cryptic switch-II pocket, wherein their mildly electrophilic acrylamide warheads undergo Michael addition to the nucleophilic Cys12 (Fig. 1c, bottom)^15^. The combination of cryptic-pocket targeting and mutant-specific covalency affords K-Ras(G12C) inhibitors greater proteome-wide selectivity compared to other covalent drugs and forms a strong basis for SLP probe biorthogonality. In addition to their selectivity, X-ray crystal structures of drug-bound K-Ras(G12C) show the inhibitors have suitable exit vectors to attach payloads^13,14,16–18^. Lastly, the inherent reactivity of Cys12 allows inhibitors to achieve second-order rate constants within the same range as existing SLP systems (Fig. S2 and Table S1)^19^.

While K-Ras(G12C) contains many suitable properties for an SLP, a major hurdle is its biological role as part of a cell-signaling network. To scrub its interactions with binding partners, we aimed to radically redesign its sequence using deep-learning-guided protein design, while maintaining the key features for covalent drug recognition (Fig. 1, c and d).

## Design strategy

We started from a crystal structure of K-Ras(G12C) with an inhibitor bound to the switch-II pocket as input to several graph neural networks for structure-based sequence design^20–22^ (Fig. 1d and Table S2). Our primary goal was to design sequences that stably fold to the structure of K-Ras(G12C) and maintain binding of the covalent inhibitor while changing residues involved in protein–protein interactions. First, we truncated the C-terminal HVR that anchors the protein to the membrane, shortening the K-Ras scaffold to 169 residues (19 kDa). We expected removal from the membrane would hinder the protein’s interactions with membrane-associated binding partners that drive cell signaling and proliferation. We held constant the residues of the switch-II pocket and the GDP-Mg^2+^ binding site (heavy atoms within 4.8 Å of adagrasib or GDP-Mg^2+^) and allowed all other residues (surface and core, 80% total) to be designable. We disallowed use of Cys residues during design, ensuring that the only Cys in our designs would be Cys12.

We filtered designed sequences based on surface-charge dissimilarity with native K-Ras(G12C), then used AlphaFold2^23^ and Boltz-1^24^ to filter for structural self-consistency (low C*a* root mean square deviation, RMSD, between intended and predicted structures) and ligand confidence (pLDDT), selecting 48 of the 20,000 total designs for experimental interrogation (Figs. 1d and S4, Table S3). In contrast to HaloTag7 and SNAP-tag, which have 91% and 89% sequence identity to their native proteins, respectively, our designs had sequence identities ranging from 43–65% relative to K-Ras(G12C), a much more extensive overhaul of the native sequence (Fig. S1). This radical redesign of K-Ras(G12C) is now possible due to the dramatic improvements in performance of protein-design methods using deep learning.

We ordered synthetic DNA for the 48 designs and expressed and purified the proteins from *E. coli*. All designs expressed with good yields (>100 mg/L), were monomeric by size-exclusion chromatography (Fig. S5), and most were highly thermostable (T_m_ > 95 °C, Fig. S6) – much more so than a minimally engineered version of K-Ras(G12C), Cys-light ΔHVR^19^ (T_m_ = 60 °C), which we refer to here as K-Ras(G12C). For each design, we characterized the kinetics of covalent binding of the switch-II inhibitor, ARS-1323, using fluorescence polarization spectroscopy. Using a fluorophore-conjugated ARS-1323 derivative, we measured the change in fluorescence anisotropy of the dye as a function of time and protein concentration. We observed rapid labeling for half of the designs, with top designs achieving 2^nd^-order rate constants (*k*_app_) that were 10–100-fold faster than K-Ras(G12C) (Fig. S7). We chose the design with the fastest labeling kinetics and cleanest proteomics data (see below) as our lead design, which we call LUCI-tag (Labeling Using Covalent Inhibitors).

LUCI-tag has 54% sequence identity to K-Ras(G12C) while preserving the switch-II pocket and GDP-Mg^2+^ binding site (Fig. 2a, S8). In addition, LUCI-tag has drastically altered surface geometry and electrostatics compared to K-Ras(G12C), particularly at regions involved in protein– protein interfaces (Fig. 2b).

**Figure 2.**
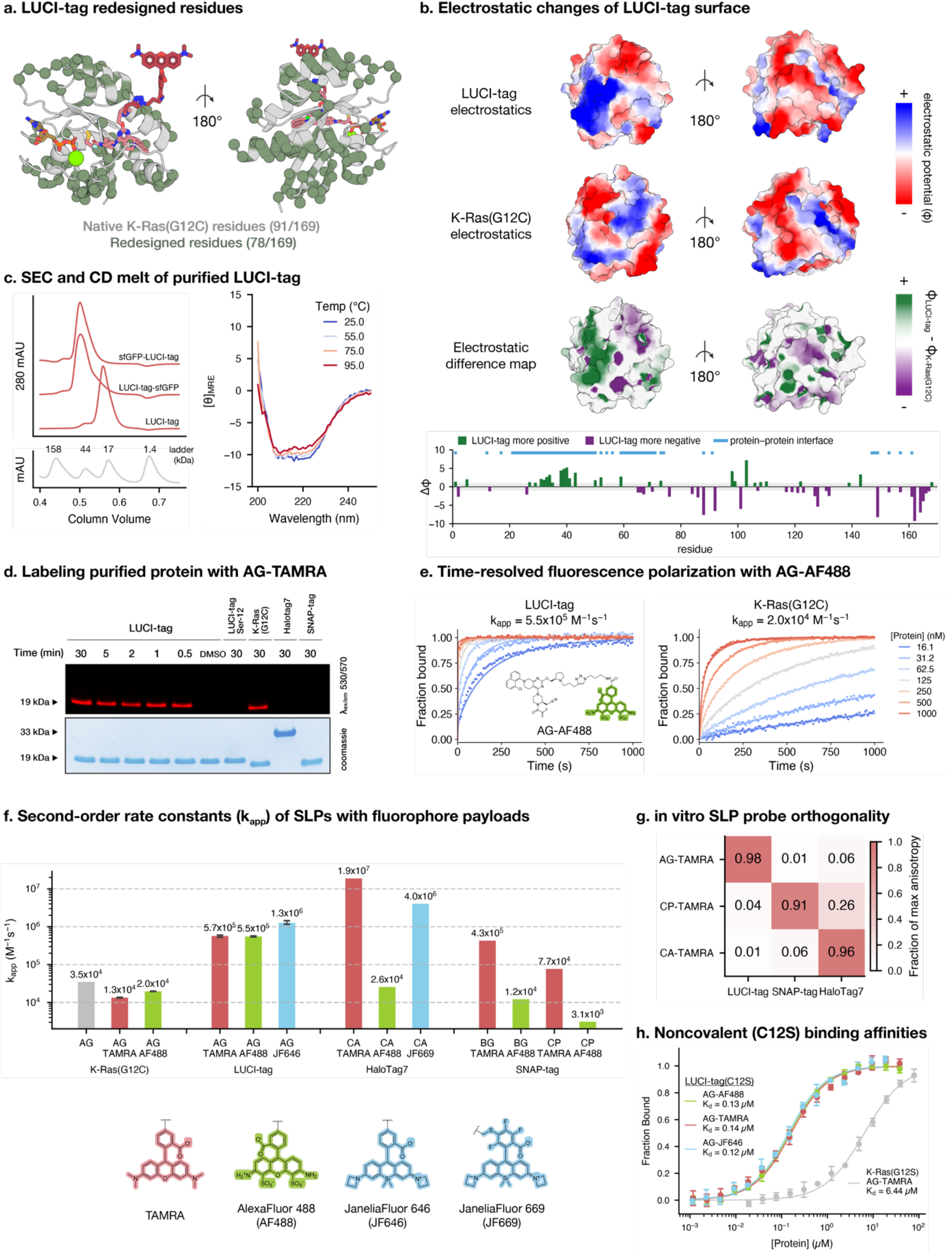
Characterization of LUCI-tag. **a,** LUCI-tag contains 78 total amino-acid substitutions relative to K-Ras(G12C), such that 46% of the K-Ras(G12C) sequence was altered in LUCI-tag (green spheres). **b**, The surface electrostatic potential of LUCI-tag is dramatically altered compared with that of K-Ras(G12C), including in known regions involved in protein–protein interactions (determined from co-crystal structures of K-Ras(G12C) with bound binding partners 6OB2, 6XI7, 1NVW, and 1HE8). **c**, LUCI-tag is monomeric by size-exclusion chromatography (left), both as a stand-alone protein and as a sfGFP fusion. Circular dichroism spectra of drug-free LUCI-tag show the protein is highly thermostable and does not melt at 95 °C. **d**, Polyacrylamide gel electrophoresis of purified LUCI-tag and other proteins (10 µM) showing covalent labeling of LUCI-tag with adagrasib-TAMRA (AG-TAMRA, 0.5 µM) after quenching at various timepoints. **e**, Time-resolved fluorescence polarization of AG-Alexa Fluor 488 (AG-AF488, 10 nM) and various concentrations of purified LUCI-tag or K-Ras(G12C). Lines denote global fits across each protein concentration to a two-step kinetic model to determine the second-order rate constant of labeling (k_app_). **f**, k_app_ for K-Ras(G12C), LUCI-tag, HaloTag7, and SNAP-tag with their respective covalent probes bearing various dye payloads; Halo- and SNAP-tag values are taken from ref. 25. **g**, All-by-all comparison of probe reactivity across LUCI-tag, SNAP-tag, and HaloTag7 and their respective TAMRA-bearing probes. [Protein] = 400 nM; [probe] = 50 nM; incubation time = 10 min. **h**, Non-covalent equilibrium binding affinities of LUCI-tag(C12S) for AG bearing different dyes and of K-Ras(G12S) with AG-TAMRA. [probe] = 50 nM. Error bars show standard deviation of 3 technical replicates.

## LUCI-tag biochemical characterization

LUCI-tag expresses well in *E. coli,* is monomeric by SEC as a standalone or fusion protein, and is stably folded at temperatures up to 95 °C (Fig. 2c). Intact mass-spectrometry showed complete covalent labeling with ARS-1323 and adagrasib (AG), with the latter showing markedly faster kinetics (Figs. S7 and S16). To generate the fluorescent LUCI-tag probe, AG-TAMRA, we attached a TAMRA dye via click chemistry to a commercially available clickable derivative of AG (Figs. S10-14). In-gel labeling of LUCI-tag with AG-TAMRA is complete within 30 s (Fig. 2d), in stark contrast to labeling of K-Ras(G12C), which requires >30 min under identical conditions (Fig. S15). In step with these results, time-resolved FP with AG-AlexaFluor488 (AF488) showed LUCI-tag possesses a fast *k*_app_ of 5.5×10^5^ M^-1^s^-1^, which is a 27-fold increase over the *k*_app_ of K-Ras(G12C) (Fig. 2e). With respect to their TAMRA-bearing cognate ligands, LUCI-tag shows similar labeling kinetics to SNAP-tag^25^ but is slower than HaloTag7, owing to HaloTag7’s extensive optimization toward rhodamine-dye payloads (Fig. 2f, Table S4). As expected, all-by-all labeling experiments showed that LUCI-tag and AG-TAMRA are orthogonal to HaloTag7 and SNAP-tag (Figs. 2g and S23).

We measured the non-covalent affinity of LUCI-tag with its inhibitor-based probes via expression of the protein with a C12S substitution, which eliminates covalent attachment of the inhibitors (Fig. 2h). LUCI-tag(C12S) binds AG-TAMRA with a dissociation constant (*K*_d_) of 0.14 µM, ∼46- fold tighter than K-Ras(G12S) (Fig. S18). Competition experiments with unlabeled AG showed 3.8-fold tighter binding to LUCI-tag than the fluorophore-tagged version (*K*_d_ = 0.03 µM, Fig. S21), indicating that the current linker may be a candidate for future optimization. The high non-covalent affinity of LUCI-tag with its probe contrasts with that of catalytically inactive mutants of HaloTag and SNAP-tag, which bind their payload-free probes with affinities in the high µM to mM range (Fig. S19, Table S5^25^).

We reasoned the tight non-covalent binding of AG could drive payload-agnostic labeling kinetics of LUCI-tag. Indeed, LUCI-tag reacts with AG-AF488, AG-TAMRA, and AG-JaneliaFluor646^26^ (AG-JF646) with *k*_app_ values all within a 2-fold range (Fig. 2f). The non-rhodamine-based cyanine dye, AF647, also labels LUCI-tag with a similarly fast rate when conjugated to AG (Fig. S16). This payload-agnostic labeling contrasts starkly with HaloTag7, where probes bearing negatively charged dyes, such as AF488, label 1000-fold more slowly than probes with uncharged rhodamines^25^. To our knowledge, LUCI-tag is the fastest reported SLP for labeling with AF488.

Fluorogenicity, i.e. turn-on post-labeling fluorescence, of dye payloads with LUCI-tag could be advantageous in no-wash microscopy protocols. Time-resolved fluorescence intensity experiments demonstrated modest fluorogenicity of AG-JF646, resulting in a 9-fold increased brightness of the JF646 dye over 90 min compared to free AG-JF646 (Fig. S33). Thus, LUCI-tag exhibits modest fluorogenicity on par with SNAP-tag, without explicit engineering for such.

## Role of GDP-Mg^2+^ cofactor in LUCI-tag

Because covalent inhibitors preferentially bind the GDP-bound state of K-Ras(G12C) over the active GTP-bound state, we sought to understand the role of nucleotide state and GDP-Mg^2+^ cofactor availability on LUCI-tag activity. We performed nucleotide exchange to uniformly load LUCI-tag with either GDP or GTP. Both preparations showed robust labeling of LUCI-tag with AG-AF488 (Fig. S24). The ability of LUCI-tag to label with AG in the GTP-bound state contrasts with that of K-Ras(G12C), which does not typically bind switch-II inhibitors when loaded with GTP^27^.

During protein expression, LUCI-tag efficiently acquires and retains cofactors without additional supplementation because GDP, GTP, and Mg²⁺ are abundant intracellular metabolites across bacteria, yeast, and mammalian cells^28^. Still, to enable the use of LUCI-tag in nucleotide- or metal-depleted environments, we designed a cofactor-free version by removing GDP-Mg^2+^ from the input structure and allowing residues within the cofactor-binding pocket to be designable (Fig. S3). Expression and testing of the resulting sequences identified LUCI-tag-CF (cofactor-free) as an SLP whose labeling is not dependent on GDP-Mg^2+^, albeit with slightly slower labeling kinetics than LUCI-tag (Figs. S16 and S25).

## X-ray crystal structures of LUCI-tag

We determined an X-ray crystal structure of AG-bound LUCI-tag (2.9 Å resolution), which showed excellent agreement with the native K-Ras(G12C) structure (C*a* RMSD of 1.1 Å), including specific interactions with the inhibitor in the switch-II pocket and evidence of covalent bond formation (Fig. 3a,b). An additional structure of LUCI-tag bound to the inhibitor ARS-1323 shows its switch II pocket can faithfully recapitulate K-Ras(G12C) interactions for other inhibitors (Fig. S45). We additionally determined the structure of drug-free (GDP-Mg^2+^ bound) LUCI-tag (2.1 Å resolution), which contained a well-resolved switch-II loop that forms an open inhibitor-binding pocket, perhaps preorganizing LUCI-tag for probe binding (Figs. 3c and S26). In K-Ras(G12C), the flexible switch-II loop samples different conformations that alter the volume of the cryptic pocket. Indeed, most drug-free structures of K-Ras and K-Ras(G12C) in the PDB show no cryptic pocket at all^29^, in stark contrast to LUCI-tag (Fig. 3d).

**Figure 3.**
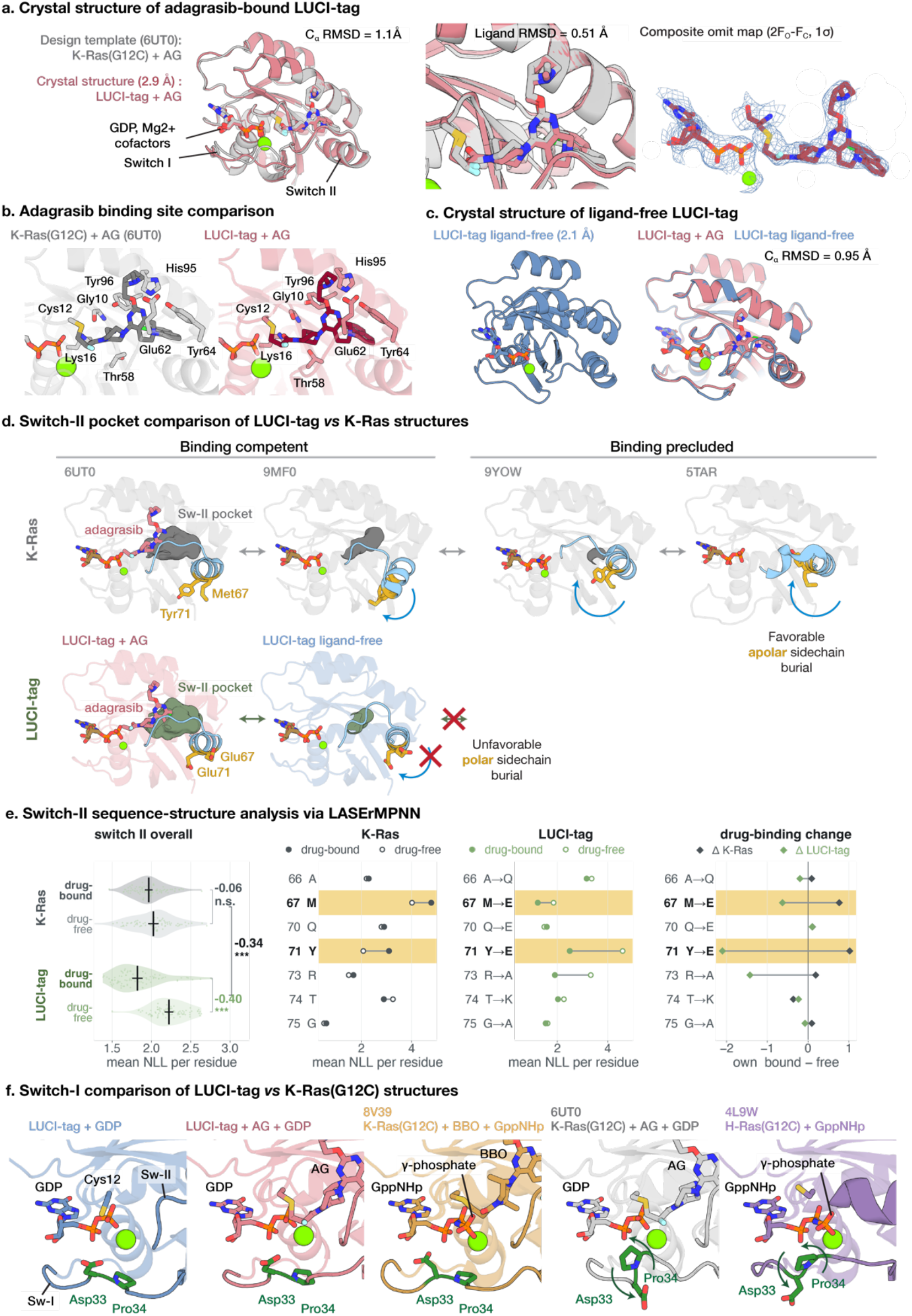
X-ray crystal structures of drug-bound and drug-free LUCI-tag. **a,** Crystal structure (2.9 Å resolution) of AG-bound LUCI-tag (red) superimposed with crystal structure of AG-bound K-Ras(G12C) (gray, PDB 6UT0), showing low C*a* root mean square deviation (RMSD) and low AG heavy-atom RMSD. Both structures contain the GDP-Mg^2+^ cofactor. The composite omit map shows strong density for the cofactor, AG, and the Cys12 covalent bond. **b**, Close-up of the crystal structures showing that the switch-II binding pocket of LUCI-tag is nearly identical to that of K-Ras(G12C). **c**, The drug-free (GDP-Mg^2+^-bound) structure of LUCI-tag (2.1 Å resolution) is highly similar to the structure of AG-bound LUCI-tag, with a resolved switch-II loop. **d**, Samples of structures in the PDB of K-Ras showing the range of switch-II (Sw-II) pocket sizes. In drug-free structures (right), the switch-II loop rotates inward and collapses the cryptic pocket. PDB accession codes are shown above each structure in gray. **e**, LASErMPNN analysis of the switch-II sequence of K-Ras(G12C) (gray) and LUCI-tag (green) applied to 170 crystal structures of drug-free and drug-bound K-Ras and K-Ras(G12C) from the PDB. The mean negative log likelihood (NLL) over the switch-II residues is shown, with each point a different structure in the PDB. K-Ras(G12C) and LUCI-tag sequences are evaluated over the same structures. Individual amino-acid contributions to the NLL are shown (right), with predominantly contributing residues highlighted in yellow. Each point in the rightmost plot shows the difference (bound – free) between the points in each protein’s plot on the left. E67 and E71 of LUCI-tag show the largest gap between drug-bound and drug-free structures, with overwhelming preference (lower NLL) for drug-bound structures. **f**, Close-up of the switch-I region (bottom) of LUCI-tag and K-Ras(G12C) in various nucleotide- and inhibitor-bound states. The switch-I loop of both drug-free and AG-bound (GDP-Mg^2+^-bound) LUCI-tag closely resembles that of a “dual-state” inhibitor-bound (GppNHp-Mg^2+^-bound) structure of K-Ras(G12C), each of which contrasts with the structure of the switch-I loop in AG-bound (GDP-Mg^2+^-bound) and drug-free (GppNHp-Mg^2+^-bound) K-Ras(G12C). The open structure of the switch-I loop of LUCI-tag is consistent with switch-II inhibitors binding in both the GDP- and GTP-bound states.

Two key amino-acid substitutions show that the sequence of LUCI-tag is biased toward a pocket-open structure. In K-Ras(G12C), the apolar switch-II residues, Met67 and Tyr71, are solvent exposed in drug-bound structures, whereas these residues rotate inward and pack within the protein core in drug-free structures, closing the cryptic pocket (Fig. 3d, yellow). Both Met67 and Tyr71 were mutated to negatively charged glutamates in LUCI-tag, which should energetically destabilize a pocket-closed structure that would require their burial. To quantify this effect, we used LASErMPNN to analyze sequence–structure compatibility across 170 crystal structures of drug-bound and drug-free K-Ras and K-Ras(G12C) in the PDB (Fig. 3e). We threaded K-Ras(G12C) and LUCI-tag sequences onto each structure and calculated the per-residue negative log likelihood (NLL), where a lower NLL implies greater sequence–structure compatibility at that position. The K-Ras switch-II sequence had similar NLL distributions for both the drug-bound and drug-free structures, indicating the native sequence is similarly compatible with both ensembles, as expected for the evolved protein function. By contrast, the NLL of LUCI-tag’s switch-II sequence—which differed from native by only 7 solvent-exposed residues—showed that LUCI-tag strongly favors a drug-bound structure over a drug-free structure. This effect is primarily driven (>75%) by just two substitutions, M67E and Y71E (Fig. 3e, right).

In addition to switch II, amino-acid substitutions in LUCI-tag’s switch-I loop may also contribute to faster labeling kinetics. Relative to K-Ras(G12C), the switch-I loop in both drug-free and drug-bound LUCI-tag is further removed from its switch-II loop, closely resembling a drug-bound structure of K-Ras(G12C) loaded with GTP (Fig. 3f). A few key mutations might account for the alternate structure of the switch-I loop in LUCI-tag. Thr35, a functionally important residue that dictates K-Ras signaling via coordination of the GTP γ phosphate^30^, was changed to Asn in LUCI-tag. This substitution ablates the γ-phosphate coordination and promotes an outward-facing, signaling-incompetent orientation of this residue in switch I. Pro34 is also tucked inward, pointing away from the nucleotide pocket, a conformation observed in “dual state” inhibitors that bind K-Ras(G12C) in the GTP-bound state (e.g., PDB 8V39). The similar switch-I loop geometry to dual-state inhibitor-bound K-Ras(G12C) corroborates our experimental observations that probe binding can occur in both the GTP- and GDP-bound forms of LUCI-tag. Taken together, the structures of drug-bound and drug-free LUCI-tag suggest that increased sequence–structure compatibility via deep-learning-based sequence design led to structural preorganization of the switch-I and switch-II regions, resulting in enhanced labeling kinetics and dual GDP/GTP labeling.

## Bio-orthogonality and functional abrogation

The surface of LUCI-tag was dramatically altered relative to K-Ras(G12C) (Fig. 2b), including charge reversals, shifts in polarity, and changes in sterics (Fig. S28). These changes should eliminate interactions with native K-Ras binding partners, such as RAF kinases and nucleotide exchange factors. To confirm the biological orthogonality of LUCI-tag, we performed mass-spectrometry-based proteomics experiments with human cell lysates (Fig. 4a, S29). We used an affinity-tagged LUCI-tag to perform immunoprecipitation-mass spectrometry (IP-MS) on lysates of Molt4 cells and compared changes in pulled-down protein abundance against those from an affinity-tagged K-Ras(G12C) control. As expected, K-Ras(G12C) pulled down its known binding partners, while LUCI-tag showed a near-complete loss of Ras-family and Ras-associated binding. Thus, LUCI-tag was effectively disengaged from its biological interaction network without forming any neointeractions relative to K-Ras(G12C).

**Figure 4.**
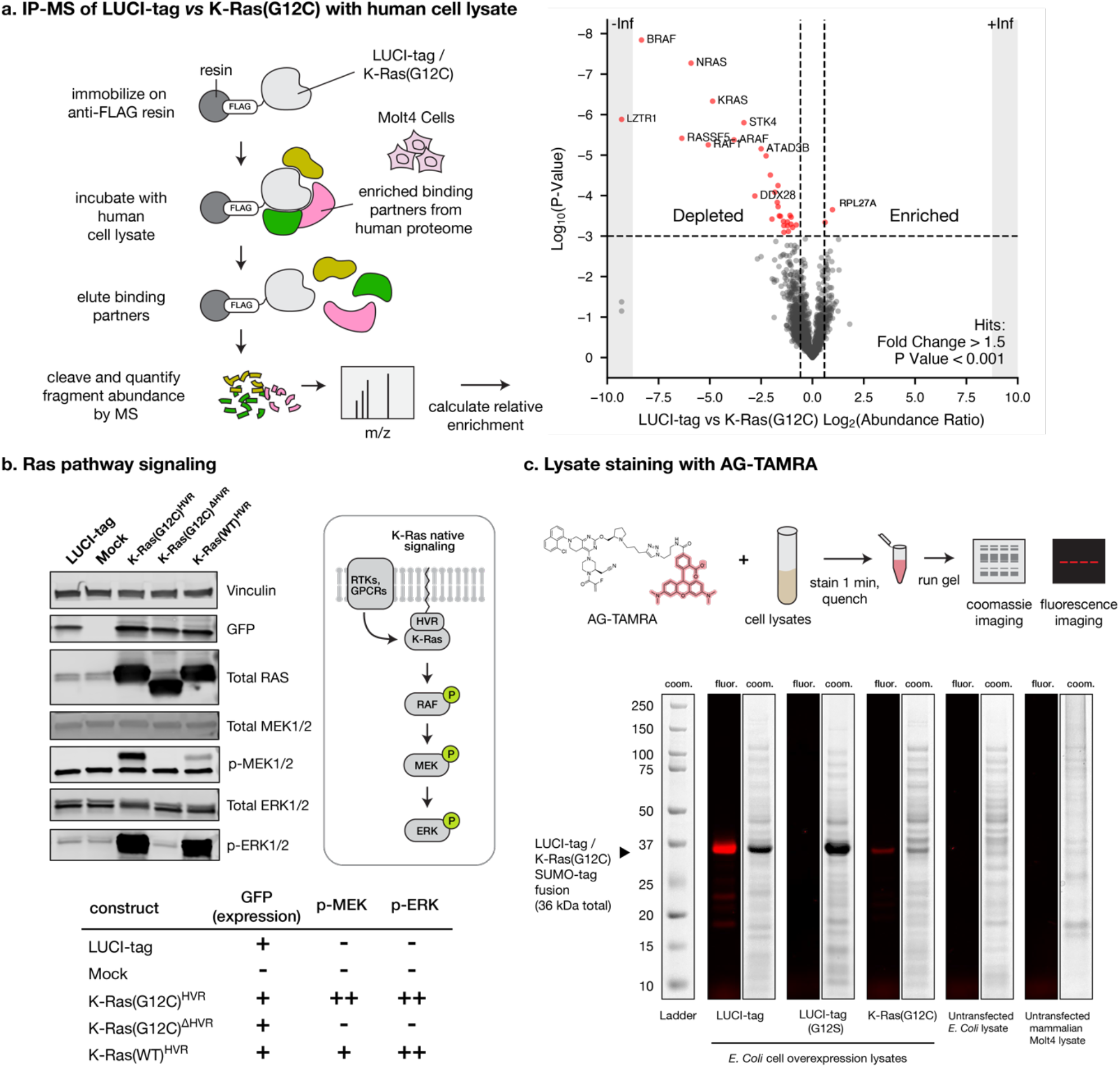
Bio-orthogonality of LUCI-tag and its probes. **a,** Immunoprecipitation–mass spectrometry (IP-MS) of endogenous proteins in Molt4 mammalian cell lysate pulled down via FLAG-tagged LUCI-tag or K-Ras(G12C). The abundance ratio of proteins pulled down via LUCI-tag *vs* K-Ras(G12C) shows the typical binding partners associated with K-Ras are depleted in the pulldown with LUCI-tag, while no neo-interactions with LUCI-tag are observed. **b,** Ras-pathway cell-signaling assay of transfected HEK293T cells quantified via western blot for protein expression and MEK/ERK phosphorylation. Mock is an empty plasmid control. GFP on the expression plasmid serves as an expression marker. The table below summarizes the findings from the assay. Note that the anti-RAS antibody used to measure total K-Ras does not bind LUCI-tag, highlighting the extensive surface redesign. HVR denotes an untruncated K-Ras, which contains the hypervariable region that confers membrane association, whereas ΔHVR denotes truncation of this region. **c,** 1-min stain of AG-TAMRA (100 nM) in *E. coli* lysates overexpressing LUCI-tag, its non-covalent variant, or K-Ras(G12C). Lysates from untransfected *E. coli* or mammalian Molt4 cells showed no off-target staining of the probe. Fluorescence channel was 550 nm. Coom. = coomassie. Proteins were expressed with an N-terminal SUMO-tag fusion, accounting for their larger size (36 kDa).

To confirm that LUCI-tag does not function in the cell-signaling pathways of K-Ras(G12C), we performed cell-signaling experiments and assessed downstream phosphorylation via western blot (Fig. 4b). While cells transfected with recombinant full-length K-Ras(G12C) showed expected phosphorylation of signaling partners, cells transfected with LUCI-tag were indistinguishable from an untransfected control. Expression of C-terminally truncated K-Ras(G12C) also showed no phosphorylation of signaling partners, in agreement with prior results that K-Ras(G12C) is only functional when anchored to the cell membrane^11^. Thus, as intended, intracellular overexpression of LUCI-tag does not elicit unwanted aberrant signaling that defines oncogenic K-Ras(G12C).

Lastly, we evaluated the specificity of LUCI-tag labeling in *E. coli* and mammalian proteomes. After incubation in cell lysates with overexpressed LUCI-tag, fluorescence imaging of the soluble fraction within SDS-PAGE gels showed that LUCI-tag is uniquely stained by AG-TAMRA (Fig. 4c). Noncovalent LUCI-tag(C12S) and untransformed lysates showed no such labeling. Taken together, these results show LUCI-tag is both chemically precise and biologically inert.

## Live-cell microscopy

We next sought to demonstrate rapid, wash-free fluorescent labeling of proteins with LUCI-tag in live-cell microscopy. We transiently transfected HEK293T cells with a construct that fused LUCI-tag to a nuclear-localized protein, PITX2, together with T2A-based expression of GFP as a transfection marker (Fig. 5a). After 24–48 h of protein expression, we stained the cells with AG-JF646 for 30 min and collected confocal fluorescence images without washing. We observed a strong, nucleus-specific signal for LUCI-tag while GFP was spread throughout the cytoplasm and nucleus, as expected. Staining was restricted to transfected, GFP-positive cells (Fig. 5a, panel 3) and remained robust at probe concentrations down to 125 nM (Fig. S30). Additional washing and incubation in cell media did not affect signal relative to background (Fig. S31). The control constructs, K-Ras(G12C) and LUCI-tag(C12S), showed no detectable labeling when similarly expressed in HEK293T cells (Figs. S41 and S42). Overall, processed image quality was comparable to the same construct using HaloTag7 instead of LUCI-tag, although raw HaloTag7 images were brighter due to it being more fluorogenic with JF646 (Figs. S32 and S34). Consistent with this, live-cell time-course imaging using JF646 payloads showed that LUCI-tag has similar turn-on fluorescence kinetics as SNAP-tag, although both are slower than HaloTag (Fig. S34).

**Figure 5.**
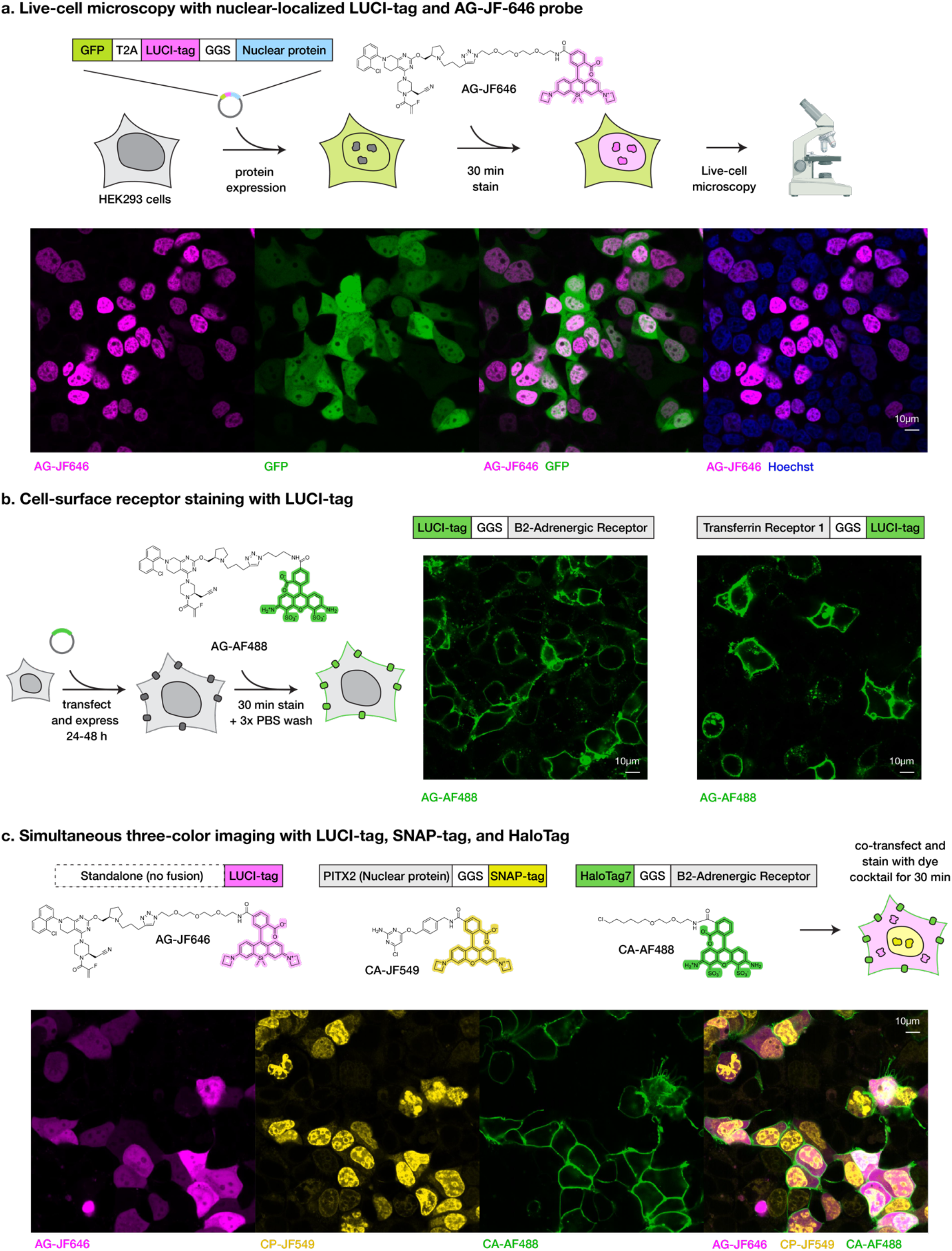
LUCI-tag enables three-color live-cell imaging with SLPs. **a,** Protocol and images for live-cell microscopy with LUCI-tag in HEK293T cells. Plasmids included a GFP expression marker and nuclear-localized LUCI-tag (1 µM AG-JF646, 30 min stain, no wash). Nuclear staining is only observed in transfected, GFP-positive cells. **b,** Cell-surface receptor staining of LUCI-tag via fusion to the extracellular terminus of membrane proteins transferrin receptor 1 (TFR1) and β2-adrenergic receptor (B2AR) (500 nM AG-AF488, 30 min stain, 3x PBS washes). Robust membrane staining is observed, with puncta seen in cells with labeled TFR1 due to the rapid internalization of the receptor. **c,** Three-color imaging with B2AR-fused HaloTag7, nuclear-localized SNAP-tag, and standalone LUCI-tag. HEK293T cells were co-stained with CA-AF488, CP-JF549, and AG-JF646 dye cocktail for 30 min followed by a 30-min wash in imaging media. Robust, orthogonal, three-color staining is observed.

We next wondered if LUCI-tag could be used for extracellular staining. We generated LUCI-tag fusions to the extracellular termini of transferrin receptor 1 and β2 adrenergic receptor and expressed these constructs in HEK293T cells, followed by staining with AG-AF488 and washing (Fig. 5b). Confocal images revealed robust extracellular labeling of the receptors, with equivalent performance by the corresponding HaloTag7 constructs with HaloTag ligand CA-AF488 (Fig. S35). Live-cell time-course imaging with AG-JF646 showed fast extracellular labeling kinetics similar to HaloTag (Fig. S36).

As expected from our biochemical experiments, LUCI-tag is orthogonal to both HaloTag7 and SNAP-tag in multicolor live-cell imaging. We performed two-color imaging by co-expression of nuclear-localized LUCI-tag with standalone SNAP-tag or HaloTag7, followed by simultaneous staining with differently colored dyes. We observed spatially exclusive labeling of the nucleus by LUCI-tag (Fig. S37). We next expressed a construct where both LUCI-tag and HaloTag7 were fused in tandem to a nuclear-localized protein (LUCI-tag-GGS-HaloTag7-PITX2), followed by staining with differently colored CA and AG probes. As expected, the tandem construct showed identical localization for both tags (Fig. S38), indicating facile two-color labeling with minimal interference between channels.

A central goal of this work was to establish LUCI-tag as an orthogonal SLP for three-color live-cell imaging with HaloTag7 and SNAP-tag. To directly test this, we co-transfected HEK293T cells with plasmids encoding standalone LUCI-tag, membrane-directed HaloTag7, and nucleus-directed SNAP-tag. We simultaneously stained with tag-specific ligands (500 nM) containing spectrally distinct dyes (JF646, JF549, and AF488). After staining for 30 min followed by a wash, we collected confocal fluorescence images, which showed spatially exclusive three-color labeling with no cross-reactivity between the tags (Fig. 5c).

## Discussion

Here, we repurposed an infamous oncoprotein, K-Ras(G12C), to create a new, biologically silent SLP for molecular biology. LUCI-tag fully labels within seconds at nanomolar concentrations and is compatible with HaloTag7 or SNAP-tag for mammalian live-cell microscopy with short, wash-free staining protocols. Its compact size (169 amino acids, 19 kDa), exceptional thermostability (T_m_ > 95 °C), and close positioning of the N- and C-termini make it an ideal fusion partner in a wide variety of contexts (Fig. S44), potentially facilitating loop insertions^8^ and enabling harsh conditions that would denature most tags.

LUCI-tag shows payload-agnostic labeling kinetics, with fast rates regardless of the charge of the dye payload. This performance contrasts with HaloTag7, whose reaction kinetics drop over 1000-fold for payloads other than the rhodamine dyes for which it was explicitly optimized^25^. Thus, LUCI-tag should be equally amenable for attaching payloads such as cyanine dyes (Fig. S16), click chemistry handles, biotin, oligonucleotides, or degradation handles, providing a distinct advantage for a variety of molecular biology protocols.

LUCI-tag is distinct from other tags in that it uses allosteric covalent inhibitors rather than a trapped enzyme intermediate (HaloTag7) or product (SNAP-tag). Relative to other synthetic covalent inhibitors, targeting the allosteric pocket of mutant K-Ras(G12C) affords hyperspecificity of LUCI-tag’s covalent probe, which shows no off-target labeling in bacterial and mammalian lysates. The unique K-Ras(G12C) switch-II pocket, mutation-specific covalency of its inhibitors, and dramatic sequence alterations by protein design allow for sufficient orthogonality to biological proteins and other SLPs. While most common cell lines such as HEK293 and U2OS do not contain the K-Ras(G12C) mutation^31^, LUCI-tag can also be used in K-Ras(G12C)-endogenous cell lines, as we found no interference from background staining in such cells under usual incubation times (Fig. S42). LUCI-tag’s faster labeling kinetics, likely facilitated by its preorganized, pocket-open structure, enables it to outcompete K-Ras(G12C) in cells.

Because LUCI-tag keeps the same switch-II binding site of K-Ras(G12C), we expect that any new covalent inhibitors developed for K-Ras(G12C) could be rapidly repurposed for LUCI-tag. For example, the next-generation inhibitor divarasib^18,32^, which has very fast labeling kinetics to K-Ras(G12C), also covalently labels LUCI-tag (Fig. S9). FP experiments showed divarasib has three-fold higher noncovalent affinity than AG with LUCI-tag(C12S) (*K*_d_ = 9.8 nM; Fig. S22). This suggests derivatives of divarasib could be developed as next-generation LUCI-tag probes with even faster labeling kinetics. Importantly, the use of clinically developed, med-chem-optimized, covalent K-Ras(G12C) inhibitors might confer good bioavailability of these reagents *in vivo*, as these molecules have been optimized to minimize plasma protein binding and off-target reactivity.

The clinical importance of K-Ras(G12C) inhibitors confers wide probe accessibility for LUCI-tag: clickable versions of some inhibitors, such as AG, are already commercially available, providing immediate and affordable access to inhibitor–payload conjugates. In this work, we synthesized LUCI-tag probes using standard click chemistry reactions from commercially available reagents. To make the technology even more accessible, we developed a “direct to biology” protocol that does not require purification of the clicked ligand–payload conjugates yet maintains similar performance in live-cell microscopy (Figs. S13, S14, and S43; Table S8). While triazole linkers generated by click chemistry might hinder cell permeability, we observed bright staining with minimal background in a 15-min, no-wash protocol. Fit-for-purpose, non-clickable AG probes with optimized linkers could be synthesized to maximize cell permeability, as has been done previously for K-Ras(G12C) PROTACs^33^.

LUCI-tag does not require the addition of exogenous cofactors in most contexts such as mammalian or bacterial expression due to the abundant availability of GDP and Mg^2+^. However, *in vitro* labeling kinetics depend on the availability of Mg^2+^ in the buffer, with EDTA addition leading to decreased kinetics. This is expected as LUCI-tag kept the same high affinity GDP-Mg^2+^ binding site as native K-Ras(G12C). Our cofactor-free design, LUCI-tag-CF, showed similarly rapid labeling kinetics but with negligible dependence on GDP and Mg²⁺ (Fig. S25). LUCI-tag-CF was biologically silent by IP-MS (Fig. S29) and functional as an SLP in live-cell mammalian microscopy (Fig. S40). Thus, while LUCI-tag is performant under a variety of staining conditions, LUCI-tag-CF can be used in nucleotide- or Mg²⁺-depleted buffers.

Our work shows that protein-design methodologies can rapidly repurpose natural proteins to create biologically orthogonal molecular tools. While our main motivation for LUCI-tag was to tap into the extensive med-chem optimization of K-Ras(G12C) inhibitors, it also portends a near future where such proteins could be created completely de novo using bespoke bioorthogonal covalent ligands^20,34,35^. Future engineering efforts could focus on probe optimization and improving the fluorogenicity of LUCI-tag with rhodamine dyes. We expect LUCI-tag to emerge as an immediately accessible and powerful SLP that can be used alongside Halo- and SNAP-tags (and their future variants), unlocking facile three-color imaging with bright, synthetic dyes.

## Methods

See supplementary information.

## Data Availability

Coordinates and data files for X-ray crystal structures of adagrasib-bound LUCI-tag, drug-free LUCI-tag, and ARS-1323-bound LUCI-tag have been deposited in the PDB with accession codes 39AS, 39AT, and 39AU, respectively. Proteomics raw data has been deposited in the PRIDE archive under accessions PXD084198, PXD084227 and PXD084242.

## Code Availability

Code for the LUCI-tag design pipeline can be found in Zenodo^36^.

## Acknowledgements

J.M. and N.F.P. thank SBGrid for support of on-site compute infrastructure and crystallography software. N.F.P. and J.M. thank John Alberta for help with equipment and resources for experiments. J.M. thanks Jeremy Quintana for sharing ARS-1323–Cy5 probe for initial testing.

N.F.P. and J.M. thank AI Proteins for sharing sumo-tag and sumo-protease expression plasmids for protein expression and purification, respectively. This research used resource 17-ID-1 of the National Synchrotron Light Source II, a U.S. Department of Energy (DOE) Office of Science User Facility operated for the DOE Office of Science by Brookhaven National Laboratory under Contract No. DE-SC0012704.

## Author contributions

J.M. performed the design computations, protein biochemical and structural characterization, and microscopy. S.G. and Q.K. performed probe synthesis. J.W.L. performed cloning and transfections for microscopy. M.M. performed transfections and Western blots. K.A.D. performed immunoprecipitation-mass spectrometry. R.M. performed intact mass spectrometry. F.L.S. and K.S. performed crystallography. D.L. and D.X.C., contributed to microscopy methodology and A.D. contributed to computational methodology. K.M.H., E.S.F., F.C., and N.F.P. contributed lab resources. J.M. and N.F.P. performed analysis and wrote the paper.

## CRediT author statement

J.M.: Conceptualization, Methodology, Investigation, Writing J.W.L.: Investigation

S.G.: Methodology, Investigation

M.M., K.D., R.M., K.S., F.L.S., Q.K.: Investigation

D.L., D.X.C., A.D.: Methodology

K.M.H., E.S.F., F.C.: Resources

N.F.P.: Conceptualization, Writing, Resources, Supervision

## Additional information

Correspondence and requests for materials should be addressed to Nicholas Polizzi.

## Computational Methods

### Structural templates and preprocessing

Crystal structures of K-Ras(G12C) covalently bound to switch-II inhibitors were obtained from the Protein Data Bank (PDB), including K-Ras(G12C) bound to BI-0474 (PDB: 8AFB), sotorasib (PDB: 6OIM), and adagrasib (AG) (PDB: 6USZ). These structures were chosen for their relatively high resolution and the commercial availability of inhibitors. Structures were processed by removing waters and non-relevant heteroatoms and standardizing residue numbering. We kept the modeled switch-II inhibitors in each structure. GDP and Mg²⁺ were retained or removed depending on the design category (see below). RFdiffusion-2^1^ was used to fill in the backbone of unresolved residues in 6OIM (residues 105–107) and 8AFB (residues 166–169). The C-terminal membrane-associated hypervariable region (HVR) was absent from each of the structural templates and was not used in design.

### Designable residue definition and PPI disruption strategy

We refer to a “designable” residue (as opposed to a fixed residue) as one that can be mutated by sequence-design methods. Our goal in defining fixed *vs* designable residues was to maintain fast inhibitor-binding kinetics while disrupting the interaction of K-Ras(G12C) with downstream signaling partners. We defined the following sets of residues as (i) inhibitor binding-site residues: within 4.8 Å heavy-atom distance of inhibitor, (ii) cofactor binding-site residues: within 4.8 Å heavy-atom distance of cofactors GDP or Mg²⁺, and (iii) Protein–protein interaction (PPI) residues: those involved in known K-Ras PPIs. In all designs, Cys12 and Lys16 were fixed, as Cys12 is the site of covalent-inhibitor attachment and Lys16 is suggested to contribute to cysteine reactivity^2^. Five sequence-design categories were created with varying levels of designable residues, ranging from more conservative fixed-pocket design to more radical design allowing all residues (except Cys12 and Lys16) to be designable, (Supp. Fig. S3):

Category 0: Inhibitor- & cofactor-binding sites are fixed

Category 1: Inhibitor- & cofactor-binding sites are fixed except for key K-Ras PPI residues Category 2: Inhibitor binding site fixed; GDP-Mg^2+^ cofactor removed

Category 3: GDP-Mg^2+^ cofactor binding site fixed; inhibitor binding site is designable Category 4: Only Cys12 and Lys16 are fixed; GDP-Mg^2+^ cofactor removed

In design category 1, PPI residues were determined from structures with binding partners^3^. In particular, several effector-involved Switch-I/II residues originally classified as part of the inhibitor binding site were made designable due to their outward orientation. Categories 2 and 4 explicitly removed the GDP-Mg^2+^ cofactor with the goal of creating designs that did not depend on GDP-Mg²⁺, while categories 3 and 4 redesigned the inhibitor binding site to explore the possibility of improving inhibitor binding.

### Sequence design strategies

Sequence redesign was performed using ProteinMPNN^4^, LigandMPNN^5^, and LASErMPNN^6^, with separate design runs for each structural template and design category. We generated 512 sequences per model/structural template/design category combination (see Table S2 for model weights and parameters). ProteinMPNN, which does not recognize ligands, was only used for design categories 0 and 1.

### Structure prediction

Designed sequences were evaluated using two complementary structure-prediction approaches. ColabFold^7^ (AlphaFold2^8^, AF2) was used in single-sequence mode (no multiple sequence alignment) without ligands to compute predicted LDDT (pLDDT), predicted aligned error (PAE), and backbone C⍺ RMSD relative to starting design templates (e.g., PDB structures). Because AF2 struggled with the flexible loops of K-Ras(G12C) in single-sequence mode, we used “core pLDDT” and “core RMSD” metrics consisting of core secondary structure (residues 1-23,40-55,70-114,127-169), omitting most of the Switch-I/II loops. While the pre-organization of switch II is important to binding, we separately assessed this using binding-site RMSD, described below. To bias toward monomeric designs, ColabFold-multimer was used to fold two copies of each design, and designs were chosen that had high interfacial PAE between contacting residues across the predicted homodimer interface.

To model the three-dimensional interactions between the inhibitor and designed proteins, we used the ligand-aware co-structure predictor, Boltz-1^9^. Boltz-1 was run with 5 recycles and 5 output models per designed sequence. The covalent bond between the inhibitor and Cys12 was specified using explicit atom–atom constraints between the Cys12 sulfur and the electrophilic carbon of the inhibitor. We computed ligand heavy-atom RMSD after superposing the predicted protein–ligand co-structure onto the input PDB model via the C⍺ atoms of inhibitor binding-site residues. We also extracted ligand pLDDT from the predicted models for downstream filtering. Cofactors GDP-Mg²⁺ were included in the prediction when part of the design category. Each structural template was paired with its corresponding inhibitor for prediction, using its CCD code.

In addition to ligand-centric calculations, we also looked at the backbone heavy-atom RMSD of the binding-site residues (residues within 4.8 Å heavy-atom distance of ligands or cofactors). This allowed us to ensure residues around the ligand were correctly oriented. To calculate this binding-site RMSD, predicted structures were aligned to their corresponding templates using the N, Ca, C, O atoms of binding-site residues to avoid alignment bias from distal regions, and the RMSD of these backbone atoms was computed. To assess binding-site preorganization, we computed binding-site RMSD using ligand-bound (Boltz-1) and ligand-free (ColabFold) structures.

### Filtering criteria

We filtered designs using a multi-stage pipeline. First, we applied a sequence-level filter to designs. We removed any sequences with poly-alanine repeats, excessive charge, or high sequence identity to native K-Ras. To promote disruption of native PPIs, we selected designs with different surface-charge patterning relative to native K-Ras, defining surface residues as those with > 30 Å^2^ solvent-accessible surface area (using freesasa). The charge similarity between K-Ras(G12C) and designed sequences was calculated across all surface residues (assigned -1 for D, E, +1 for K, R, and 0 for all others). We favored designs with low charge similarity relative to the charged surface residues of K-Ras(G12C) that are involved in PPIs. We reasoned this would bias designs to disengage from the biological pathway of K-Ras.

We constrained the overall charge of our designs to between −9 and +7 [K-Ras(G12C) has a charge of -7.]. Sequences were also filtered for diversity, with different sequence-identity cutoffs used for the different design categories, described below. Generally, designs did not exceed 60–70% sequence identity to native K-Ras(G12C).

### Filtering pipeline

ProteinMPNN-, LigandMPNN-, and LASErMPNN-based designs were first filtered by the metrics and cutoffs listed in Table S3. The designs that passed these filters were clustered by sequence identity and ranked by multiple criteria (see below) to choose the top 48 for experimental characterization. We processed designs from ProteinMPNN and LigandMPNN separately from designs from LASErMPNN, since these models required different filtering thresholds.

We used three ranking metrics derived from structure prediction: binding-site RMSD from Boltz-1, mean ligand pLDDT (Boltz-1), and the protein-core pLDDT (ColabFold). Protein-core pLDDT was defined as secondary structure elements of K-Ras (127 residues total, including 1–23, 40–55, 70–114, 127–169), omitting loops that were often poorly predicted using single-sequence mode (no multiple-sequence alignment). A fourth metric, charged PPI residue similarity, measures the extent a design keeps the identities of residues involved in native PPIs. We considered 53 charged residues in K-Ras(G12C) at interfacial positions, and sought to minimize the number of residues in designs that contain the same charge as the native protein.

We combined the individual metrics into a single ranking score by normalizing each metric and taking a weighted sum. We used a weight of 0.4 for binding-site RMSD, 0.3 for mean ligand pLDDT, and 0.3 for charged PPI residue similarity. We then clustered the designs from ProteinMPNN and LigandMPNN at a sequence-identity threshold of 80%, and we kept the best-scoring design from each cluster.

We next partitioned these filtered designs into sub-groups for each of the five design categories and the three input design templates (15 sub-groups total). Within each sub-group, we clustered the designs at a sequence-identity threshold between 0.70 and 0.80 and kept the best-scoring design from each cluster, then selected the top-ranked designs by score. The score for this step weighted four metrics equally: binding-site RMSD, mean ligand pLDDT, charged PPI residue similarity, and the protein-core pLDDT. We chose four designs from each of the three template sub-groups for design-categories 0 and 1 (24 total designs) and one design from each template sub-group for design-categories 2, 3, and 4 (9 total designs), which yielded 33 designs. Because the groups were defined by category and template rather than by design method, designs from ProteinMPNN and LigandMPNN competed directly within each group, and the resulting split between the two methods was an outcome of the ranking rather than an explicit quota.

Separately, we applied the same sub-grouping to the designs from LASErMPNN. Within each of the fifteen sub-groups, we clustered at a sequence-identity threshold between 0.70 and 0.80 and selected the single top-ranked design. The score for this step weighted five metrics equally: the four metrics listed above along with protein-core RMSD (ColabFold). This resulted in 15 selected designs.

Concatenating the two branches (ProteinMPNN/LigandMPNN vs LASErMPNN) gave a final set of 48 designs, comprising 22 designs from ProteinMPNN, 11 designs from LigandMPNN, and 15 designs from LASErMPNN. Chosen designs were evenly distributed across the three structural templates, with 16 designs from each.

### Biochemistry Methods

#### Cloning, protein expression, and protein purification

Genes of the chosen 48 designs were codon optimized for *E. coli,* and synthetic DNA was ordered from Integrated DNA Technologies (IDT), followed by cloning into a peAIP32 plasmid using a Golden Gate reaction (cloning site BsaI). The construct included a 6x His tag followed by a Cth Sumo protease cleavage sequence (Sumo tag) and the protein of interest. The Golden Gate reaction was added directly into competent BL21(DE3) *E. coli* prepared with the Zymo Research Mix-n’-Go kit. Cells were then plated onto LB/kanamycin agar plates and single colonies were selected for whole-plasmid nanopore sequencing (Quintara Biosciences). Once cells were verified to contain the correct plasmid, glycerol stocks were created and used for subsequent growths.

The buffer used for all biochemistry experiments (“protein buffer”) consisted of 20 mM Tris pH 7.4, 100 mM NaCl, 2 mM MgCl2, 10 µM GDP, and 1 mM TCEP. Growths were either done in 3 mL small-scale culture in autoinduction media (Teknova 3S2000) with kanamycin or 500 mL LB/kanamycin media containing 40 mM lactose. Cultures were grown shaking overnight for at least 16 hr at 30 °C. Following overnight growth, cells were centrifuged and media was discarded. Cell pellets were immediately resuspended in lysis buffer (protein buffer with DNAse, lysozyme, EDTA-free protease inhibitor) then directly lysed on ice by sonication. Cell debris was removed by centrifugation at 11,600 g for 40 min. Protein was captured from lysate by incubation with either agarose or magnetic Ni NTA beads (GenScript) for at least 1 hr at 4 °C. For 500 mL growths, 1.5 mL of regular beads were added to each growth, and the slurry was added to a gravity column with the flow-through discarded. For 3 mL growths, 50–100 µL of magnetic bead slurry was added and washes were done with magnetic immobilization of the beads. Three washes with protein buffer plus 20 mM imidazole were performed followed by three washes with protein buffer only. The resin-bound His-tagged fusion protein was incubated overnight at 4 °C with Cth protease (1 µM) to cleave the N-terminal affinity tag, releasing the target protein from the beads via on-resin proteolysis. Elution containing the purified cleaved protein was collected. The resulting protein was stored in protein buffer at 4 °C or flash frozen for long-term storage.

### Size Exclusion Chromatography

We performed size exclusion chromatography (SEC) to determine the oligomerization state of purified proteins using a BioRad NGC system paired with a BioRad ENrich 650 chromatography column. Proteins were concentrated to 100 µM or greater before injection, and SEC was performed in protein buffer (see above).

### Circular Dichroism Spectroscopy

Circular dichroism (CD) spectroscopy was used to determine the secondary-structure content and thermal stability of LUCI-tag. A sample of 0.1 mg/mL LUCI-tag protein was prepared in protein buffer. CD spectra were collected using a 0.1 cm path-length quartz cuvette (VWR) in a Jasco J-1500 CD Spectropolarimeter. Spectra were collected from 195 nm to 260 nm in continuous scanning mode, with a band width of 1 nm, scanning speed of 50 nm/min, data pitch of 0.1 nm, digital integration time of 1 sec, and an average of 5 accumulations. A Peltier temperature controller was used to vary the sample temperature in the range of 25 °C to 95 °C with an interval of 10 °C, an increase rate of 10 °C/min, and an average of 3 accumulations.

### In-Gel Fluorescence of Purified Proteins and Lysates

For purified protein in-gel fluorescence, 10 µL of 10 µM protein was incubated with AG–TAMRA probe before quenching with 10 µL of 4% formic acid. Loading dye was added (ThermoFisher Cat # NP0007) and run on a NuPage Bis-Tris 4-12% gel (ThermoFisher Cat # NP0322). An Amersham Typhoon was used to image the probe fluorescence, followed by Coomassie staining and imaging. For lysate gels, transformed BL21(DE3) *E. coli* were grown overnight at 30 °C in 3 mL autoinduction media and lysed by sonication. 10 µL of the soluble fraction was obtained and stained/imaged in the same manner as purified protein. MOLT4 mammalian lysates were prepared according to the same protocol described below for the immunoprecipitation assay.

### Fluorescence Anisotropy Experiments

Fluorescence anisotropy experiments were performed in protein buffer with either 10 µL/well 384-well format (C12S mutant binding experiments and slower time-course experiments) or 100 µL/well 96-well half-area plate (faster time-course experiments). Measurements were done with a BMG LabTech PHERAstar FSX using appropriate FP module filters.

For binding affinity measurements, serial dilutions of protein were performed in triplicate with 50 nM of AG–conjugate probes. For time-course experiments, 50 µL of 100 nM AG–fluorophore was first added. At measurement time, 50 µL of protein was added (bringing the AG–fluorophore concentration to 50 nM) and immediately followed by pipette mixing and plate-reader scanning. The dead time was measured manually from the end of mixing to the first reading by the plate reader. For competition experiments to measure the affinity of unconjugated adagrasib and divarasib, AG–TAMRA was first mixed with LUCI-tag C12S mutants to a final concentration of 50 nM AG–TAMRA and 1 µM protein. Serial dilutions of competitor starting from 20 µM were performed in 10 µL wells in triplicate. Gain settings for all probes were calculated by auto-adjusting the focus on a well containing free probe and buffer and setting the millipolarization (mP) of free ligand to mP = 35.

### Fluorescence Intensity Experiments

For *in vitro* measurements of fluorescence intensity changes over time, 96-well half-area plates were used with 100 µL/well total volume. 50 µL of 200 nM AG–JF646 was added, followed by a 1 hr incubation (covered) at RT to allow for equilibration of the fluorophore in the well. Following this, 50 µL of 400 nM protein was added, bringing the final concentrations to 100 nM AG–JF646 and 200 nM protein. Fluorescence intensity was measured over time for 90 min on a BMG LabTech PHERAstar FSX.

### Nucleotide exchange

Nucleotide exchange protocols to generate fully GDP- and GTP-loaded LUCI-tag and K-Ras(G12C) were adapted from existing protocols^10^. Proteins were purified as described above and buffer-exchanged from protein buffer into Exchange Buffer A (20 mM Tris pH 7.4, 100 mM NaCl, 200 mM (NH₄)₂SO₄, 25 mM EDTA, 1 mM TCEP) using 10 kDa MWCO centrifugal concentrators over three wash cycles to strip Mg²⁺ in the buffer. Each protein was split into separate GDP and GTP aliquots and loaded with a 20-fold molar excess of nucleotide (from 10 mM stocks in ice-cold water), added in two doses with rotation at 4 °C. EDTA chelation of Mg²⁺ lowered nucleotide affinity, allowing exchange to proceed by mass action. MgCl₂ was then added to a final concentration of 10 mM and incubated for 15 min at 4 °C to restore Mg²⁺ coordination and lock the exchanged nucleotide in place. Excess free nucleotide, EDTA, and salts were removed by size-exclusion chromatography into Buffer B (20 mM Tris pH 7.4, 100 mM NaCl, 2 mM MgCl₂, 1 mM TCEP), and peak protein fractions were pooled. Time-resolved fluorescence anisotropy experiments were performed as above. GDP- and GTP-loaded LUCI-tag and K-Ras(G12C), along with non-exchanged controls, were serially diluted in Buffer B and mixed with AG–AF488 to a final concentration of 1000 nM to 16 nM protein and 10 nM probe.

### Fitting *K*_d_ from fluorescence anisotropy (direct binding)

Direct binding data (Fig. 2, Supp. Fig. S18) was fit using a python script to the quadratic form of a single-site binding model:

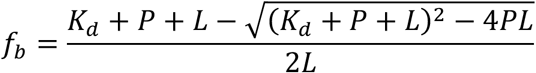

Where *f_b_* is the fraction bound, *K_d_* is the dissociation constant, *P* is protein concentration and *L* is ligand concentration (fixed at 50 nM). Anisotropy was modeled with parameters *A*, *B*:

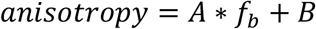

The mean anisotropy values across replicates were taken and the SciPy curve_fit package was used to fit the *K_d_*.

### Fitting *K*_d_ from fluorescence anisotropy (competition binding)

Competition binding data were collected by titrating varying amounts of unlabeled competitor ligand into a fixed amount of fluorescently-labeled ligand (50 nM) and protein (concentration determined by ∼1.5x K_d_ with fluorescently-labeled ligand). Direct-binding and competition fluorescence anisotropy data (Figs. S21 and S22) were fit in Python (scipy.optimize.least_squares) in two stages. In the first stage, the dissociation constant of the fluorescent ligand (*K_d_*_1_) was fit from a direct protein titration using the approach described above, with *L* being held constant at 50 nM.

In the second stage, *K_d_*_1_, *A*, and *B* were held fixed at their first-stage values, and competition of fluorescent ligand *L* by unlabeled inhibitor *C* for protein *P* was described by the coupled equilibria:

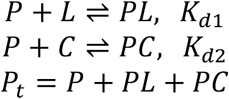

The system of equations was solved using cubic coefficients numerically derived for free protein. Code for fitting is provided in Zenodo^11^.

### Fitting *k*_app_ from time-resolved fluorescence anisotropy data

#### Fitting fast labeling kinetics with a complete binding model

AG time-course data was fit to a binding model accounting for the noncovalent and covalent binding steps, based on methods from ref. ^12^:

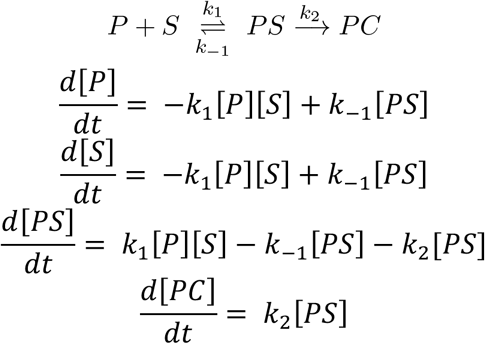

Where *P* is the protein, *S* is the AG–fluorophore probe, *PS* is the noncovalent protein–probe complex, and *PC* is the covalent complex.

Since the early timepoints are most informative, the data used for fitting was manually truncated to ∼ 3 x t_1/2_ (time taken to reach half of maximum anisotropy) to minimize overfitting the plateau phase. ODEs were solved using SciPy odeint with initial conditions *P*_0_, *S*_0_provided from experimental conditions and *PC*_0_ = 0. The anisotropy signal was mapped to fraction bound using:

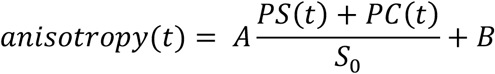

Parameters were fit globally across all concentrations using bounded nonlinear least squares (SciPy least_squares). We anchored the fit with a pseudo-point at t=0 using measured dead time. Fitted parameters were *k*_1_, *k*_−1_, *k*_2_. From fitted rate constants, the apparent second-order rate constant *k_app_* was calculated as:

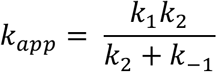

Confidence intervals were obtained by bootstrapping: the data was first fit globally, and 500 synthetic datasets were then generated by adding data residuals (resampled i.i.d. within each concentration) to the best-fit curve. Each synthetic dataset was refit globally across concentrations using a warm start at the initial optimum). Reported error bars are the 2.5th–97.5th percentiles (95% CI) of the fit *k_app_* value.

#### Fitting slow kinetics with a simplified model

Because ARS-1323 reaction kinetics were slower than AG reaction kinetics (Fig. S7), we used a simplified, analytical, single-exponential model of anisotropy over time:

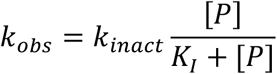

The anisotropy of the ligand-only well was subtracted from each reading due to a time-dependent baseline drift. For each protein and concentration, we fit pseudo first-order rate constant *k_obs_* using least squares with scipy.optimize.curve_fit. The *k_inact_* and *K_I_* can be fit using a hyperbolic relationship between *k_obs_* and [*P*]:

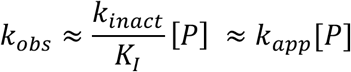

taking the slope as an approximation of *k_app_* . We note that this method provides only an estimate of *k_app_*, and was used only for the ranking of top candidates.

### X-Ray Crystallography

LUCI-tag was purified as described above. For ligand-bound structures, we incubated with 1.2x excess ligand for 1 hr with protein at 100 µM. LUCI-tag was then concentrated to 2.5 mM using 3 kDa centrifugal filters (Millipore), removing unbound ligand. We prepared the solution for co-crystallization at room temperature using the sitting drop vapor diffusion method. 0.2 µL protein was mixed 1:1 with mother liquor and set as sitting drops in a 96-well Intelli-plate (Hampton Research HR3-185) using an NT-8 Robot (Formulatrix). Crystallization conditions screened included the sparse matrix screens MCSG-1, MCSG-2, MCSG-3, and MCSG-4. Conditions that produced crystal growth are listed in Table S6.

Data was collected on a DECTRIS EIGER X 9M detector at the 17-ID-1 (AMX) beamline at the National Synchrotron Light Source II of Brookhaven National Laboratory. Raw images were indexed and merged using fastDP^13^, and the structures were phased via molecular replacement in Phenix^14^. The models were refined in Phenix and Coot^15^ with covalent ligand restraints provided by AceDrg^16^. Composite-omit maps (Figs. 3a and S26) were produced in Phenix. Table S7 statistics were also produced with Phenix.

### Switch II sequence–structure analysis using LASErMPNN

For the analysis of Fig. 3e, we started by retrieving all wild-type K-Ras and K-Ras(G12C) crystal structures from the PDB (512 protein chains from 263 entries). Because chains of the same protein within an asymmetric unit are not independent, a single representative chain was taken per entry. Among chains resolving all 20 switch II positions (residues 57–76), the chain with the lowest mean B-factor over protein atoms was selected, giving 233 entries. Because the PDB query was not restricted by sequence, the amino-acid sequence of each representative chain was read directly from the deposited coordinates and compared with canonical K-Ras4B (UniProt P01116). Sequences were compared over residues 2–150, the region in which the K-Ras4A and K-Ras4B splice isoforms are identical, so that isoform differences in the hypervariable C terminus were not counted as substitutions; residue 1 (an expression-tag remnant) and the C51/C80/C118 substitutions of cysteine-light constructs were likewise excluded. Chains carrying any substitution other than G12C were discarded, removing 63 entries (most commonly Q61H, G13D and G13C, of which 26 were substituted within switch II itself) and leaving 170 entries (106 G12C and 64 wild-type) with 90 containing a switch-II pocket inhibitor and 80 drug-free. For each representative, we used only backbone atoms (N, Cα, C, O) of residues 1–169, with side chains, nucleotide, metal and all heteroatoms removed. The K-Ras(G12C) and LUCI-tag sequences were threaded onto these backbones so that the two sequences differ only in identity and never in geometry. Sequence compatibility was scored with LASErMPNN (soluble weights, soluble_weights_no_heldout_drop_clusters_optstep_65000.pt) in fixed-sequence mode (--repack_only_input_sequence, --sequence_temp 0.2), using five independent decoding orders per structure–sequence pair. Per-residue probabilities of the threaded amino acid were read from the output B-factor field of the scored PDB files, floored to 0.005 to account for any near-zero probabilities, converted to negative log-likelihood (NLL), and averaged over the five decoding orders to give a per-residue score. The switch-II score is the mean of these over residues 57–76.

### Intact Mass Spectrometry

Protein samples were acidified and desalted over C4 resin (ZipTip, Millipore). Samples were analyzed on an Orbitrap Eclipse mass spectrometer coupled to an Ultimate 3000 RSLCnano (Thermo Fisher Scientific). Proteins were separated over a 15-cm polystyrene divinylbenzene column (ES907, Thermo Fisher Scientific) with a 15-min gradient of 10–50% acetonitrile in 0.1% formic acid and electrosprayed (2 kV, 305°C) with an EasySpray ion source. Scans (600–2,000 m/z) were obtained in the orbitrap (15,000 resolution, profile). Deconvolution of the spectra and relative quantification of the peaks was done in the UniDec software. Percent labeling of proteins was calculated based on the relative ratio of labeled species to the parent species.

### Immunoprecipitation-Mass Spectrometry (IPMS) Binding Assay

#### Preparation of FLAG–tagged constructs

A C-terminal 1x FLAG tag was fused to the protein sequence separated by a 2x GGS (sequence-GGS-GGS-DYKDDDDK). FLAG–tagged LUCI-tag and FLAG–tagged K-Ras(G12C) (the Cys-light version) were cloned, expressed in *E. coli*, and purified in protein buffer using the methods described above.

#### Immunoprecipitation

A total of 1 × 10^7^ MOLT4 cells per IP were collected and lysed in lysis buffer (50 mM Tris pH 8, 200 mM NaCl, 2 mM TCEP, 0.1% NP-40, 10 units turbonuclease/200 µL buffer, 1x cOmplete protease inhibitor tablet/5 mL buffer) and sonicated on ice for 5 rounds of 2 sec followed by 10 sec pauses at 25% amplitude. After centrifugation clarification, lysate was transferred to new lobind tubes. 10 µg of FLAG–K-Ras(G12C) or FLAG–LUCI-tag were added to each lysate and incubated with end-over-end rotation for 1 hr in the cold room. 20 µL of pre-washed and resuspended anti-FLAG magnetic agarose slurry (Pierce) was added to each sample and incubated with end-over-end rotation for 1 hr in the cold room. Beads were washed three times with wash buffer (50 mM Tris pH 8, 2 mM TCEP, 0.1% NP-40, 1`×` cOmplete protease inhibitor tablet/5 mL buffer) followed by three non-detergent washes (50 mM Tris pH 8, 2 mM TCEP, 1`×` cOmplete protease inhibitor tablet/5 mL buffer) before elution with 0.1 M Glycine-HCl, pH 2.7. Tris (pH 8.5) was added to elution to reach a pH of 8. Samples were then reduced with 10 mM TCEP for 30 min at room temperature, followed by alkylation with 15 mM iodoacetamide for 45 min at room temperature in the dark. Alkylation was quenched by the addition of 10 mM DTT. Samples were digested with 2 μg LysC and 1 μg Trypsin overnight at 37 °C. Sample digests were acidified with formic acid to a pH of 2–3 prior to desalting using C18 solid phase extraction plates (SOLA, Thermo Fisher Scientific). Desalted peptides were dried in a vacuum centrifuge and reconstituted in 0.1% formic acid for LC-MS analysis.

#### Label free quantitative mass spectrometry with diaPASEF and data analysis

Data were collected using a TimsTOF HT or TimsTOF Ultra2(Bruker Daltonics, Bremen, Germany) coupled to a nanoElute2 LC pump (Bruker Daltonics, Bremen, Germany) via a CaptiveSpray nano-electrospray source. Peptides were separated on a reversed-phase C_18_ column (25 cm x 75 µm ID, 1.6 µM, IonOpticks, Australia) containing an integrated captive spray emitter. Peptides were separated using a 40 min gradient of 2–30% buffer B (acetonitrile in 0.1% formic acid) with a flow rate of 250 nL/min and column temperature maintained at 50 °C.

The TIMS elution voltages were calibrated linearly with three points (Agilent ESI-L Tuning Mix Ions; 622, 922, 1,222 *m/z*) to determine the reduced ion mobility coefficients (1/K_0_). To perform diaPASEF, we used py_diAID^17^, a python package, to assess the precursor distribution in the *m/z*-ion mobility plane to generate a diaPASEF acquisition scheme with variable window isolation widths that are aligned to the precursor density in *m/z*. Data was acquired using twenty cycles with three mobility window scans each (creating 60 windows) covering the diagonal scan line for doubly and triply charged precursors, with singly charged precursors able to be excluded by their position in the *m/z*-ion mobility plane. These precursor isolation windows were defined between 350–1250 *m/z* and 1/k0 of 0.6–1.45 V.s/cm^2^.

The diaPASEF raw file processing and controlling peptide and protein level false discovery rates, assembling proteins from peptides, and protein quantification from peptides was performed using library free analysis searched against a Swiss-Prot human database (January 2021) in DIA-NN 1.8^18^. Database search criteria largely followed the default settings for directDIA including: tryptic with two missed cleavages, carbamidomethylation of cysteine, and oxidation of methionine and precursor Q-value (FDR) cut-off of 0.01. Precursor quantification strategy was set to Robust LC (high accuracy) with RT-dependent cross run normalization. Resulting data was filtered to only include proteins that had a minimum of 3 counts in at least 4 replicates of each independent comparison of K-Ras redesigns to the K-Ras(G12C) control. Protein abundances were globally normalized using in-house scripts in the R framework (R Development Core Team, 2014). Proteins with missing values were imputed by random selection from a Gaussian distribution either with a mean of the non-missing values for that treatment group or with a mean equal to the median of the background (in cases when all values for a treatment group are missing)^19^. Protein abundances were scaled and significant changes comparing the relative protein abundance of K-Ras redesign (e.g., LUCI-tag) to the K-Ras(G12C) control and comparisons were assessed by moderated t-test as implemented in the limma package within the R framework^20^.

### RAS Signaling Western Blots

For analysis of signaling through RAS effector pathways, HEK293T cells were transiently transfected with plasmids carrying GFP-tagged K-Ras or LUCI-tag using PolyJet (SignaGen Laboratories #SL100688) according to the manufacturer’s instructions. 24 hr after transfection, cells were lysed in 1x MLB buffer (Millipore 20-168) containing EDTA-free protease inhibitor cocktail (Roche 11836170001) and phosphatase inhibitor cocktails 2 and 3 (Sigma P5726 and P0044, respectively). Whole cell lysates were quantified and normalized using the Pierce BCA assay (Thermo Scientific 23225).

Western blotting was performed with 40 µg protein per lane on 4–15% Mini-PROTEAN TGX precast gels (BioRad 4561085) and transferred to nitrocellulose membranes using the Trans-Blot Turbo Transfer system (BioRad 1704150) according to the manufacturer’s instructions. Membranes were blocked for 1 hr at room temperature with Intercept (TBS) blocking buffer (LICORBio 927-60001) and then incubated overnight at 4 °C with primary antibodies diluted 1:1000 (unless otherwise indicated) in Intercept T20 (TBS) antibody diluent (LICORBio 927-65001). Primary antibodies used were as follows: Ras10 (Sigma-Aldrich 05-516), pan Akt (Cell Signaling Technology 2920), phospho-Akt (Cell Signaling Technology 4060), MEK1/2 (Cell Signaling Technology 4694), phospho-MEK1/2 (Cell Signaling Technology 9121), ERK1/2 (Cell Signaling Technology 4696), phospho-ERK1/2 (Cell Signaling Technology 4370), GFP (1:5000) (Abcam ab13970), vinculin (Sigma-Aldrich V9131). The following day, appropriate secondary antibodies were diluted 1:10,000 in Intercept antibody diluent and incubated with membranes for 1 hr at room temperature prior to visualization on an Odyssey LiCOR CLx machine. Secondary antibodies used were AlexaFluor 680 goat anti-mouse IgG (Invitrogen A-21058), AlexaFluor 800 goat anti-rabbit IgG (Invitrogen A-2633284), and AlexaFluor 680 goat anti-chicken IgG (Invitrogen A-32934).

### Chemistry Methods

#### Synthesis of LUCI-tag Probes *Materials*

Unless otherwise noted, reagents and solvents were purchased from commercial suppliers and used as received. TAMRA-azide was purchased from Lumiprobe (C7130). JF646-azide was purchased from MedChemExpress (HY-131027). AF488-azide was purchased from Lumiprobe (21830). Adagrasib-alkyne was purchased from MedChemExpress (HY-134654), as was ARS-1323-alkyne (HY-128522).

#### General synthetic chemistry methods

All reactions were monitored using a Waters Acquity UPLC-MS system (Waters Acquity QDa Detector, Waters Acquity PDA eλ Detector, Waters Acquity-I UPLC class Sample Manager-FTN, Waters Acquity-I UPLC class Binary Solvent Manager) using Acquity UPLC® BEH C18 column (2.1 x 50 mm, 1.7 μm particle size): solvent gradient 5% to 99% acetonitrile in water (0.035% TFA as additive); flow rate: 0.6 mL/min. Purification of reaction products was carried out using a Waters HPLC system using SunFireTM C18 column (19 x 100 mm, 5 μm particle size): solvent gradient 0% to 99% acetonitrile in water (0.035% TFA as additive); flow rate: 20 mL/min. The purity of all compounds was over 95% by UV and was analyzed with Waters UPLC system.

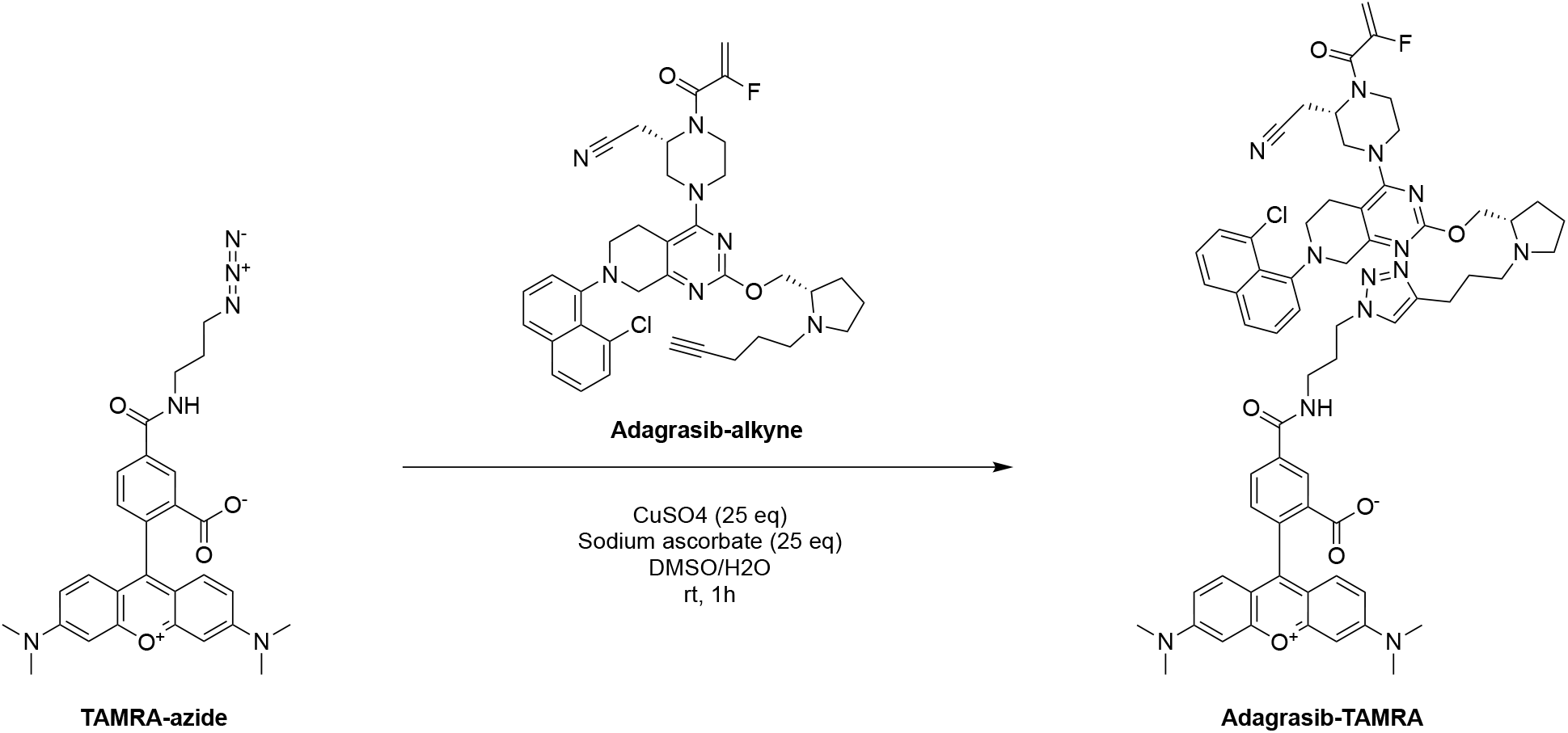

TAMRA-azide was dissolved in DMSO to a stock concentration of 20 mM. Adagrasib-alkyne was dissolved in DMSO to a stock concentration of 10 mM. CuSO_4_ and L-ascorbic acid were dissolved in MilliQ water to stock concentrations of 1 M. To a vial was added 300 µL of TAMRA-azide, 300 µL of Adagrasib-alkyne, and 600 µL DMSO. Then, 150 µL each of CuSO_4_ and L-ascorbic acid were added. After 15 minutes, the reaction was diluted with DMSO, filtered, and purified by RP-HPLC under acidic conditions to afford analytically pure Adagrasib-TAMRA. Pink powder. Calc’d [M+H]^+^: 1167.50. Found [M+H]^+^: 1168.56.

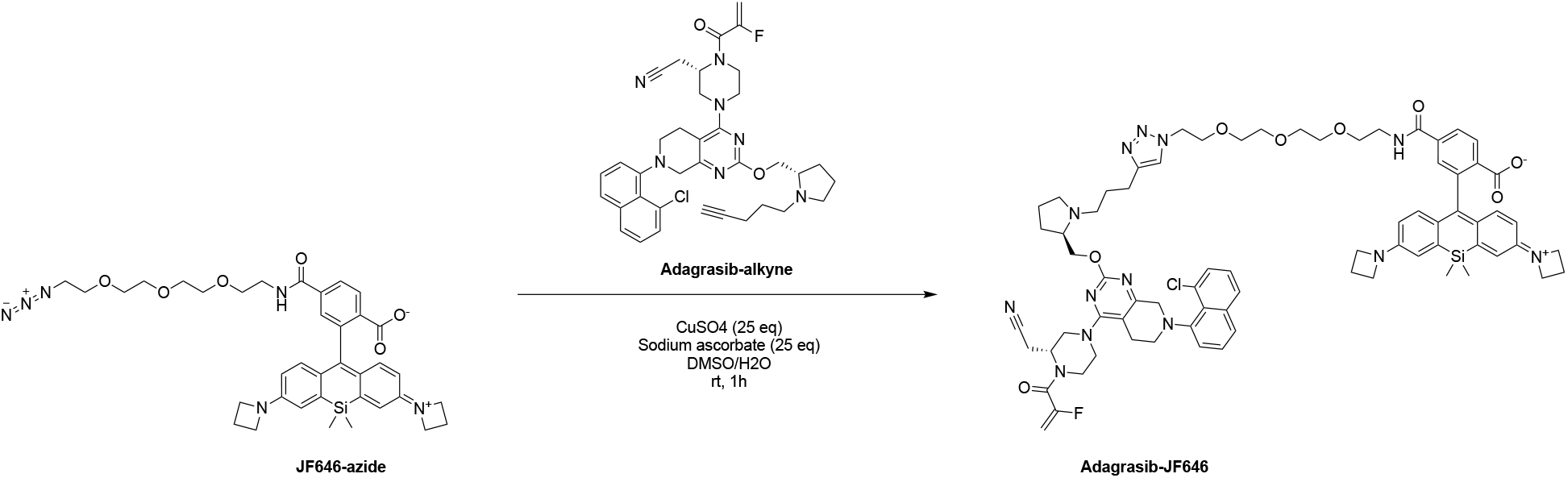

JF646-azide was dissolved in DMSO to a stock concentration of 10 mM. Adagrasib-alkyne was dissolved in DMSO to a stock concentration of 10 mM. CuSO_4_ and L-ascorbic acid were dissolved in MilliQ water to stock concentrations of 1 M. To a vial was added 200 µL of JF646-azide, 100 µL of Adagrasib-alkyne, and 200 µL DMSO. Then, 50 µL each of CuSO_4_ and L-ascorbic acid were added. After 15 minutes, the reaction was diluted with DMSO, filtered, and purified by RP-HPLC under acidic conditions to afford analytically pure Adagrasib-JF646. Blue powder. Calc’d [M+H]^+^: 1353.59. Found [M+2H]^2+^: 677.52.

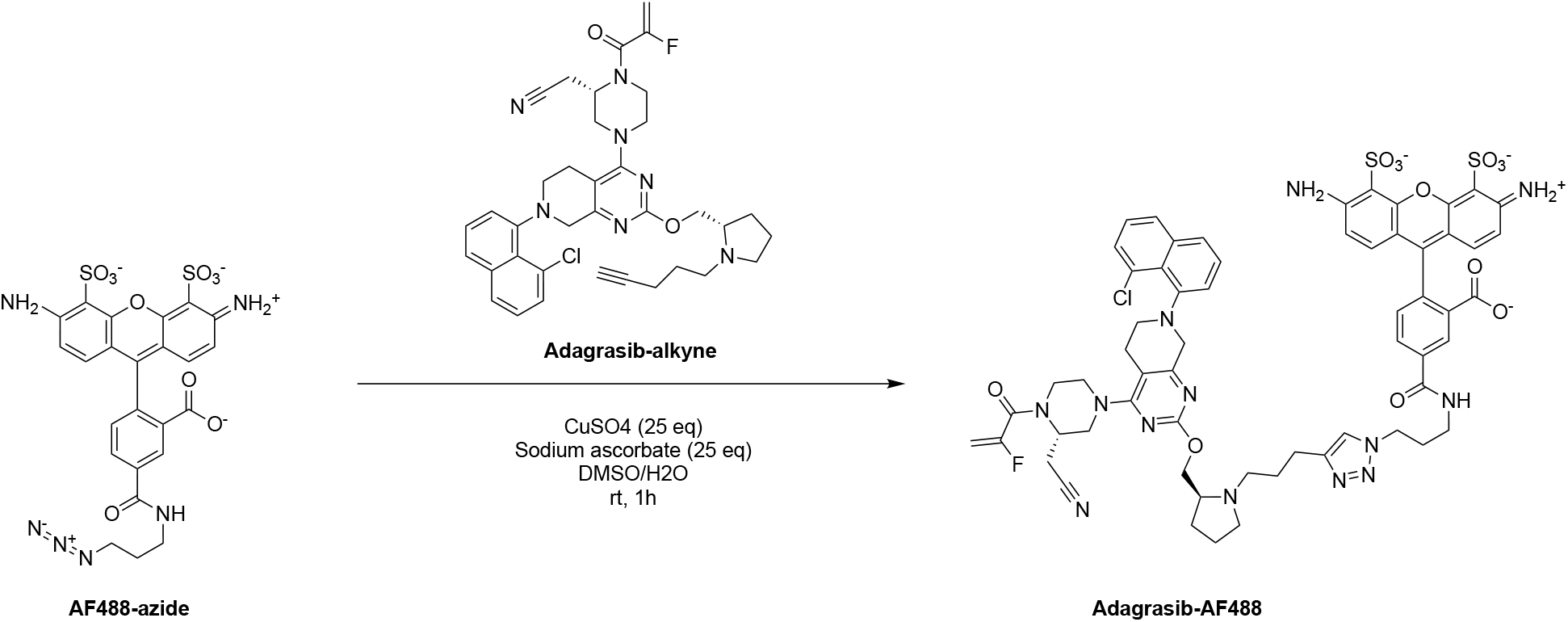

AF488-azide was dissolved in DMSO to a stock concentration of 10 mM. Adagrasib-alkyne was dissolved in DMSO to a stock concentration of 10 mM. CuSO_4_ and L-ascorbic acid were dissolved in MilliQ water to stock concentrations of 1 M. To a vial was added 200 µL of AF488-azide, 100 µL of Adagrasib-alkyne, and 200 µL DMSO. Then, 50 µL each of CuSO_4_ and L-ascorbic acid were added. After 15 minutes, the reaction was diluted with DMSO, filtered, and purified by RP-HPLC under acidic conditions to afford analytically pure Adagrasib-AF488. Yellow powder. Calc’d [M+H]^+^: 1273.34. Found [M+2H]^2+^: 636.83.

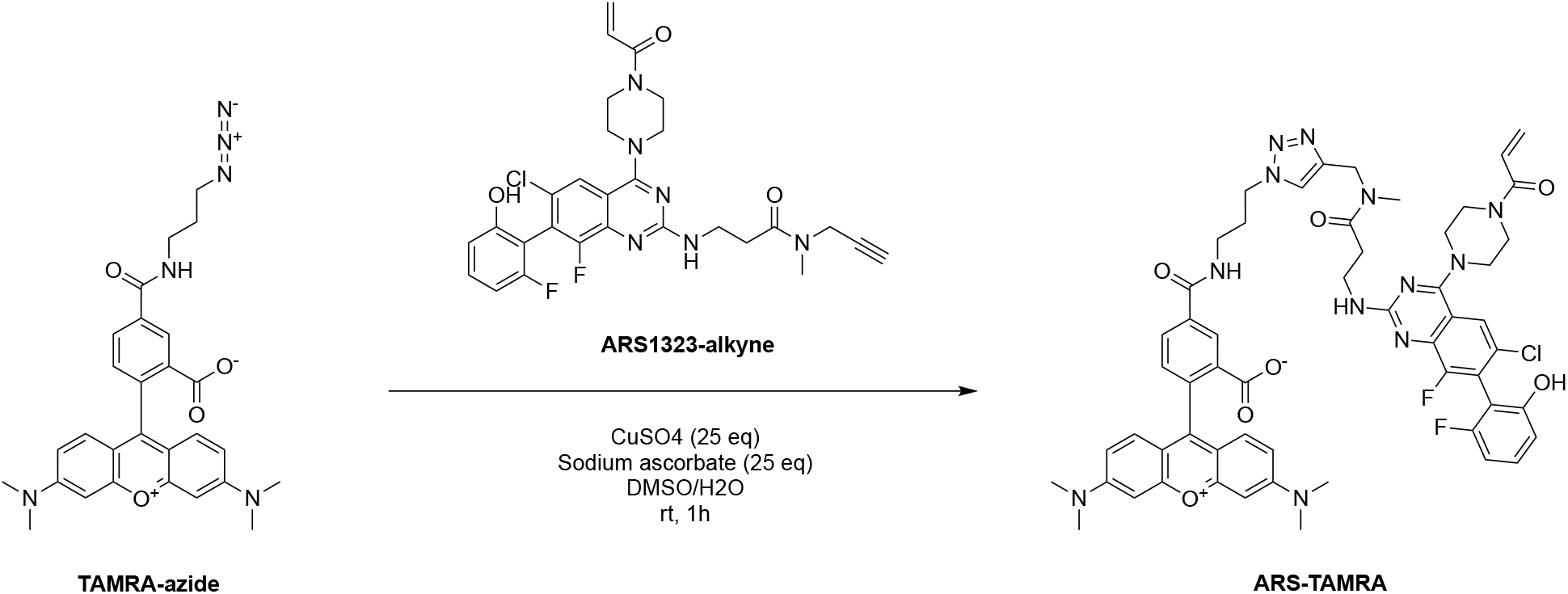

TAMRA-azide was dissolved in DMSO to a stock concentration of 20 mM. ARS-1323-alkyne was dissolved in DMSO to a stock concentration of 10 mM. CuSO_4_ and L-ascorbic acid were dissolved in MilliQ water to stock concentrations of 1 M. To a vial was added 300 µL of TAMRA-azide, 300 µL of ARS-1323-alkyne, and 600 µL DMSO. Then, 150 µL each of CuSO_4_ and L-ascorbic acid were added. After 15 minutes, the reaction was diluted with DMSO, filtered, and purified by RP-HPLC under acidic conditions to afford analytically pure Adagrasib-TAMRA. Pink powder. Calc’d [M+H]^+^: 1081.40. Found [M+H]^+^: 1081.63.

### Purification-free “Direct to Biology” Synthesis of Probes

JF646-azide and AG-alkyne were dissolved in DMSO to prepare 10 mM stock solutions. THPTA or TBTA was dissolved in DMSO to prepare a 25 mM stock solution. CuSO_4_ and L-ascorbic acid were dissolved in MilliQ water to prepare 100 mM stock solutions. 5 µL each of JF646-azide and AG-alkyne were added to a tube. 20 µL of THPTA/TBTA was added. Finally, 5 µL each of CuSO_4_ and L-ascorbic acid were added. The tube was flicked well and incubated at room temperature in the dark. The reaction was monitored by UPLC-MS after 5, 15, and 30 min. Reactions looked completed by 30 min by UV (Supp. Figs. S13, S14). The mixture was stored at -80 °C until further use.

To assist users with probe synthesis, we provide Table S8 with a list of reagents and concentrations/amounts. While this protocol was validated with JF646 dye, it should generalize to other AG-fluorophore conjugates. Stocks should be prepared carefully with high-accuracy scales when possible to obtain 1:1 azide-alkyne stoichiometry. We observed that copper protectants TBTA and THPTA (a water-soluble equivalent) performed equally well and were both usable in live-cell microscopy (Fig. S43).

### Mammalian Live-Cell Microscopy Methods

#### Plasmid Construction and Cloning

We codon-optimized LUCI-tag for human expression and cloned them into mammalian backbones. Original plasmids (pDC713) were digested with EcoRI and BamHI (NEB, R3101 and R3136 respectively) in 1x rCutSmart buffer (NEB, #B6004S) at 37 °C for 1 hr. Digested products were purified using SPRI bead clean-up (1.8× bead ratio) and used as linearized vectors. Insert sequences were PCR-amplified and purified by SPRI beads. Vector and insert fragments were assembled using the NEBuilder HiFi DNA Assembly Master Mix (New England Biolabs, E2621L) following the manufacturer’s instructions. Assembled plasmids were transformed into E. coli NEB Stable Competent cells (New England Biolabs, C3040H) and verified by whole plasmid sequencing. The nuclear-localized protein (PITX2) and its corresponding expression backbone came from the Multiplexed Overexpression of Regulatory Factors (MORF) Library, which was a gift from Feng Zhang (Addgene #1000000218).

### Mammalian cell culture

HEK293FT cells (referenced as HEK293T in the text and figures, Thermo Fisher Scientific, R70007) were maintained in Dulbecco’s Modified Eagle Medium (DMEM) with GlutaMAX (Gibco, 10566016) supplemented with 10% (v/v) fetal bovine serum (FBS; Sigma-Aldrich, F4135) and 1% penicillin–streptomycin (Gibco, 15140122). Cultures were incubated at 37 °C in a humidified atmosphere containing 5% CO₂. Cells were seeded in Matrigel-coated (Corning, 356230) SensoPlate 96-well glass-bottom plates (Greiner, 655892) at a density of 30,000 cells/well.

### General protocol for live-cell fluorescence microscopy of LUCI-tag in live mammalian cells

After 24 hr following seeding, transfection was performed with TransIT-LT1 (Mirus Bio, MIR2304) following the manufacturer’s protocol. Each well received 100 ng plasmid DNA per construct. 48 hr post-transfection, cells were incubated with ligand (generally 500 nM) and Hoechst (1:1000) (Thermo Fisher Scientific, H1399) in phenol red–free DMEM (Gibco, 31053028) and imaged using a Dragonfly 200 spinning-disk confocal microscope (Andor, Oxford Instruments) at 40x using a water-immersion lens. Images were collected with the 405nm, 488nm, 561 nm, and 646 nm lasers with the corresponding filters and bandwidths 445-46, 521-38, 594-43, 685-47 nm. The maximum eGFP signal was used for focusing. Microscopy images were processed in a custom Python script and Fiji/ImageJ. All images were processed identically in situations where direct comparison across experimental conditions is shown.

#### Live-cell fluorescence microscopy of single-construct transfected cells

For each well, 100 ng of plasmid between 0.3–1.0 *μ*g/*μ*l concentration was added to 10 *μ*l of Opti-MEM I Reduced Serum Medium (Gibco, 31985062) without antibiotics, followed by addition of 0.3 *μ*l of TransIT-LT1 (Mirus Bio, MIR2304) in a 200 *μ*l strip-tube while avoiding the walls and mixed by gentle pipetting, incubated at room temperature for 30 min then added drop-wise to cells. 48 hr post-transfection, GFP expression was confirmed when appropriate. Dye solution was freshly prepared in imaging media from DMSO stocks at 500 nM unless otherwise noted in figures. 100 µL of staining volume was used. The plate was placed back in the 37 °C incubator. For JF646 dyes, no washing or replacement of media was performed before imaging.

#### Live-cell fluorescence microscopy of dual- and tri-color microscopy

For dual- and tri-color microscopy, cells were co-transfected with multiple plasmids similar to single-construct transfection described above. For each construct, 100 ng of plasmid DNA was used together with 0.3 *μ*l of TransIT-LT1. All plasmid DNA was added and mixed in the reduced serum medium before addition of transfection reagent.

Staining was performed by combining dyes together for a 500 nM-per-dye final concentration in imaging media. Due to the background of AF488, JF549, and TAMRA dyes for HaloTag7 and SNAP-tag, cells were washed two times in 100 µL PBS and then incubated for at least 30 min in 100 µL of imaging media to allow excess dye to exit. This was followed by 3 more washes in 100 µL PBS before media was again replaced with 100 µL imaging media.

#### Live-cell fluorescence microscopy of membrane-localized constructs

Existing plasmids for HaloTag7–B2AR and TFR1–HaloTag7 were PCR-linearised to remove HaloTag7 then purified using AMPure XP beads at 1.8x ratio and used as vectors. Primers were used on previously-ordered eblocks containing LUCI-tag constructs to generate inserts with homology overlap to the vectors and also purified by AMPure XP beads at 1.8x ratio. Vector and insert fragments were assembled using the NEBuilder HiFi DNA Assembly Master Mix following manufacturer’s instructions, transformed into E. coli NEB Stable Competent cells and verified by whole plasmid sequencing.

Staining was performed using AG–AF488 or CA-AF488 HaloTag7 dye (Promega, G1001). Three 100 µL PBS washes were performed followed by replacement with 100µL of imaging media.

#### Measurement of fluorescence intensity over time in live-cell microscopy

We used a construct consisting of GFP-SLP-PITX2 where SLP was either LUCI-tag, HaloTag7, or SNAP-tag, and PITX2 served as the nuclear localization signal. Unlike previous constructs with GFP-T2A, this construct had nuclear-localized GFP to serve as an expression normalization signal. Image acquisition was performed on a Nikon Ti2-E inverted microscope equipped with a CSU-W1 spinning disk confocal and an Andor Zyla 4.2 sCMOS camera with a 20x objective.

Images were collected of each well every two min for a total of two hr. A robotic plate mover was used to move between wells containing LUCI-tag nuclear, SNAP-tag nuclear, and HaloTag7 nuclear. Autofocus was used before every well and timestep to determine the Z-plane with the highest GFP fluorescence and to prevent drift over time. The laser intensity and exposure of the JF646 channel was adjusted so the maximum intensity value was reached with a pre-stained well expressing HaloTag7 and stained with CA-646.

#### Analysis of nuclear intensities for concentration- and time-dependent data

For the time and concentration series of Fig. S30, whole-cell segmentation was performed with Cellpose v3.1.1 using the pretrained “cyto3” model with the GFP channel as cytoplasmic marker and Hoechst as the nuclear marker (expected diameter 30 µm, 199 px; flow threshold 0.4; cell-probability threshold 0.0). For time-lapse images of Fig. S34, nuclei were segmented in the GFP channel at every timepoint using Cellpose (‘nuclei’ model, fixed diameter of 16 px, flow threshold 0.4 and cell-probability threshold 0.0). For the time lapse, segmentation was performed independently on all 61 frames of each of the three recordings (183 frames in total).

For each cell, mean intensities were measured in each channel within the nuclear or cytoplasmic mask. A per-image, per-channel background was subtracted, defined as the median intensity of pixels more than 8 px from any segmented nucleus. Cells in which ≥1% of pixels were at the ceiling in either channel were excluded from the quantification.

For the time course data, nuclei were tracked between frames by greedy nearest-centroid linking. Time courses were computed on a fixed cohort of nuclei that were detected and unsaturated in all 61 frames (n = 43, n = 207, and n = 156 nuclei for HaloTag7, SNAP-tag, and LUCI-tag, respectively). Intensity ratios were reported as the background-corrected JF646 intensity of a nucleus at time t divided by the background-corrected GFP intensity of that same nucleus at t = 0. We took the first GFP timepoint to avoid effects of GFP photobleaching over repeated imaging rounds.

### Supplementary Discussion

#### Use of LUCI-tag with cells harboring K-Ras(G12C)

One potential concern for any inhibitor-derived tag is competition with endogenous targets. In the case of K-Ras(G12C), the G12C mutation is rare in most standard cell lines such as HEK and HeLa^21^ and absent in healthy tissue, reducing the risk of off-target engagement in most common experimental systems. In contexts where the KRAS(G12C) gene is present, such as patient-derived xenografts or engineered cancer models, LUCI-tag can still be used because its significantly accelerated rate of labeling outcompetes labeling of K-Ras(G12C). In addition, K-Ras(G12C) is around half as fluorogenic as LUCI-tag, meaning that any background from K-Ras(G12C) off-target staining would be low (Fig. S33). To demonstrate this, we showed clear labeling of LUCI-tag in H358 cells, a G12C heterozygous cell line. While transfection and expression was lower (as expected for this cell line), we observed clear nuclear and cytoplasmic staining in GFP+ transfected cells stained with AG–JF646 (Fig. S42a). We also performed live-cell microscopy of HEK293T cells transfected with K-Ras(G12C) plasmid and stained with AG–JF646, observing minimal staining (Fig. S42b).

#### Inhibitor binding after GDP/GTP nucleotide exchange

We prepared K-Ras(G12C) and LUCI-tag uniformly loaded with either GTP or GDP. AG probes showed minimal labeling of GTP-exchanged K-Ras(G12C), whereas GDP loading increased labeling kinetics relative to the unexchanged protein (Fig. S24). This is consistent with previous results showing inhibitor-labeled K-Ras(G12C) binds GDP preferentially over GTP^22^. In contrast, LUCI-tag retained substantial labeling activity in the GTP-bound state, labeling only 5x slower than GDP-bound LUCI-tag. These results indicate that our sequence redesign has made an accessible Switch-II-pocket independent of nucleotide state. Whereas GTP binding normally shifts K-Ras towards an inaccessible Switch-II pocket, LUCI-tag appears to retain a binding-competent pocket across both nucleotide states.

LUCI-tag labeling is reminiscent of recently developed “dual-state” covalent inhibitors have substantial reactivity for both GDP- and GTP-bound K-Ras(G12C). The trajectory of the acrylamide warhead typically clashes with the ɣ-phosphate of the GTP; dual-state inhibitors overcome this with extremely tight affinity and allosteric remodeling of the Sw-I loop^23,24^. The structure of LUCI-tag shows evidence of similar Sw-I remodeling in both ligand-free and adagrasib-bound states (Fig. 3f). In addition, the Thr35 mutation in K-Ras (as in LUCI-tag) removes the sidechain coordination to the ɣ-phosphate of GTP, allowing switch I to adopt an inactive conformation that does not impede the switch-II pocket when GTP is bound^25^.

### Crystallographic contacts of the switch-II region of ligand-free LUCI-tag

Switch II of ligand-free K-Ras is dynamic and is therefore rarely resolved in crystal structures in the PDB^25^ (Fig. S26). The exceptions are structures 8TXK and 4LDJ, whose switch-II regions include extensive contacts with neighboring chains in the crystal. In ligand-free LUCI-tag, the switch-II region was also crystallographically resolved, prompting us to investigate the role of crystallographic contacts in stabilizing its structure.

Switch II of LUCI-tag (chain A) is largely solvent exposed without many direct crystallographic contacts, with a solvent-accessible surface area (SASA) of 847 Å^2^. In contrast, switch II in K-Ras(G12C) structures 8TXK and 4LDJ show a higher degree of burial between neighboring chains and substantially less SASA of 398 Å^2^ and 521 Å^2^, respectively (Fig. S46). Thus, LUCI-tag’s resolved switch-II region could plausibly be due to structural preorganization through sequence redesign rather than from extensive crystal contacts.

### Crystal structure of LUCI-tag with ARS-1323

We determined an X-ray crystal structure of LUCI-tag bound to the switch-II inhibitor ARS-1323 (2.8 Å resolution), which showed clear evidence of the anticipated binding mode and covalent-bond formation (Fig. S45a). ARS-bound LUCI-tag has a very low C⍺ RMSD (0.97 Å) to the crystal structure of ARS-bound K-Ras(G12C) and recapitulates almost all of the sidechain rotamers involved in ligand binding (Fig. S45b). ARS-bound LUCI-tag also has a low RMSD to inhibitor-free LUCI-tag (Fig. S45c). (Note that LUCI-tag is co-crystalized with ARS-1323, whereas K-Ras(G12C) is co-crystallized with ARS-1620; ARS-1323 is a racemic mixture containing active *S*-atropisomer ARS-1620.) The ARS- and AG-bound structures of LUCI-tag confirm that its switch-II pocket can bind multiple covalent inhibitors in their native binding mode.

### Design of LUCI-tag Cofactor-Free (CF)

The goal of LUCI-tag–CF was to redesign the sequence of K-Ras(G12C) to additionally remove binding of the GDP-Mg²⁺ cofactor, allowing the cofactor-binding site to be extensively reconfigured. This represents a nontrivial design challenge, as Mg²⁺ functions as a global conformational gatekeeper in K-Ras function^26^. Recent NMR studies show that Mg²⁺ removal promotes a highly destabilized state that weakens GDP binding and promotes GDP to GTP exchange. Structurally, Mg²⁺ adopts an octahedral coordination geometry involving the GDP β-phosphate, Ser17, Asp57, and ordered waters. This arrangement is preserved in LUCI-tag but deliberately disrupted in LUCI-tag–CF through S17H and D57A mutations. Prior work has shown that perturbation of Ser17 alone is sufficient to drive Mg²⁺ dissociation^27^ and impair function^26^. Additionally, an A18E substitution introduces electrostatic repulsion with the GDP phosphates, further disfavoring nucleotide binding. Together, these changes destabilize LUCI-tag–CF for binding GDP-Mg²⁺.

We performed time-resolved fluorescence polarization experiments for LUCI-tag–CF with AG– TAMRA under both standard buffer conditions containing Mg²⁺ and GDP, and in cofactor-free protein buffer supplemented with EDTA (20 mM Tris pH 7.4, 10 mM NaCl, 1 mM TCEP, 1 mM EDTA). LUCI-tag–CF exhibits similar labeling kinetics in both conditions, whereas LUCI-tag labels 100x more slowly without GDP-Mg²⁺. While LUCI-tag–CF labels more slowly than LUCI-tag under the most favorable conditions for each, LUCI-tag–CF is 10-fold faster than LUCI-tag in cofactor-depleted buffers (Fig. S25).

“Direct to Biology” probes in live-cell mammalian imaging

To make LUCI-tag technology more accessible, we developed a “direct to biology” protocol for probe conjugation that does not require purification via liquid chromatography. The protocol uses equimolar amounts of AG–alkyne and JF-646-azide, as well as reduced copper and ascorbic acid (see Table S8 and Chemistry Methods for full protocols and reagent lists). LC-MS showed the reaction is complete within 30 min at room temperature (Figs. S13 and S14). A direct comparison of HPLC-purified and “direct to biology” probes in live-cell microscopy showed indistinguishable staining brightness and quality, with no off-target labeling (Fig. S43).

### Additional IP-MS results

We performed IP-MS on 4 additional K-Ras(G12C) designs, observing a marked depletion of K-Ras(G12C) binding partners for each (Fig. S29). LUCI-tag–CF showed a strong loss of native K-Ras binding interactions and one potential neointeraction, a zinc finger IKZF5. Each design, including designs 19, 39, and 21, had depleted interactions of RAFs and RAS proteins compared to K-Ras(G12C).

**Figure S1.**
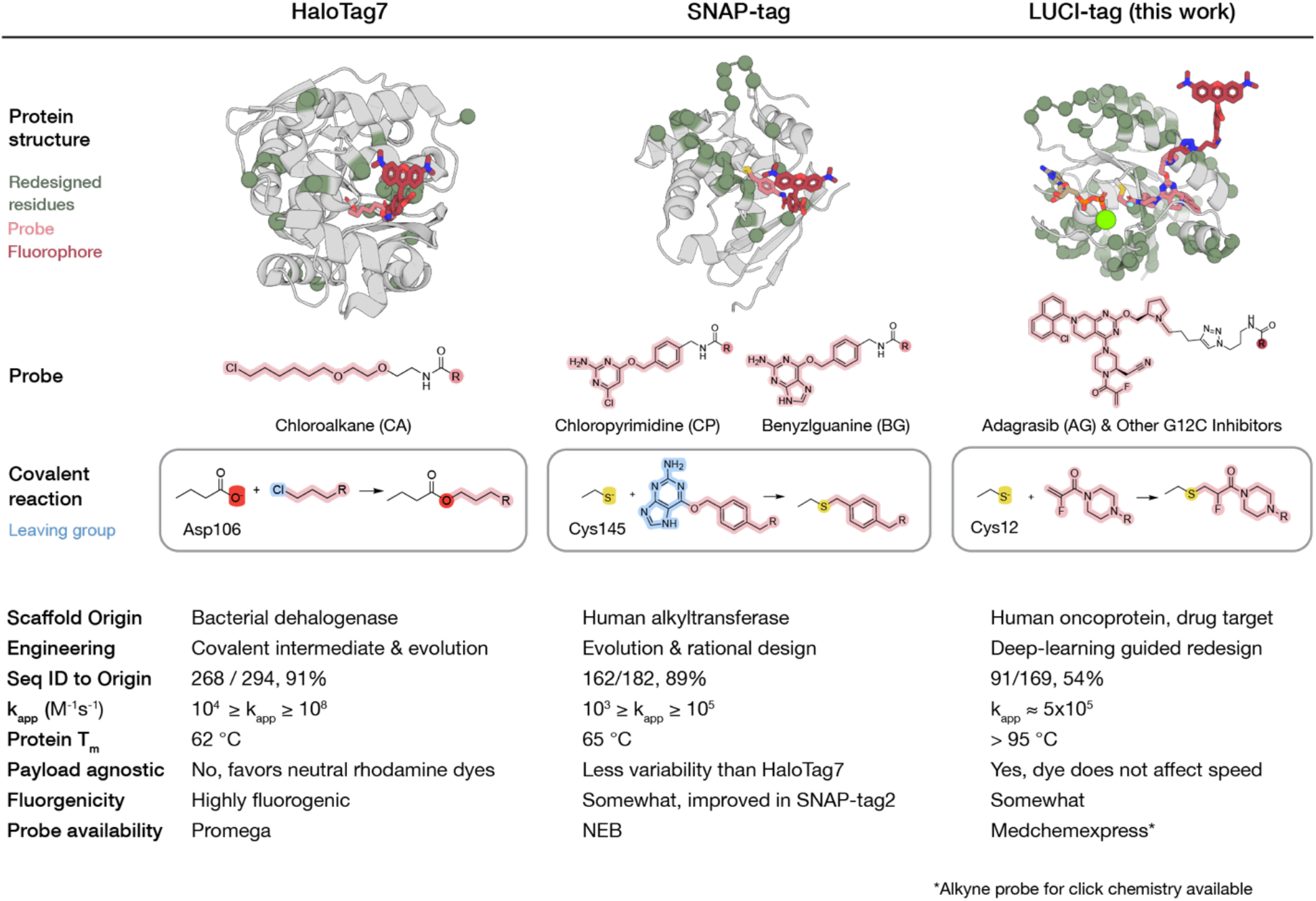
Properties of HaloTag7, SNAP-tag, and LUCI-tag. For each SLP, residues that differ from the original protein are shown as green spheres. The covalent reaction between the probe and protein sidechain is shown under each protein, with leaving groups in blue. The Michael addition in LUCI-tag does not have a leaving group. The “Probe availability” row displays the primary vendor selling commercially available probes for each tag. Protein structures displayed at the top are either from the PDB (6Y8P for SNAP-tag; 6Y7A for HaloTag7); or, for LUCI-tag, is from the crystal structure of AG–bound LUCI-tag, with the linker and rhodamine dye (dark red) modeled for visual purposes.

**Figure S2.**
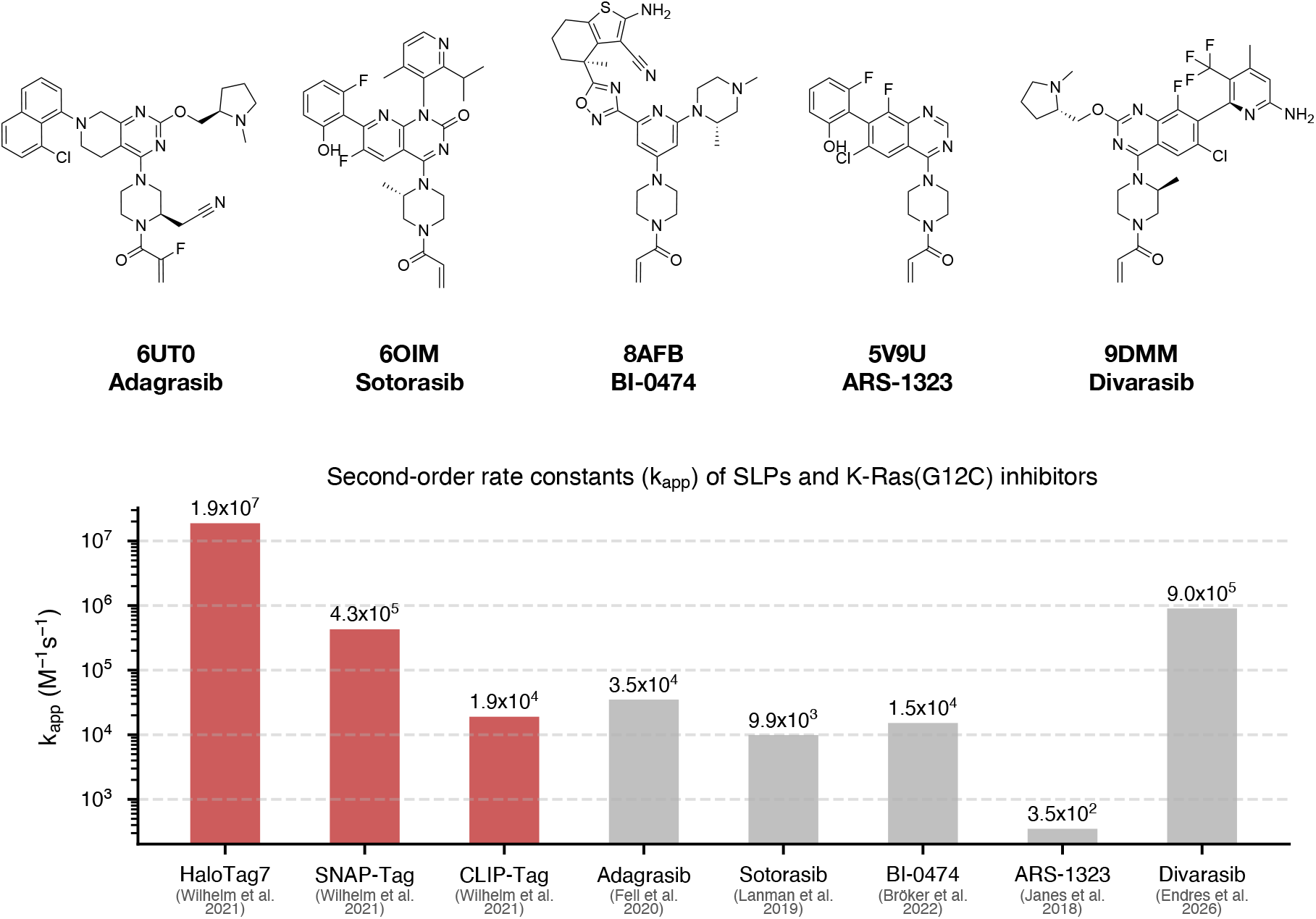
K-Ras(G12C) inhibitors and their second-order rate constants. Second-order rate constants of HaloTag7, SNAP-tag, and CLIP-tag (with their respective ligands) are also shown (red). PDB accession codes of crystal structures of inhibitor-bound K-Ras(G12C) are shown above the inhibitor name.

**Figure S3.**
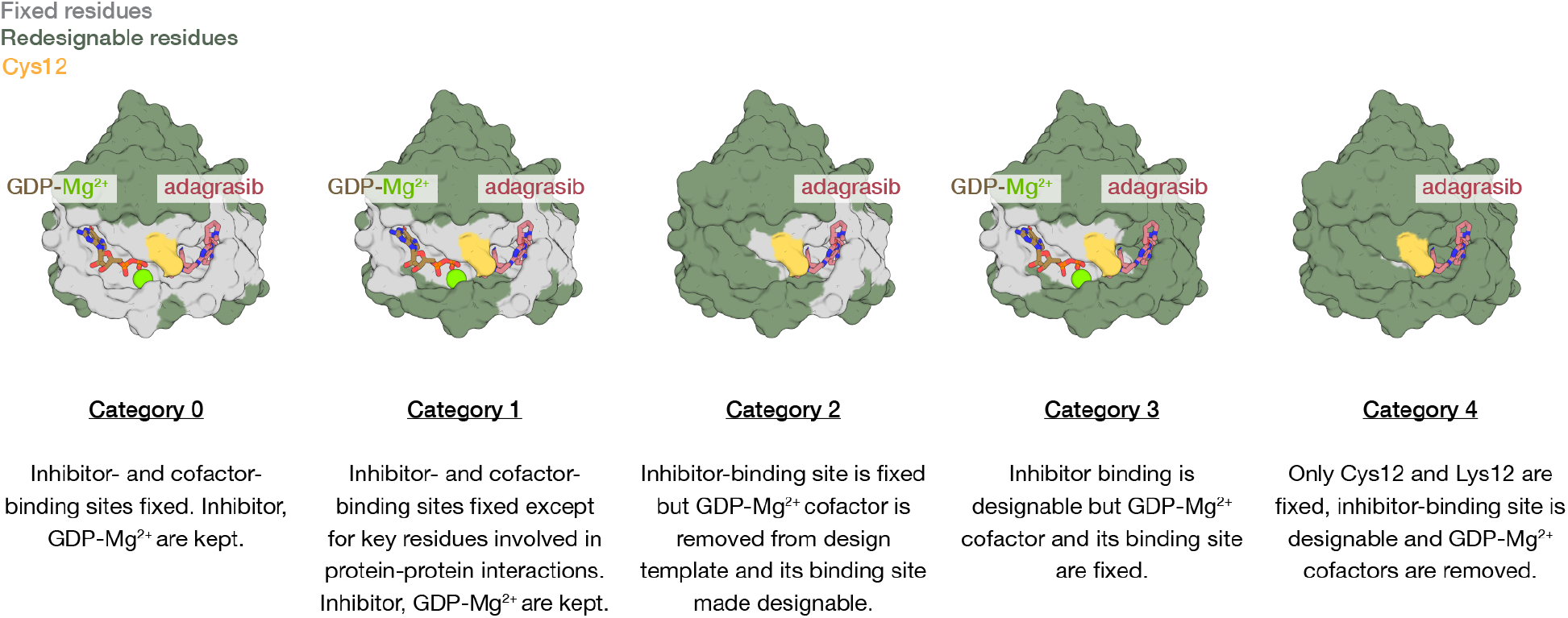
K-Ras(G12C) sequence-design categories. Five categories were used with varying amounts of fixed residues. Category 0 fixes residues within 4.8 Å heavy-atom distance of GDP-Mg^2+^ and the bound inhibitor. The remaining residues are designable. Category 1 starts with the same fixed residues as category 0 but makes designable a few additional residues nearby the ligand and cofactor that are involved in protein–protein interactions. Categories 2 and 3 allow residues contacting the cofactor or inhibitor binding sites to be designable, respectively. Category 4 allows design of all residues except Cys12 and Lys16. Note that most of the protein core (not visible here) was also included as designable residues.

**Figure S4.**
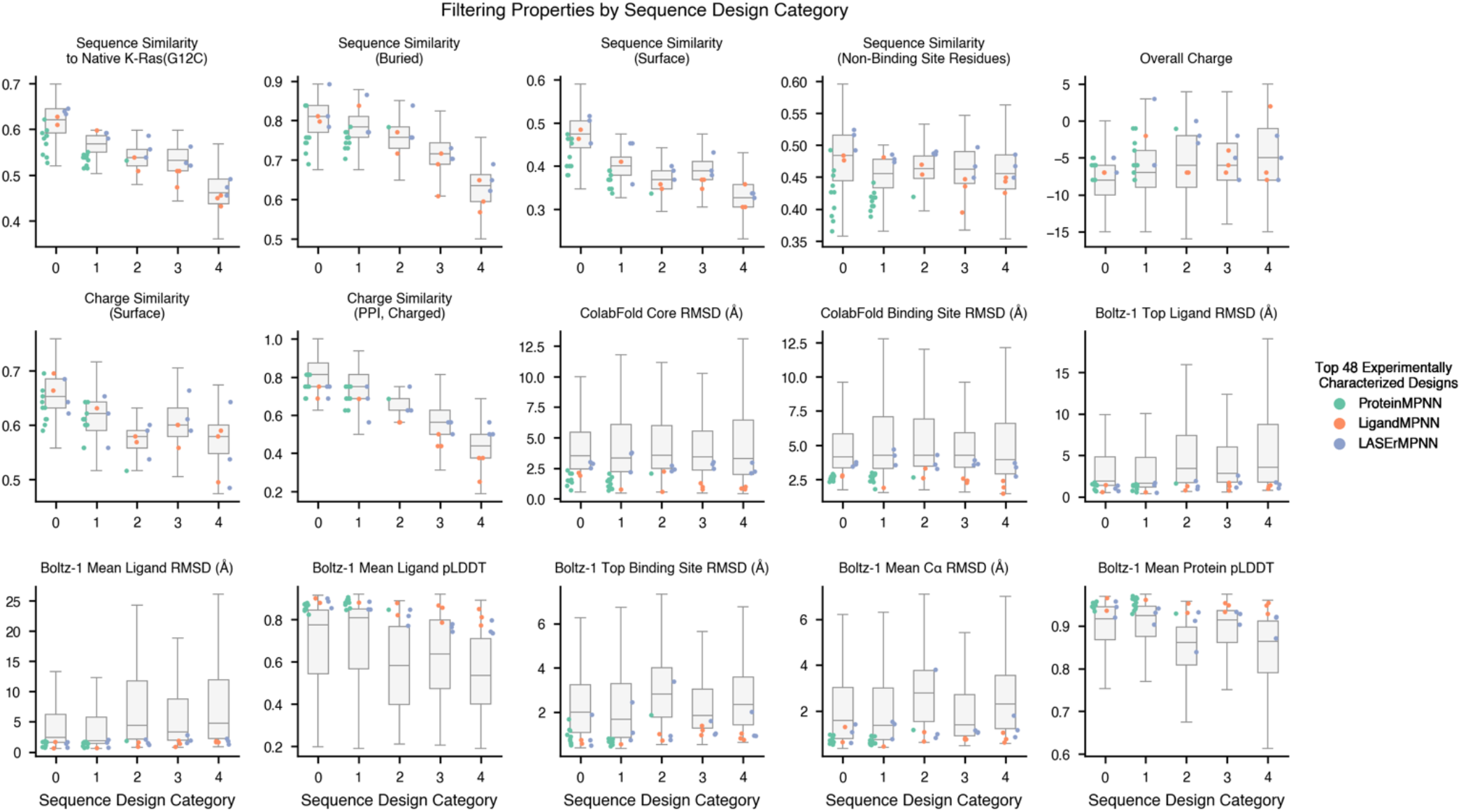
Computational design metrics. Box plots for each sequence design category; The title of each plot is also the y-axis. Colors of each point represent the sequence-design model used for that design (ProteinMPNN, LigandMPNN, or LASErMPNN).

**Figure S5.**
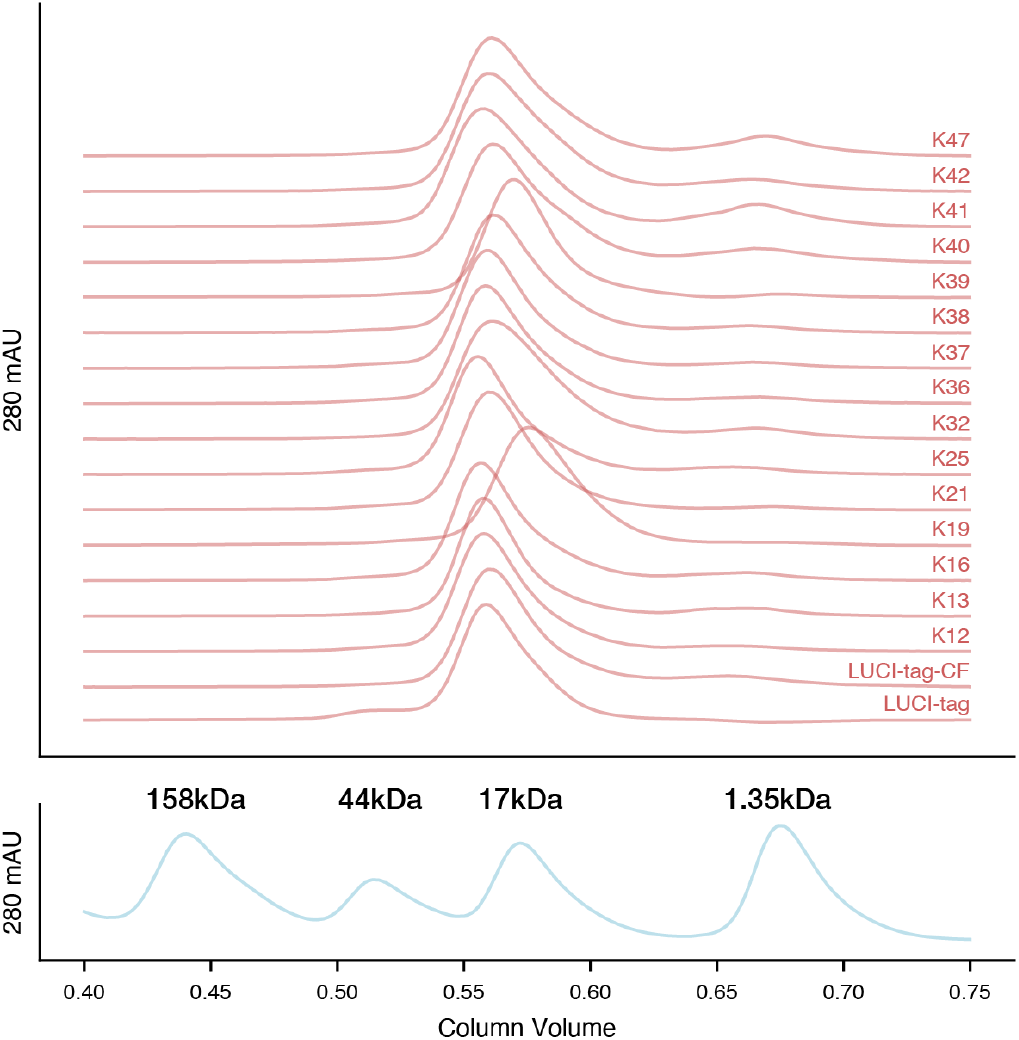
Size-exclusion chromatography traces for each tested design. All tested designs, including LUCI-tag and LUCI-tag–CF, eluted at the expected column volume of a monomeric design. Reference standards of known molecular weight are shown below.

**Figure S6.**
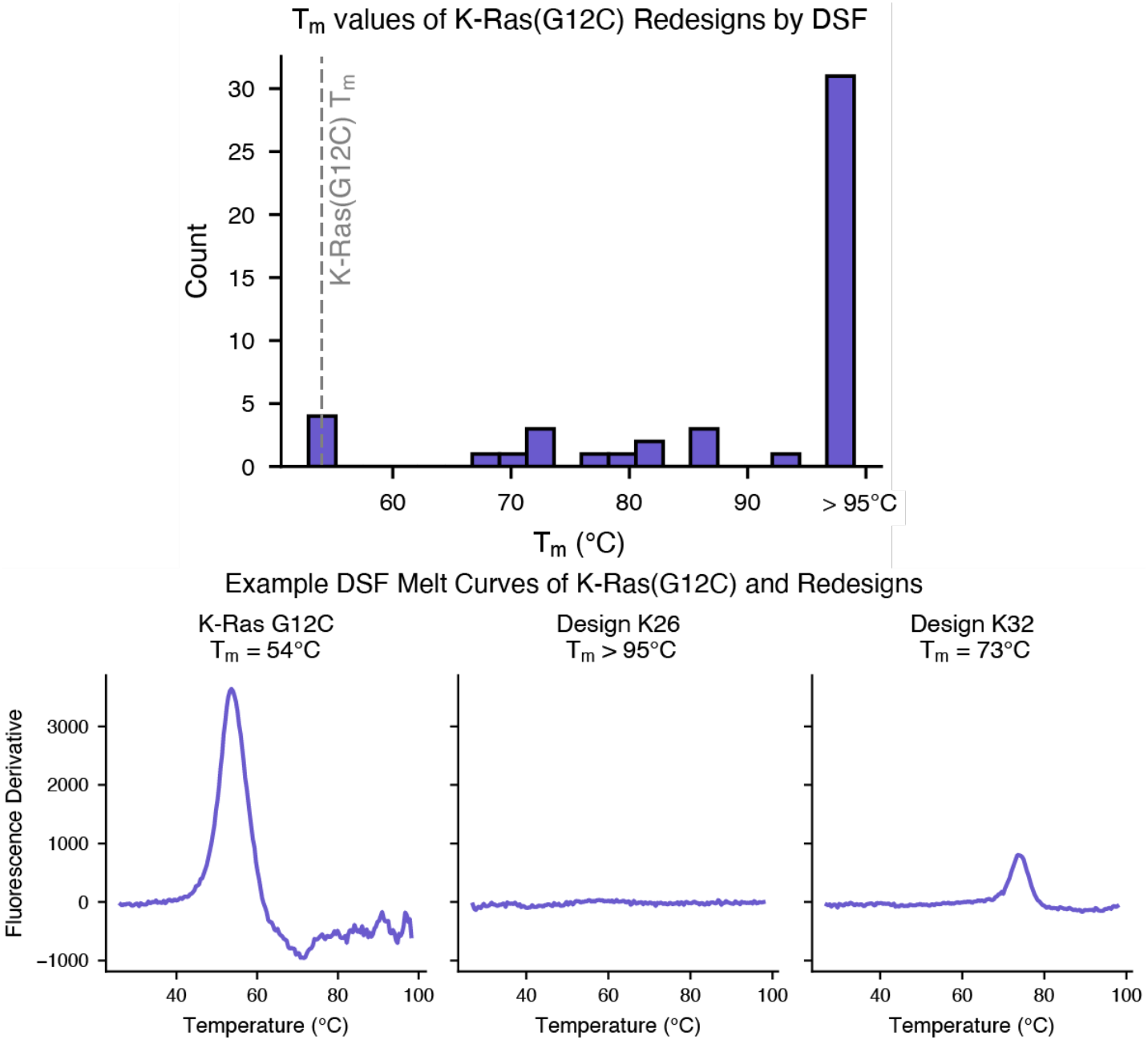
Differential scanning fluorimetry (DSF) of designs. Top, histogram of T_m_ values determined by DSF. All designs that did not unfold at the highest temperature were assigned a T_m_ of 95°C. Gray dashed line indicated the T_m_ of K-Ras(G12C) (54°C) in “protein buffer”. Bottom, DSF curves for K-Ras(G12C) and two representative designs.

**Figure S7.**
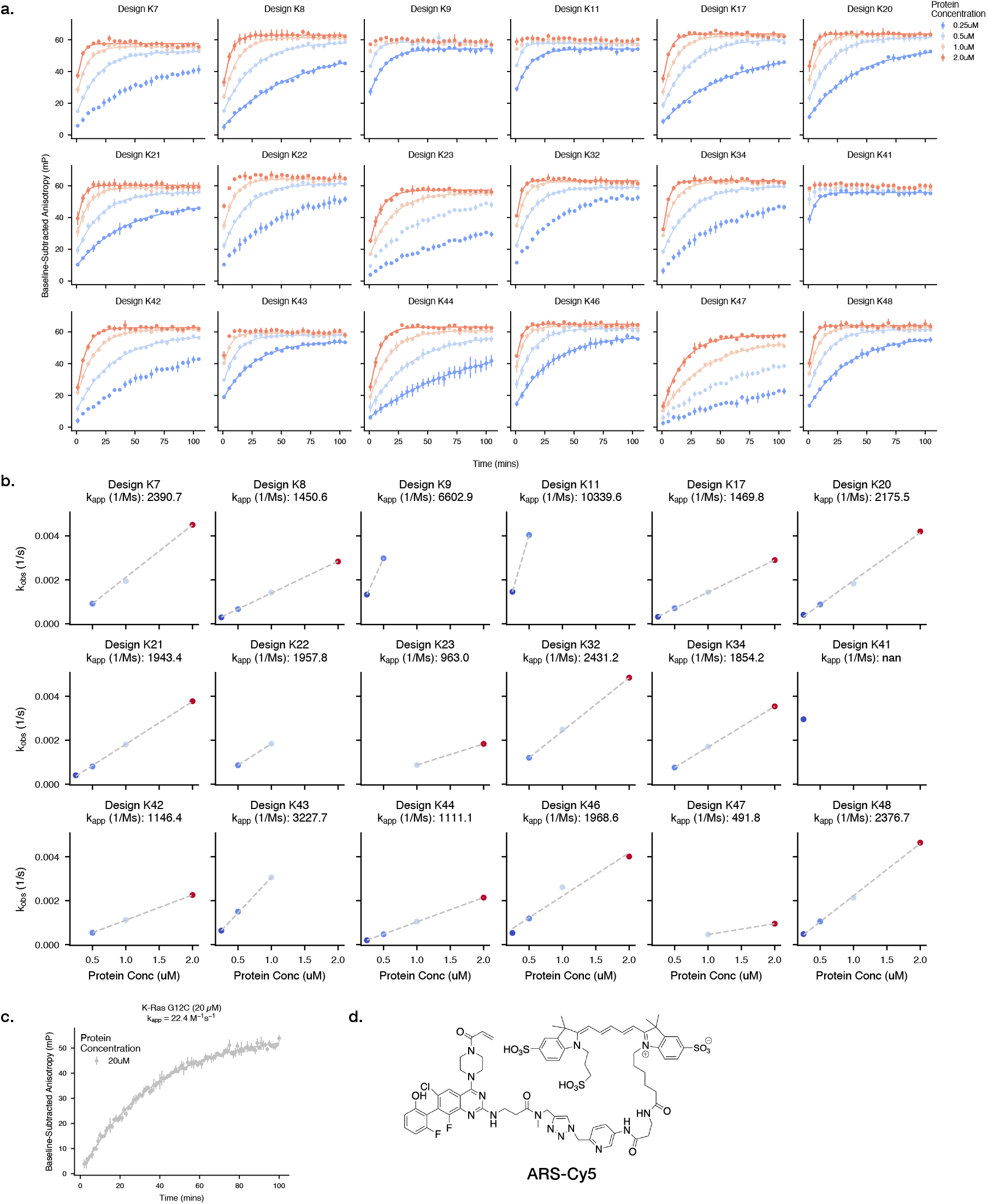
Time-resolved fluorescence anisotropy of working designs with ARS-Cy5. ARS-1323-Cy5 was used to determine labeling kinetics for the top 18 designs. **a,** Kinetics of probe association (50 nM probe) were measured at various protein concentrations. Single-exponential (pseudo-first-order) kinetics were used to fit individual kinetic traces. Note that of the 48 total designs, these 18 are the designs that were monomeric and showed probe labeling fast enough to quantify with this assay. **b,** A linear fit of k_obs_ vs [protein] was performed to obtain an estimate of the 2^nd^ order rate constant k_app_ (see Methods). **c,** K-Ras(G12C) kinetics were estimated at 20 µM protein only. Experiments were performed at protein buffer titrated to pH 8. **d,** structure of the ARS-Cy5 probe. Error bars show standard deviation across 2 (**a**) or 3 (**c**) technical replicates.

**Figure S8.**
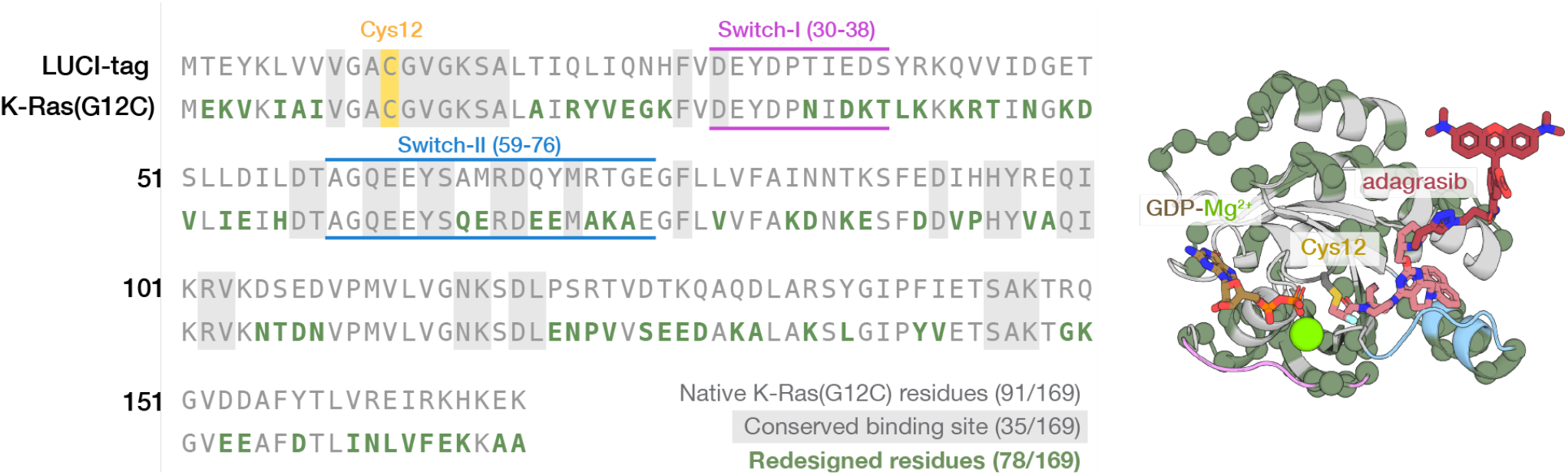
Sequence alignment of LUCI-tag compared to K-Ras(G12C). Residues that were designable but remained unchanged in LUCI-tag are shown in grey; those with a gray shaded background were fixed to their native identity during design. Residues with altered identities are shown in green. Switch-I and Switch-II regions are highlighted in magenta and blue, respectively.

**Figure S9.**
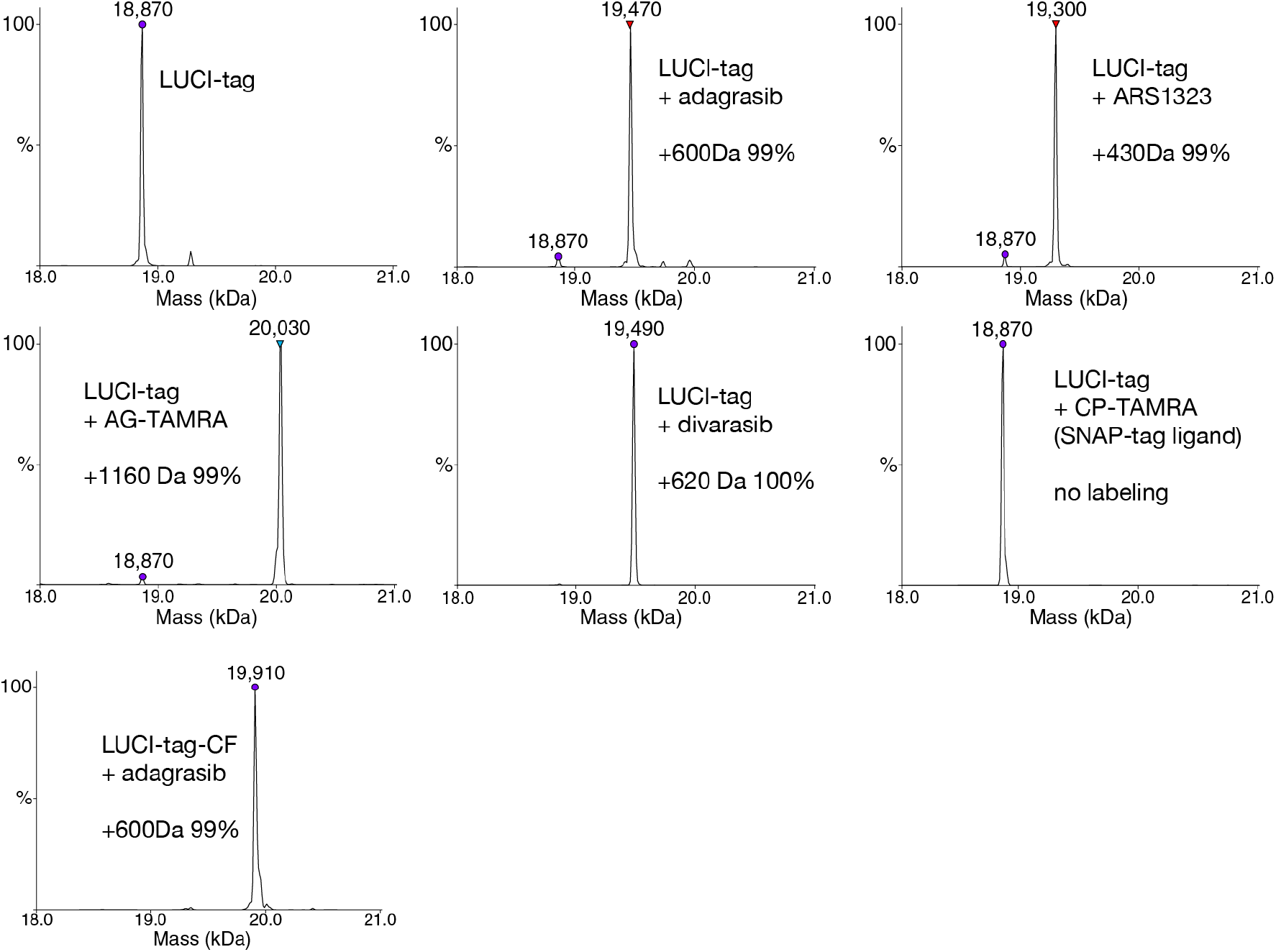
Intact mass spectrometry showing LUCI-tag is covalently labeled with various K-Ras(G12C) inhibitors. Purified LUCI-tag or LUCI-tag–CF was labeled with the indicated probe and analyzed by intact protein mass spectrometry at a ratio of 1:2 protein:ligand. LUCI-tag labeled with divarasib, ARS-1323, or LUCI-tag–CF labeled with adagrasib each showed a single mass shift (+620 Da, +430 Da, and +600 Da, respectively) consistent with complete (100%) covalent adduct formation and no unlabeled protein remaining. LUCI-tag incubated with CP-TAMRA (a SNAP-tag ligand) showed no mass shift, confirming LUCI-tag is orthogonal to the SNAP ligand. All experiments used 10 µM protein in “protein buffer” and were quenched after 10 min with formic acid, with the exception of experiments with ARS-1323 and SNAP-tag ligand, which were quenched at 60 min.

**Figure S10.**
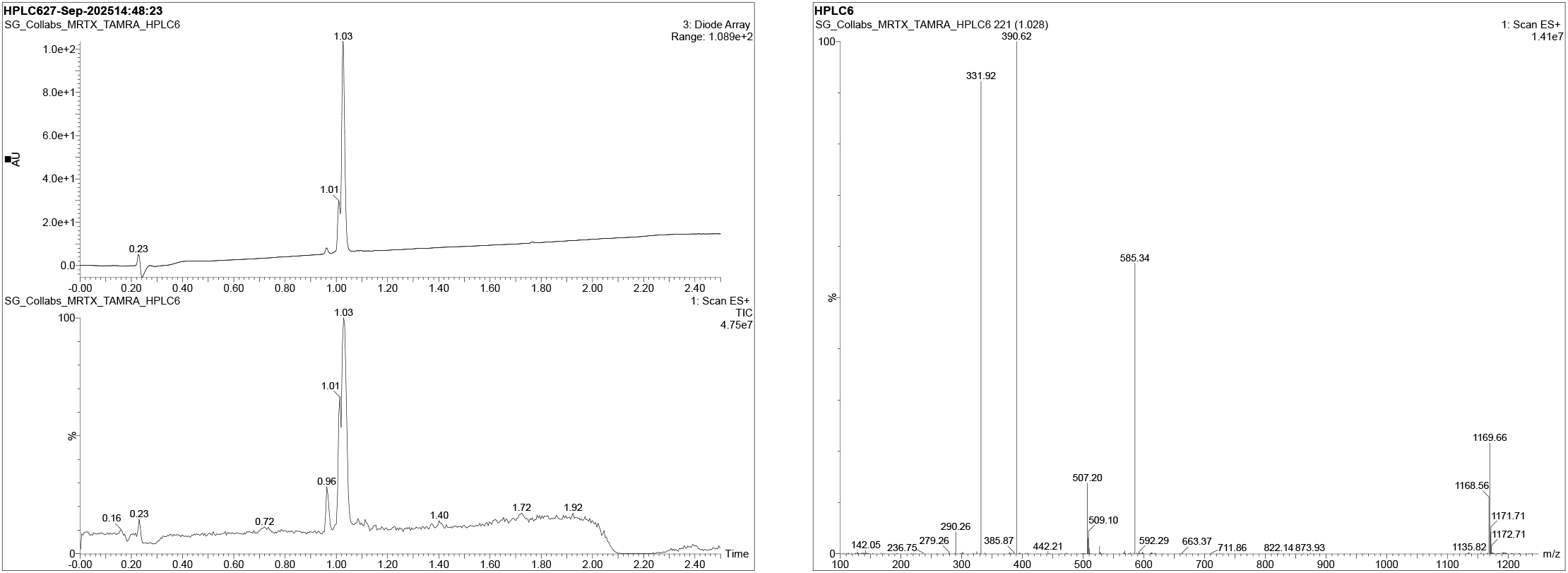
LC–UV and LC–MS characterization of purified AG–TAMRA. Left: Total Ion Chromatogram (TIC) and Analytical LC-UV trace showing single product peak at t_R_ ≈ 1.03 min. Right: LC-MS trace of product peak. Calc’d [M+H] ^+^: 1167.50. Found [M+H]^+^: 1168.56.

**Figure S11.**
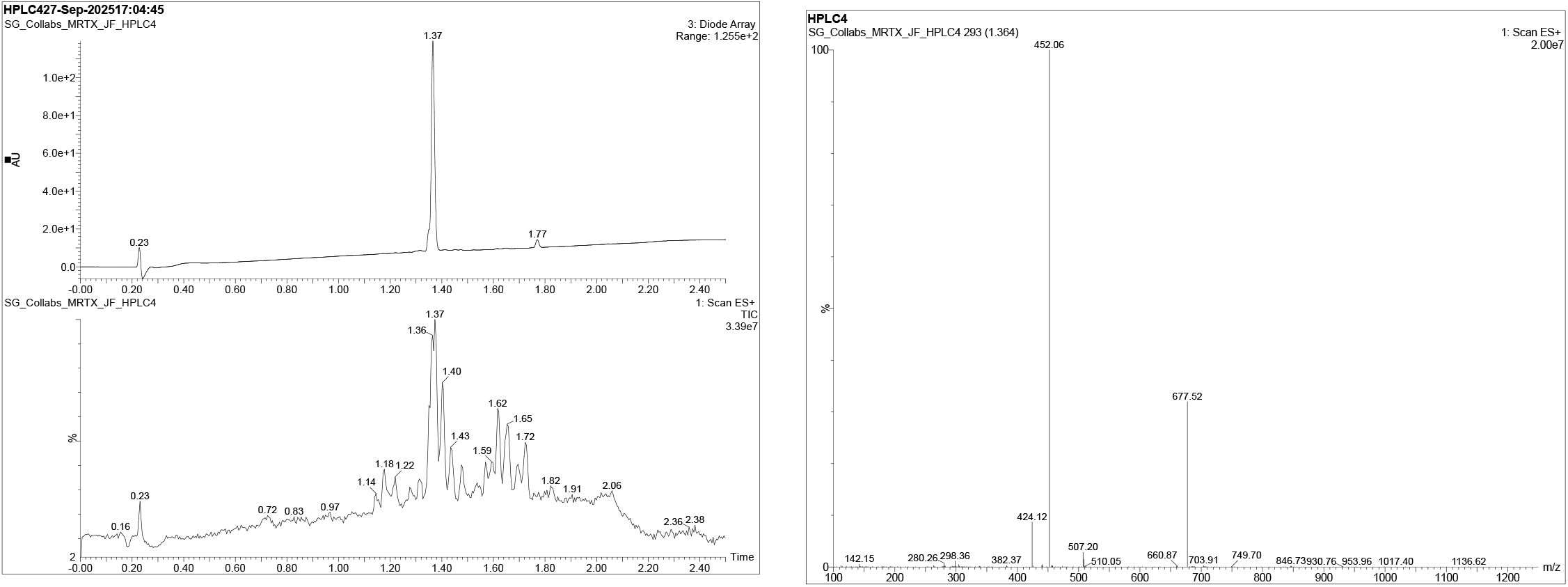
LC–UV and LC–MS characterization of purified AG–JF646. Left: Total Ion Chromatogram (TIC) and Analytical LC-UV trace showing single product peak at t_R_ ≈ 1.37 min. Right: LC-MS trace of product peak Calc’d [M+H] ^+^: 1353.59. Found [M+2H]^2+^: 677.52.

**Figure S12.**
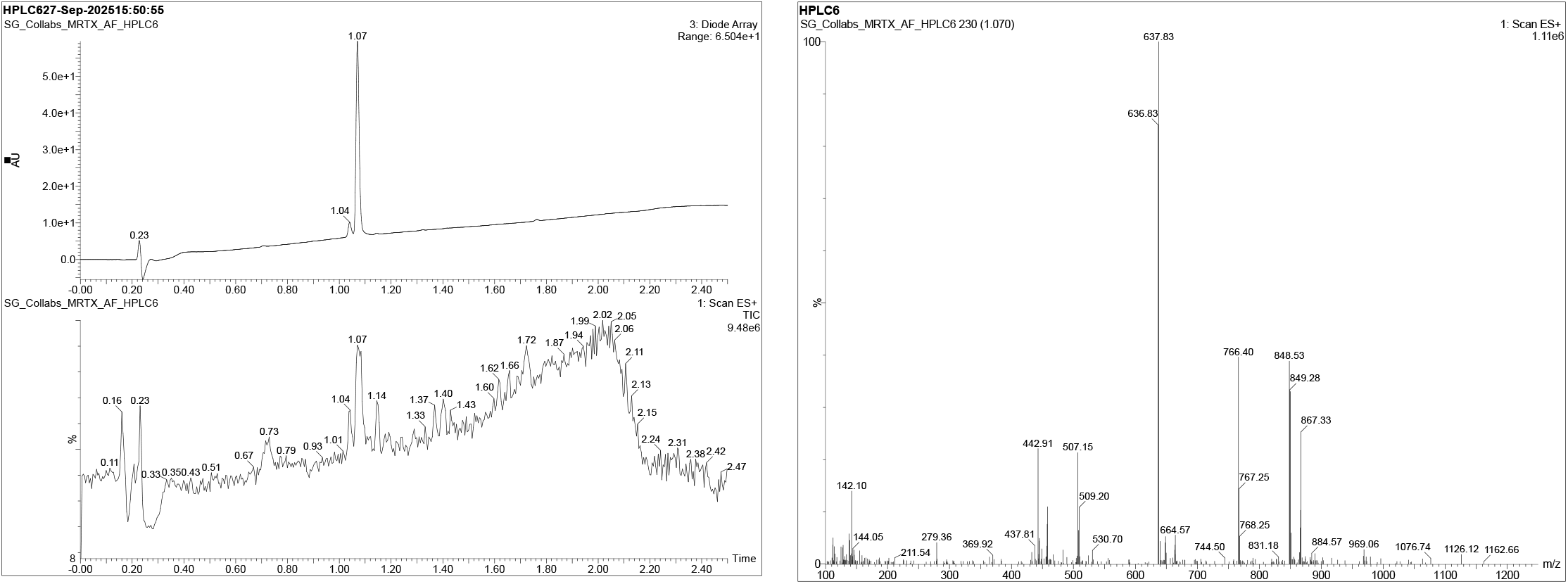
LC–UV and LC–MS characterization of purified AG–AF488. Left: Total Ion Chromatogram (TIC) and Analytical LC-UV trace showing single product peak at t_R_ ≈ 1.07 min. Right: LC-MS trace of product peak. Calc’d [M+H] ^+^: 1273.34. Found [M+2H]^2+^: 636.83.

**Figure S13.**
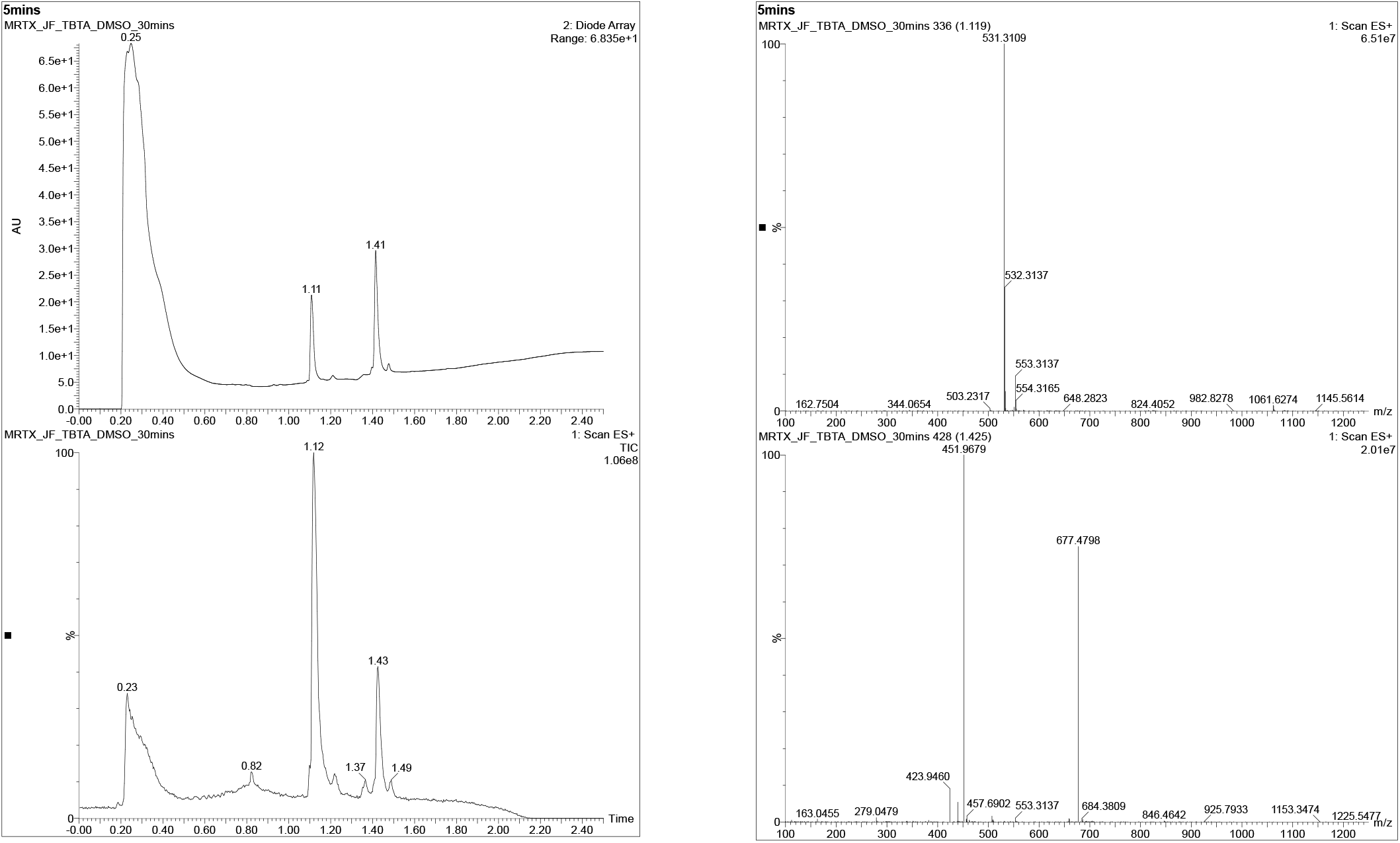
LC–UV and LC–MS characterization of AG–JF646 "Direct to Biology" synthesis with TBTA. Top: Analytical LC–UV trace of the reaction mixture after 30 min. Two major peaks are observed: TBTA (t_R_ ≈ 1.11 min) and the AG–JF646 product (t_R_ ≈ 1.41 min), with the early broad signal corresponding to the solvent front. Bottom: ESI+ mass spectrum of the product peak showing the expected multiply charged ions for AG–JF646, with dominant peaks at m/z 677 ((M+2)/2) and 452 ((M+3)/3), consistent with the calculated molecular mass of 1353 Da.

**Figure S14.**
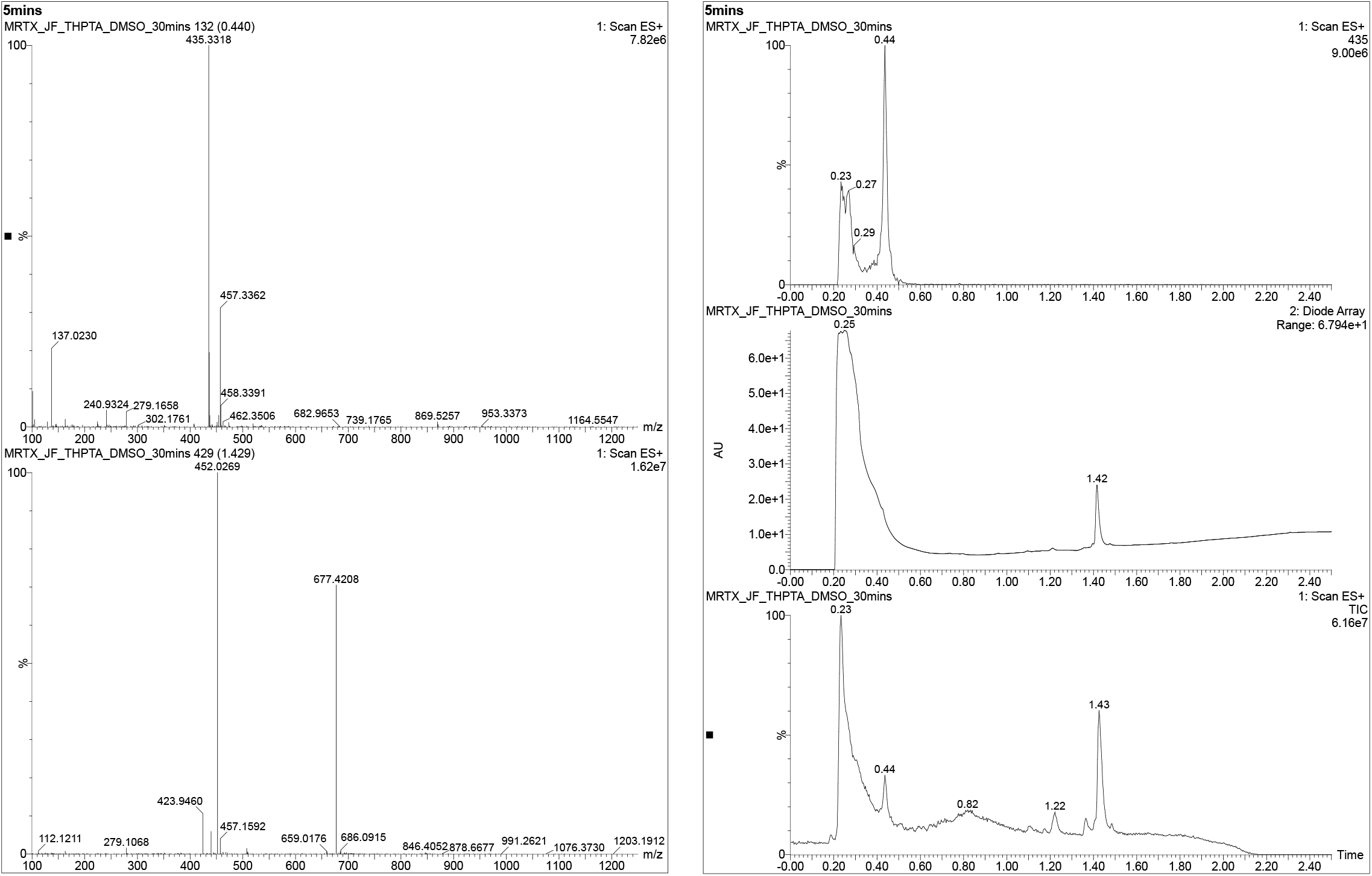
LC–UV and LC–MS characterization of AG–JF646 "Direct to Biology" synthesis with THPTA. Top: Analytical LC–UV trace of the reaction mixture after 30 min. Only one peak corresponding to the AG–JF646 product (t_R_ ≈ 1.42 min) is seen with a similar retention time to the previous figure; THPTA elutes with the solvent front. Bottom: Similar to above, ESI+ mass spectrum of the product peak shows the expected multiply charged ions for AG–JF646, with dominant peaks at m/z 677 ((M+2)/2) and 452 ((M+3)/3), consistent with the calculated molecular mass of 1353 Da.

**Figure S15.**
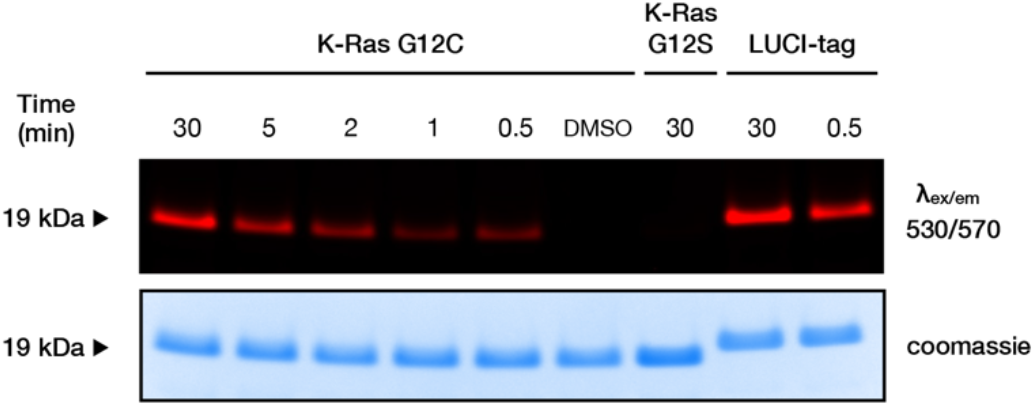
Staining time-course of purified K-Ras(G12C) in gel. Purified K-Ras(G12C) (10 µM) was mixed with AG–TAMRA (0.5 µM) and quenched at various timepoints, after which aliquots were analyzed via SDS-PAGE. Unlike LUCI-tag, where covalent staining is complete at 30 s (right), K-Ras(G12C) requires > 30 min for complete labeling. DMSO control (no AG–TAMRA) showed no labeling. K-Ras(G12S) does not show any covalent labeling due to loss of Cys12. Labeling was done in protein buffer.

**Figure S16.**
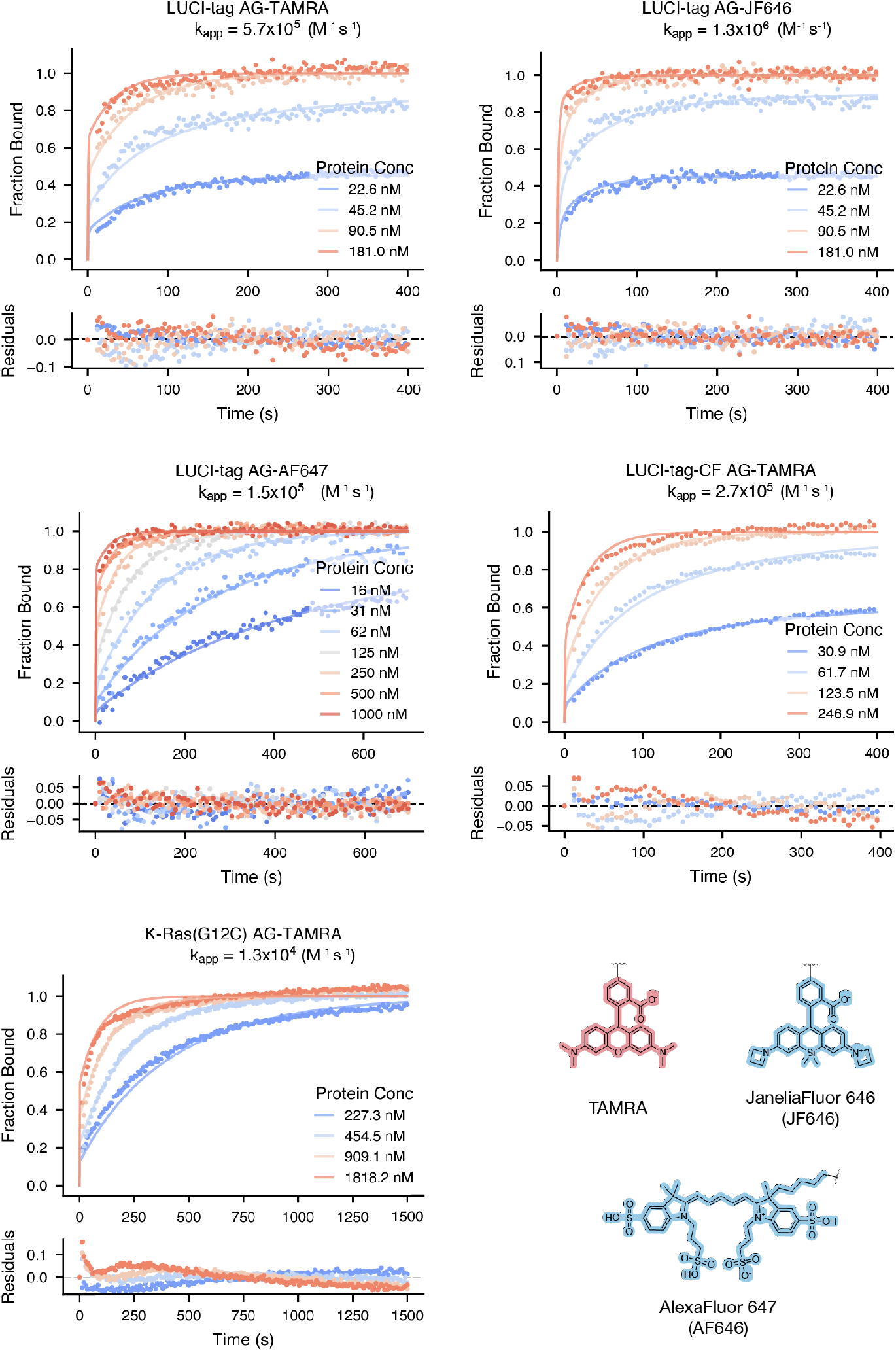
Time-resolved fluorescence anisotropy of AG bearing different payloads. [probe] = 10nM for AG-AF647 and [probe] = 50 nM for AG-TAMRA and AG-JF646. Note the higher protein concentrations used for K-Ras(G12C) due to slower labeling kinetics. Data was fit using a full two-step binding model to obtain estimated second-order rate constant k_app_ values.

**Figure S17.**
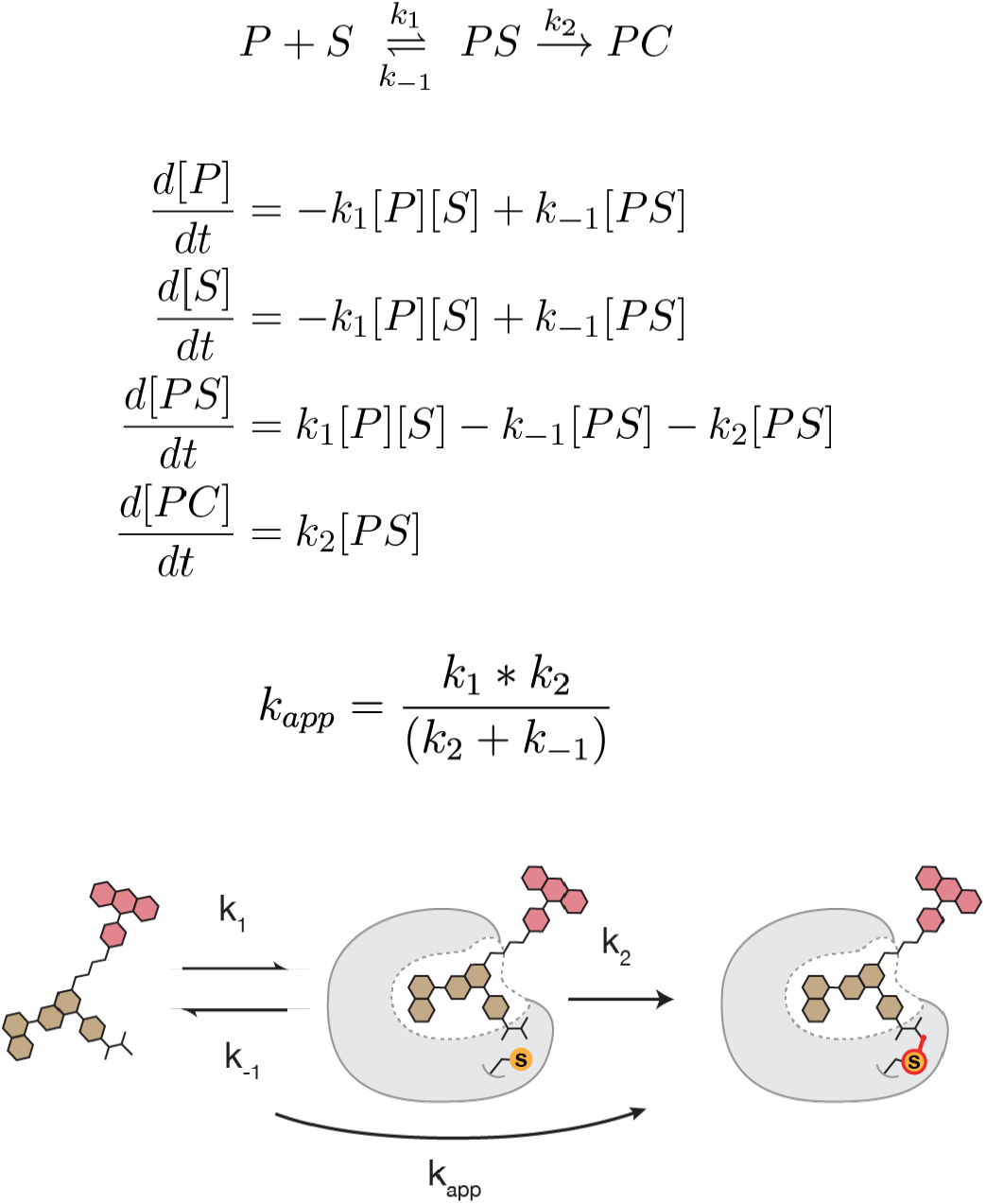
Two-step binding model considered in fitting of fluorescence anisotropy. System of differential equations based on two-step binding model described at the top. The 2nd order rate constant k_app_ (M^-1^ s^-1^) describes the overall rate of the reaction, including both the noncovalent and covalent steps.

**Figure S18.**
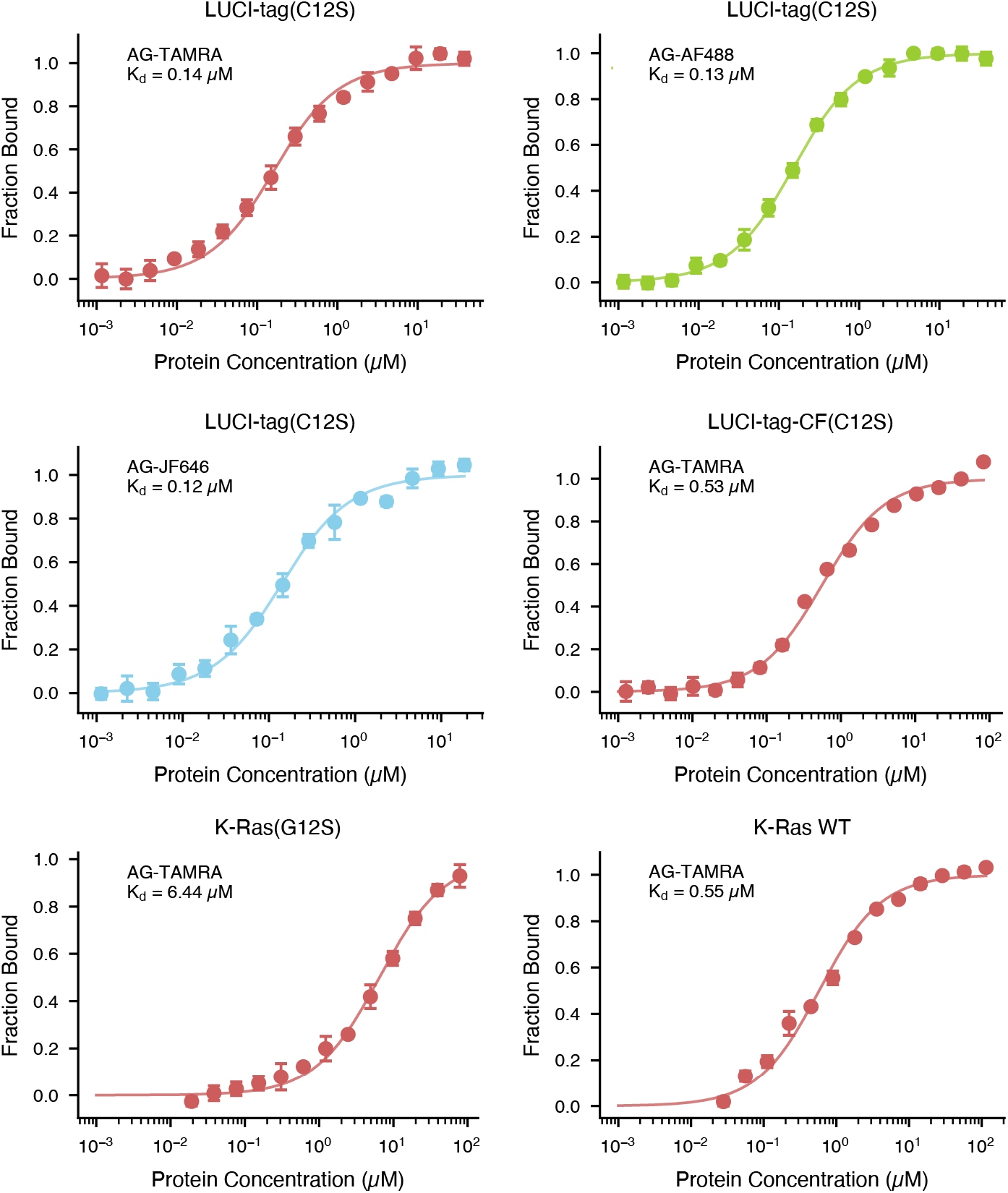
Binding curves of noncovalent variants LUCI-tag(C12S), LUCI-tag–CF(C12S), K-Ras(G12S) and K-Ras WT. Gly12 and Cys12 were mutated to Ser12 to measure noncovalent binding affinity. Protein concentration was titrated while fixing the probe-dye ligand at 50 nM. Similar to its covalent kinetics, LUCI-tag displays similar noncovalent binding affinities across payloads. Note that these proteins were not nucleotide exchanged and measurements were performed directly from purified protein after *E. coli* expression. Lines are numerical fits of a 1:1 binding model. Inset gives the AG–payload used in each experiment. Error bars show standard deviation across 3 technical replicates.

**Figure S19.**
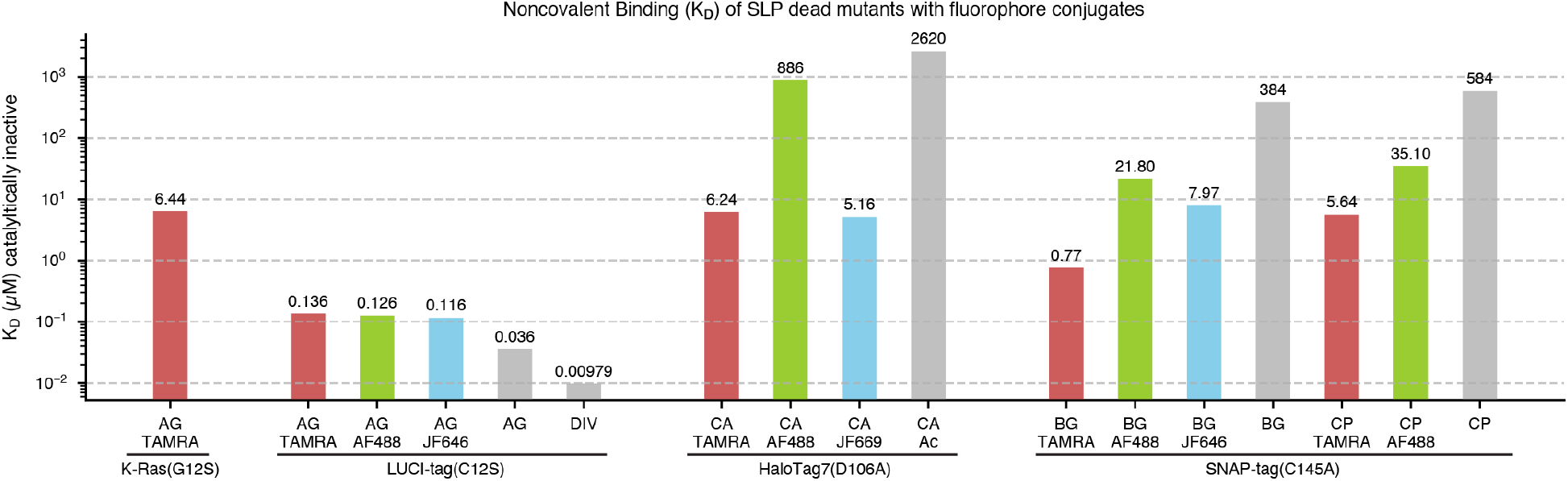
Non-covalent binding affinities of LUCI-tag(C12S), K-Ras(G12S), HaloTag7(D106A), and SNAP-tag(C145A) with probe–fluorophore conjugates. Binding affinities of mutant HaloTag7 and SNAP-tag with their probes are taken from ref. ^12^. Parentheses indicate the mutation that renders the SLP incapable of covalent reactivity with the probe. AG = adagrasib, DIV = divarasib, CA = chloroalkane, CA-Ac = chloroalkane-acetate, BG = benzylguanine, CP = chloropyrimidine.

**Figure S20.**
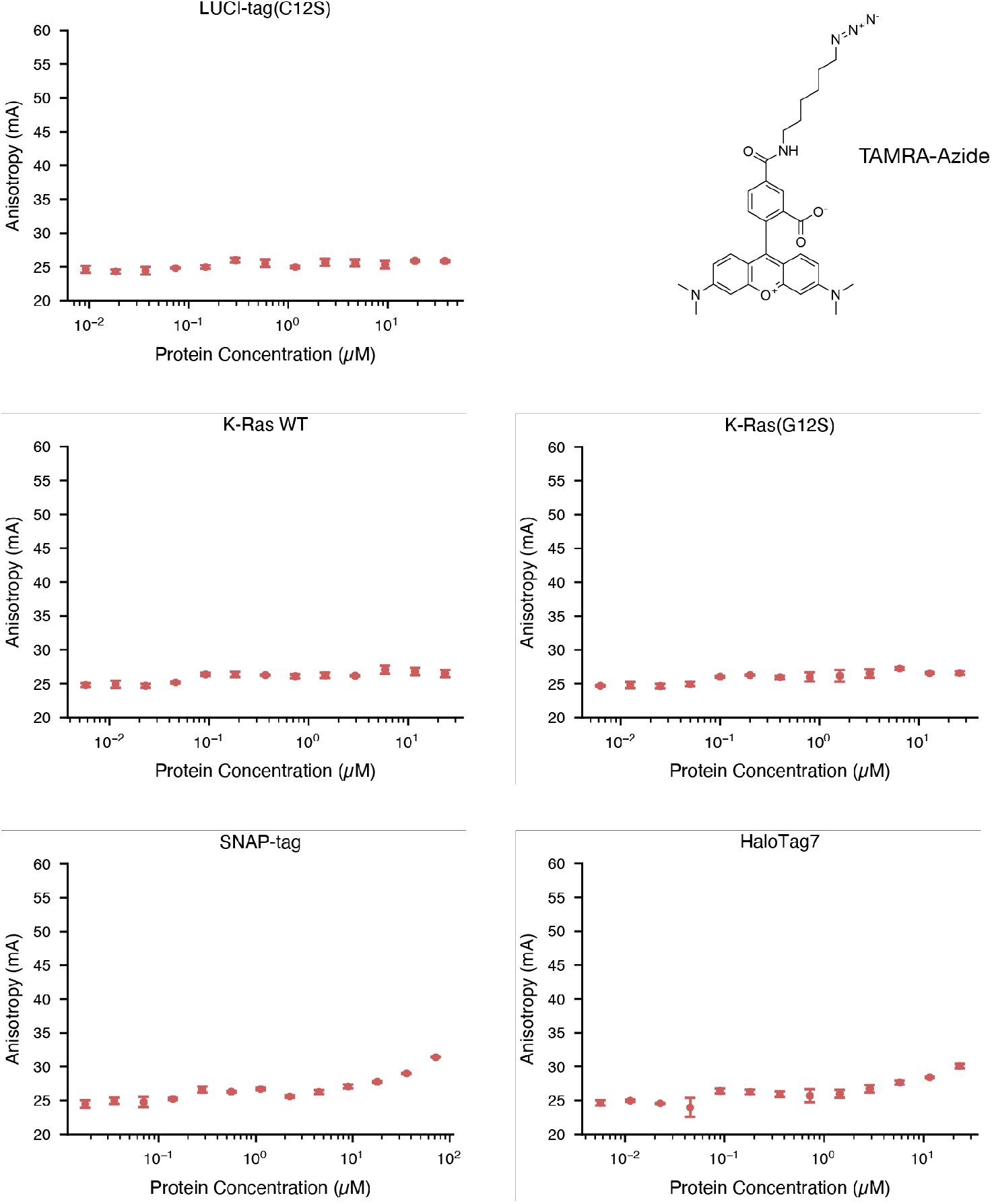
Fluorescence anisotropy of TAMRA-azide dye with various proteins. [TAMRA-azide] = 50 nM. No dye binding is observed for LUCI-tag(C12S), K-Ras(G12S), or wildtype K-Ras ΔHVR. The dye starts to show some binding to HaloTag7 and SNAP-tag at high (> 10 µM) protein concentrations. Error bars show standard deviation across 3 technical replicates.

**Figure S21.**
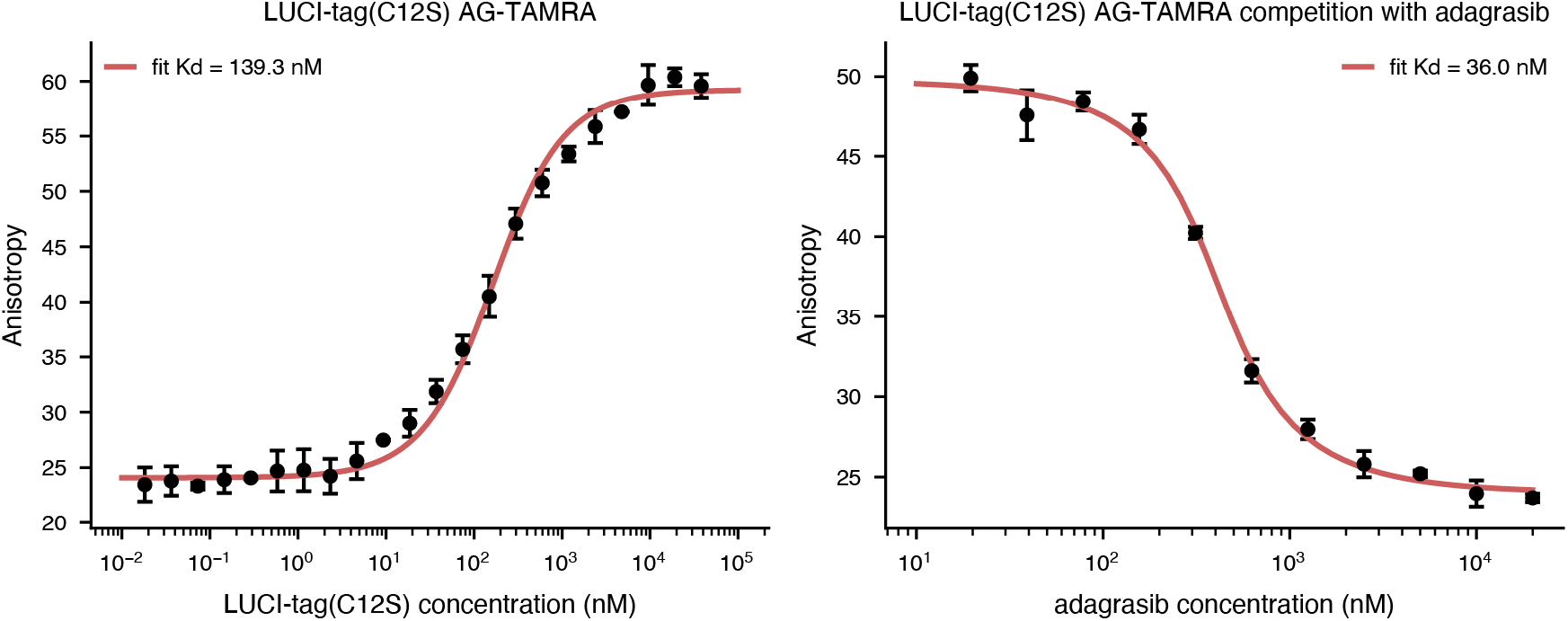
Binding affinity of AG to LUCI-tag(C12S) measured via competition fluorescence anisotropy. A binding titration of LUCI-tag with fixed AG–TAMRA (50 nM) is shown (left). For a fixed concentration of both LUCI-tag (450 nM) and AG–TAMRA (50 nM), we titrated unlabeled AG as a competitor (right). The dissociation constant (K_d_) of LUCI-tag with AG–TAMRA (via direct binding) was fit and then fixed at this value to fit the K_d_ of LUCI-tag with AG (via competitive binding). Error bars denote standard deviation across 3 technical replicates.

**Figure S22.**
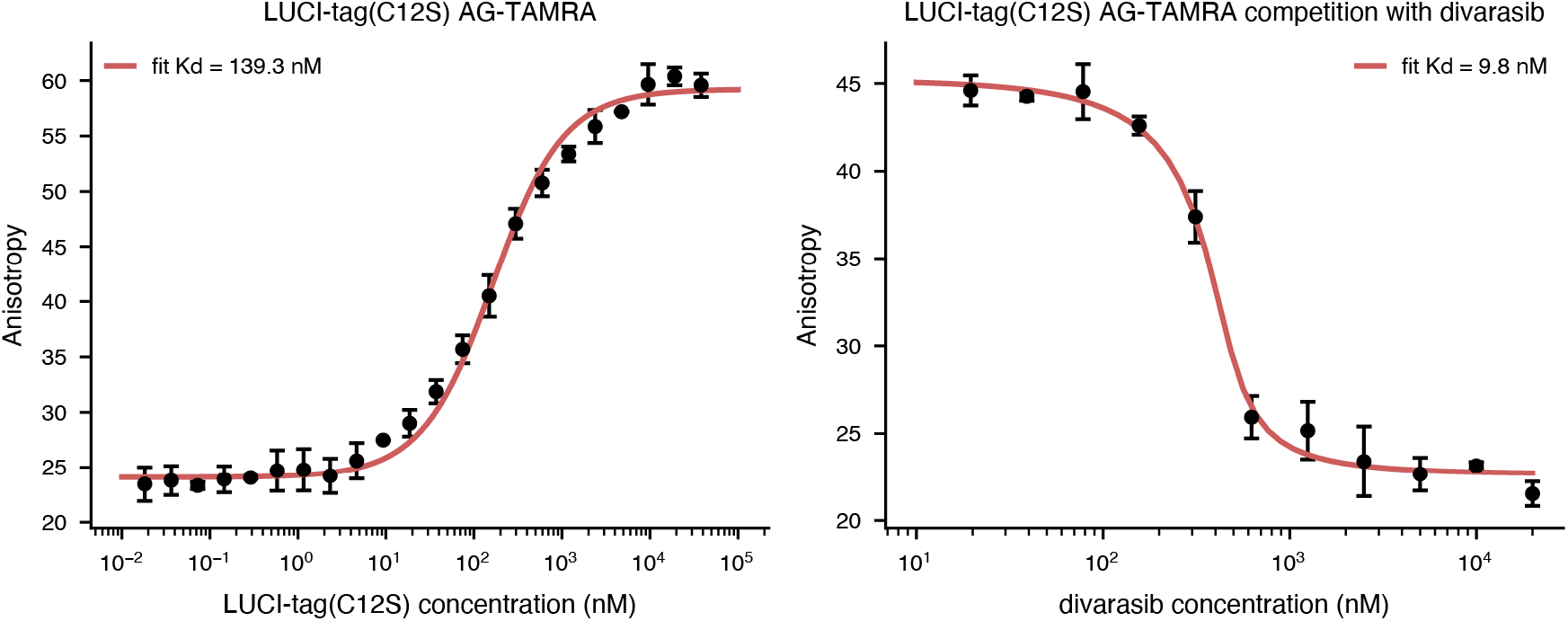
Binding affinity of divarasib to LUCI-tag(C12S) measured via competition fluorescence anisotropy. We performed a titration of LUCI-tag with fixed AG–TAMRA (50 nM). For a fixed concentration of both LUCI-tag (450 nM) and AG–TAMRA (50 nM), we titrated unlabeled divarasib as a competitor. The dissociation constant (K_d_) of LUCI-tag with AG–TAMRA (via direct binding) was fit and then fixed at this value to fit the K_d_ of LUCI-tag with divarasib (via competitive binding). Error bars denote standard deviation across 3 technical replicates.

**Figure S23.**
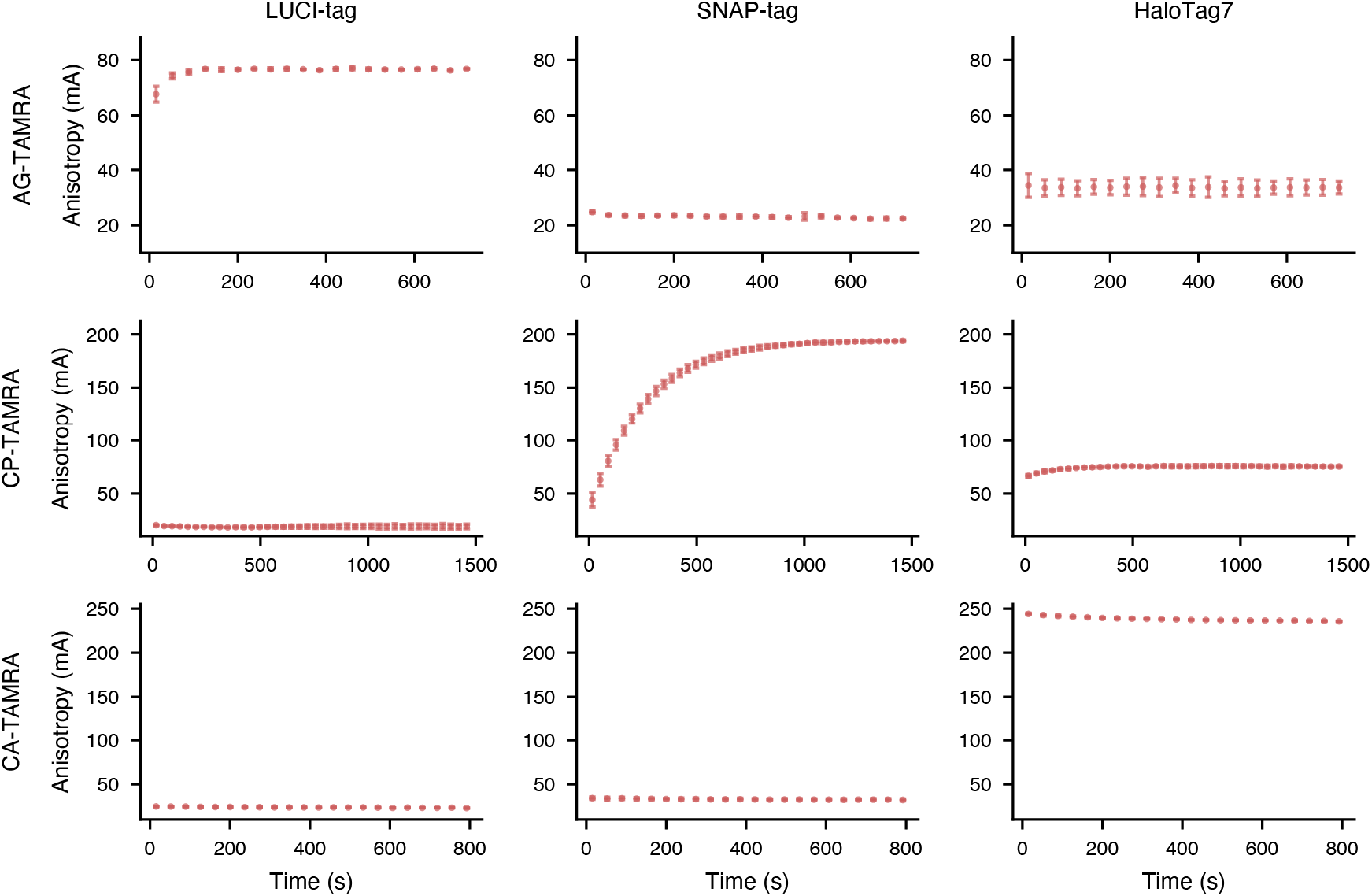
Time-resolved fluorescence anisotropy experiments for all-*vs*-all SLP *vs* probe combinations. LUCI-tag, SNAP-tag, and HaloTag7 (400 nM) were mixed with TAMRA-conjugated probes (50 nM). Each SLP reacts covalently with their respective probes, as expected, and shows minimal off-target reactivity. HaloTag7 may bind noncovalently to AG–TAMRA and CP-TAMRA probes due to its affinity for rhodamine dyes. Error bars show standard deviation across 3 technical replicates.

**Figure S24.**
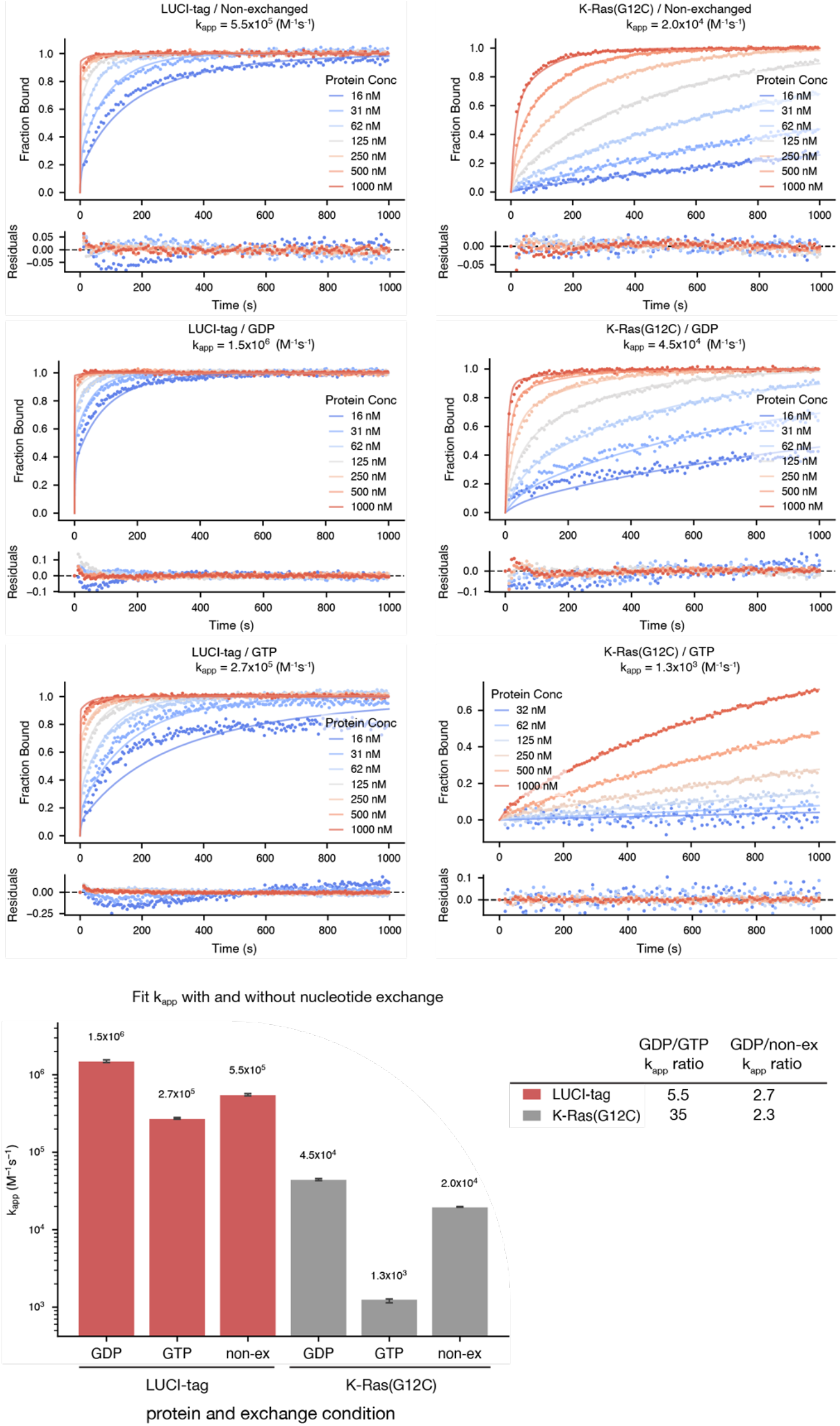
Time-resolved fluorescence anisotropy of AG–AF488 with LUCI-tag and K-Ras(G12C) in various nucleotide-bound forms. LUCI-tag and K-Ras(G12C) were exchanged with GDP, GTP, or non-exchanged prior to labeling with AG–AF488 (10 nM). Fluorescence anisotropy was globally fit (lines) for each dataset using a two-step binding model (see Supp. Fig. S17) to obtain second-order rate constants k_app_. Residuals to the fits are shown below each plot. Error bars show the 95% confidence interval of a bootstrapped error estimation.

**Figure S25.**
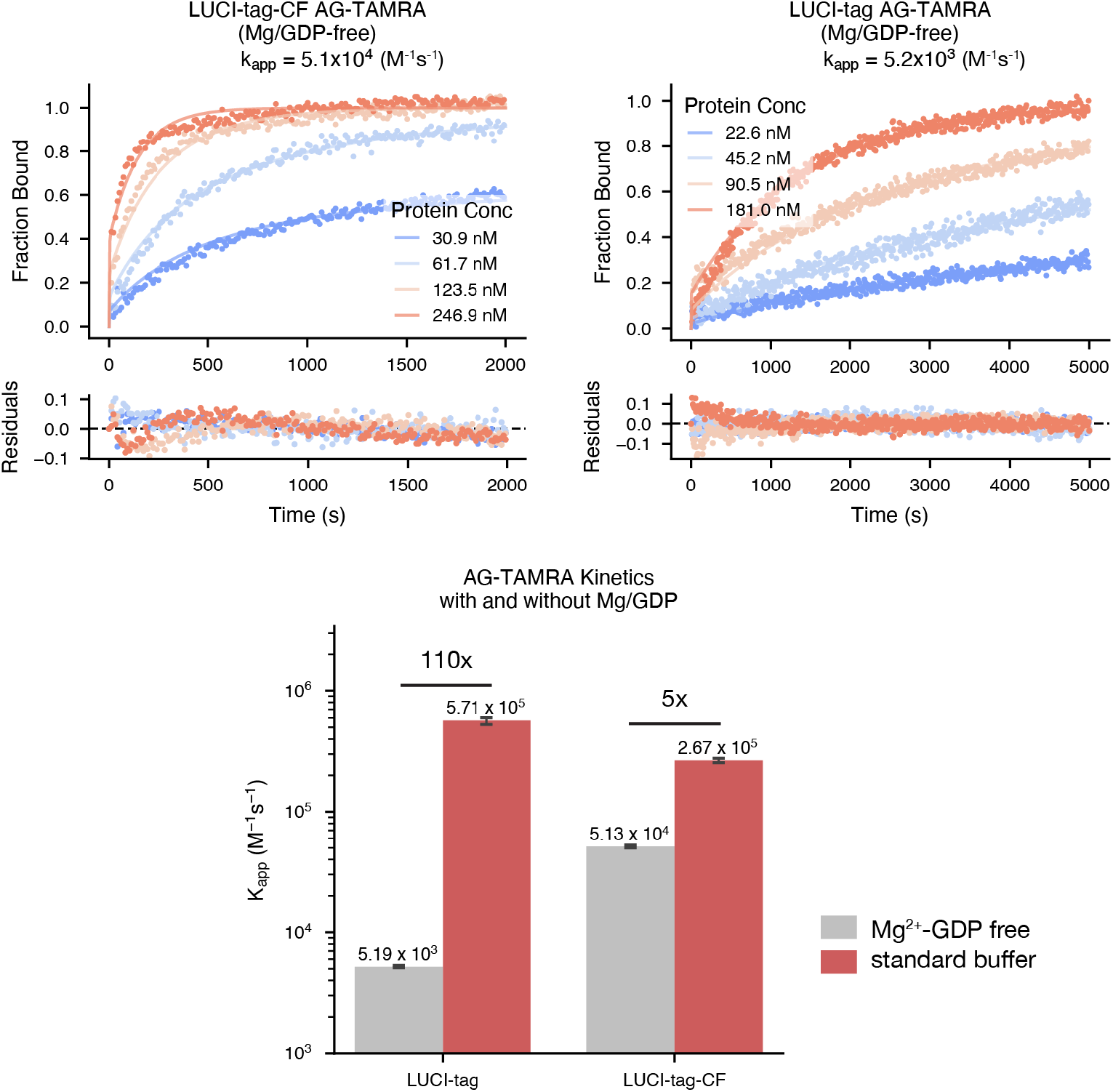
k_app_ of LUCI-tag and LUCI-tag–CF with AG–TAMRA in GDP-Mg^2+^-free buffer. In cofactor-free protein buffer (grey), LUCI-tag–CF shows a much smaller decrease in labeling rate (5x) compared to LUCI-tag (110x). Error bars show the 95% confidence interval of a bootstrapped error estimation (see methods).

**Figure S26.**
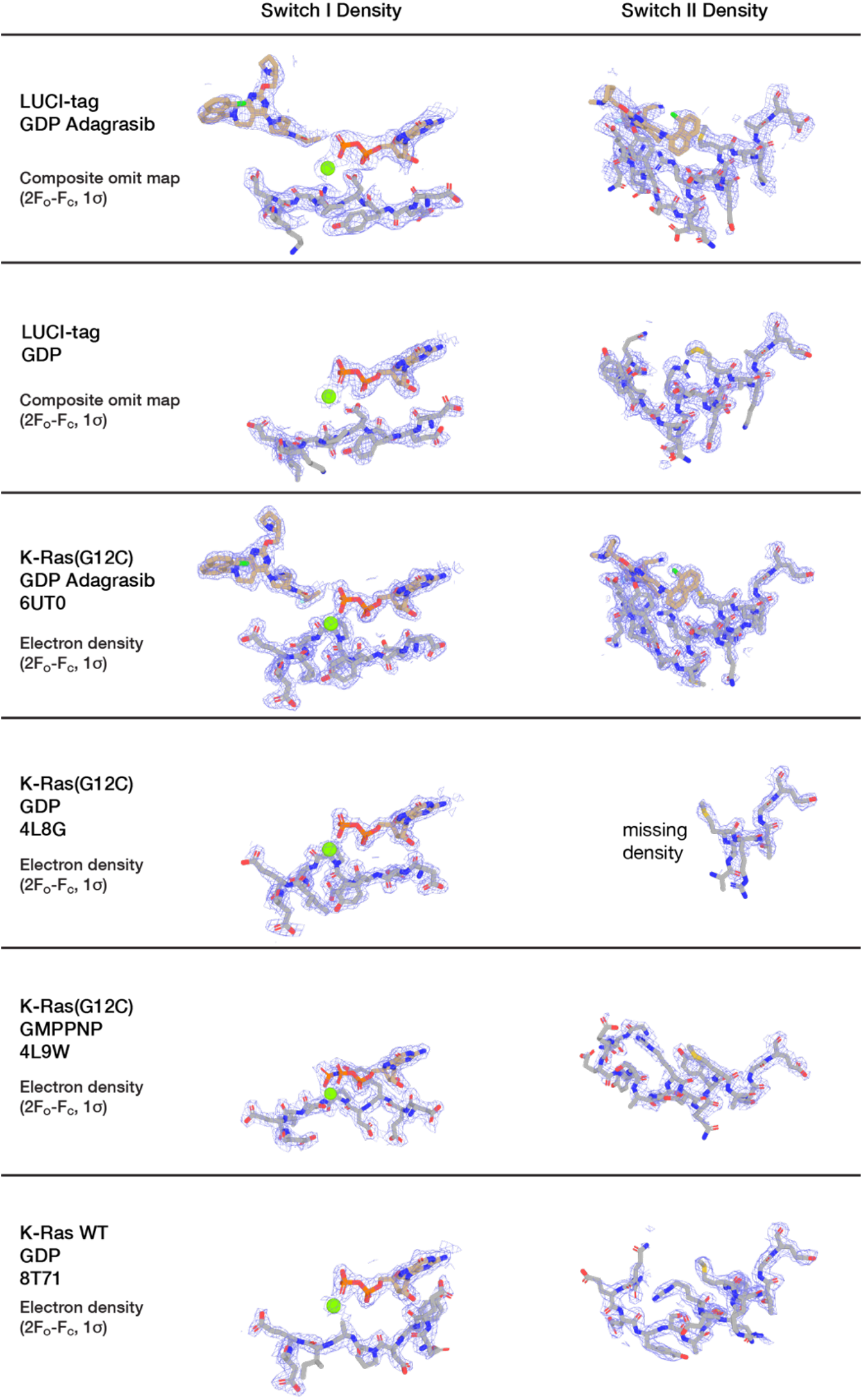
Electron density at Switch I and Switch II in LUCI-tag and related K-Ras structures. Switch I density (left column) and Switch II density (right column) are shown for each structure. Composite omit maps (2Fo–Fc, contoured at 1σ) are shown for AG-bound (GDP-bound) LUCI-tag and inhibitor-free (GDP-bound) LUCI-tag, determined in this work; 2Fo–Fc electron density is shown for the deposited structures of GMPPNP-bound K-Ras(G12C) (PDB 4L9W), AG-bound K-Ras(G12C) (PDB 6UT0), inhibitor-free (GDP-bound) K-Ras(G12C) (PDB 4L8G), and inhibitor-free (GDP-bound) K-Ras WT (PDB 8T71). Regions lacking interpretable density are annotated as missing.

**Figure S27.**
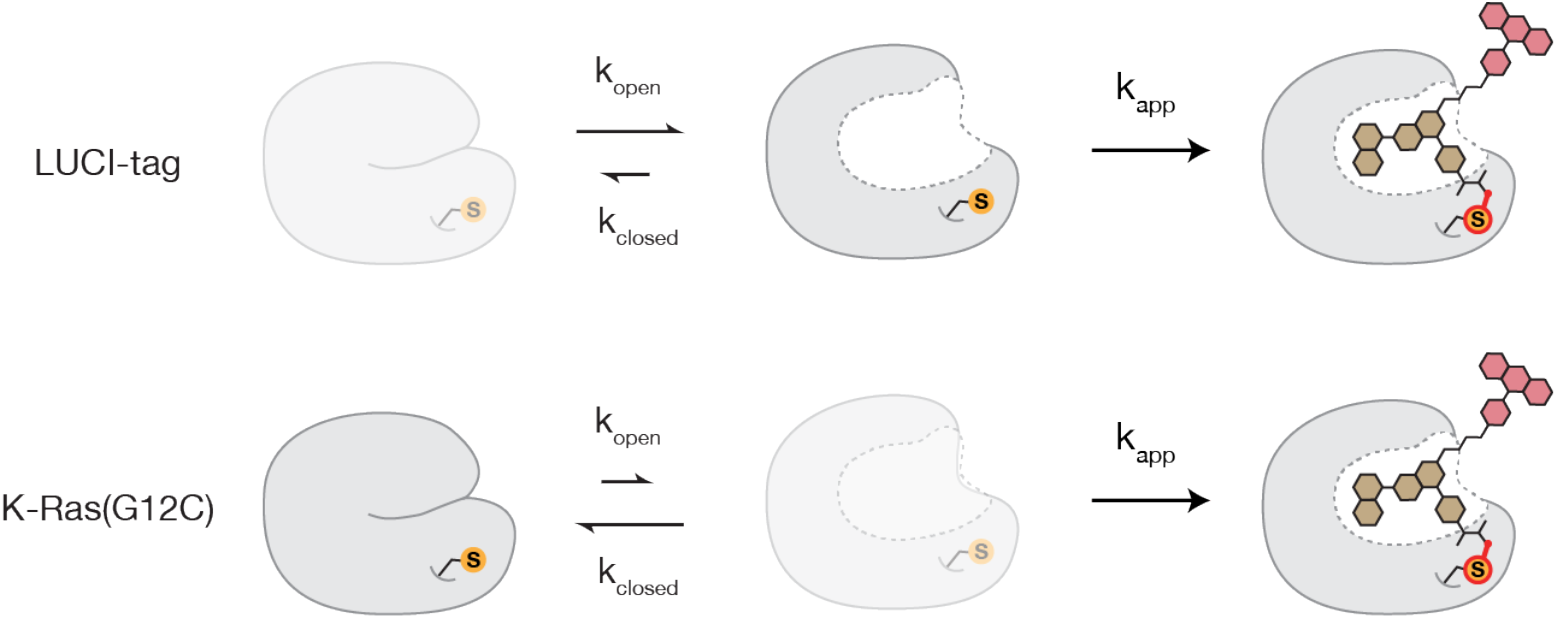
Kinetic scheme of probe binding with additional conformational step (k_closed/open_). LUCI-tag likely preorganizes the drug-free, pocket-open state, such that there is a greater population of binding-competent protein, consistent with increased binding affinity and faster labeling kinetics relative to K-Ras(G12C).

**Figure S28.**
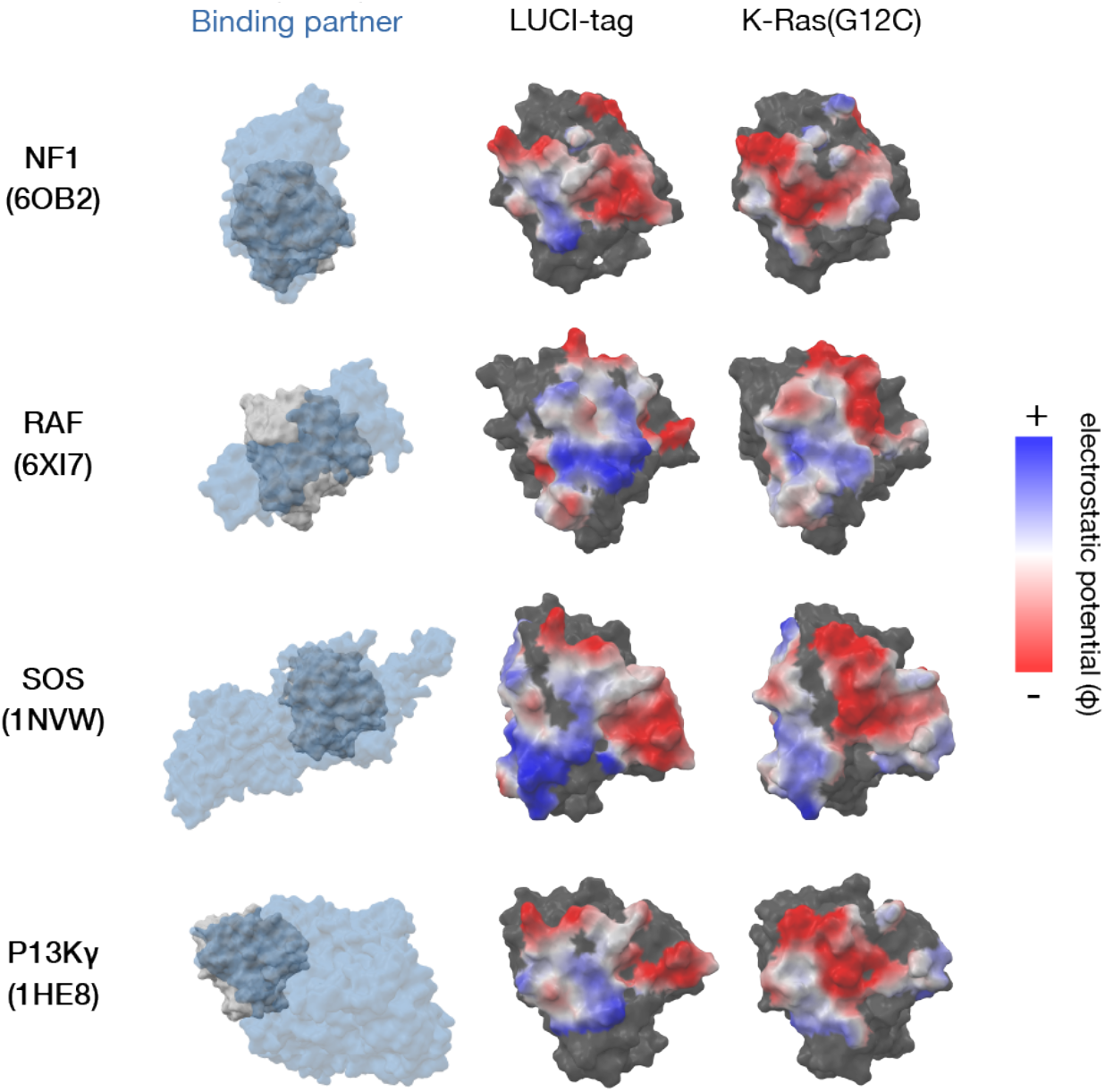
Electrostatic surface potential comparison at K-Ras(G12C) binding interfaces. Four representative structures of K-Ras(G12C) (gray) with binding partners (blue, transparent) are shown (NF1, PDB 6OB2; RAF, PDB 6XI7; SOS, PDB 1NVW; PI3Kγ, PDB 1HE8). The electrostatic surface potential of LUCI-tag and K-Ras(G12C) are shown on the right (red = negative, blue = positive). LUCI-tag has a markedly different electrostatic surface potential compared with K-Ras(G12C), which directly follows from our filtering for surface-charge dissimilarity during design.

**Figure S29.**
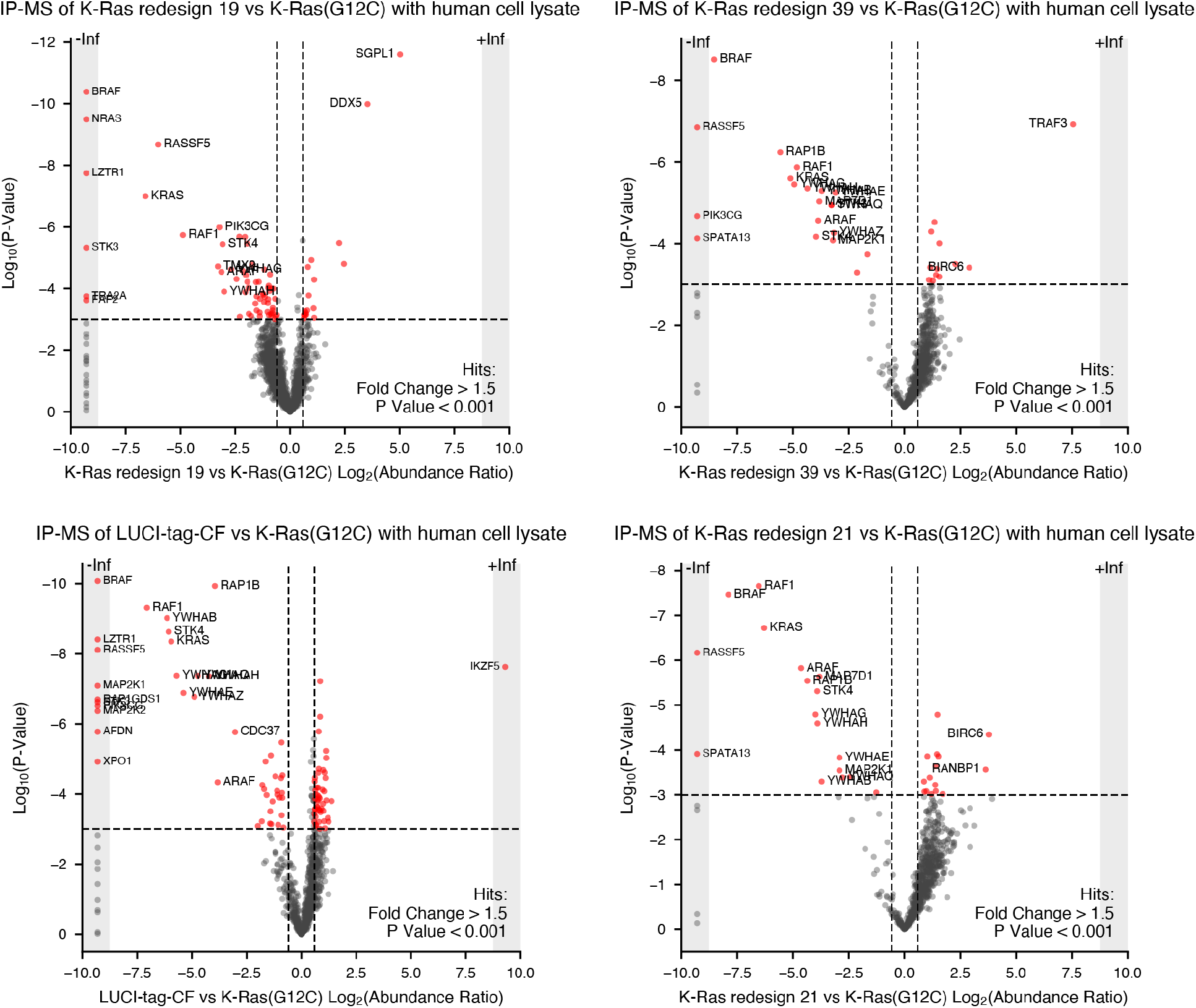
Immunoprecipitation-mass spectrometry (IP-MS) of LUCI-tag–CF and three other designs. FLAG–tagged constructs were immunoprecipitated from human cell lysates and analyzed by mass spectrometry. Each volcano plot shows the log_2_ abundance ratio of each construct versus K-Ras(G12C) against the log_10_ P-value. Red points are significant hits (fold change > 1.5, P < 0.001); gray points in the +Inf bands were fully imputed, indicating no peptides were detected in one comparison group across all four replicates. Known K-Ras interactors (e.g. BRAF, NRAS, RAF1, LZTR1, RASSF5) are depleted in the designs relative to K-Ras(G12C), consistent with loss of native K-Ras protein–protein interactions.

**Figure S30.**
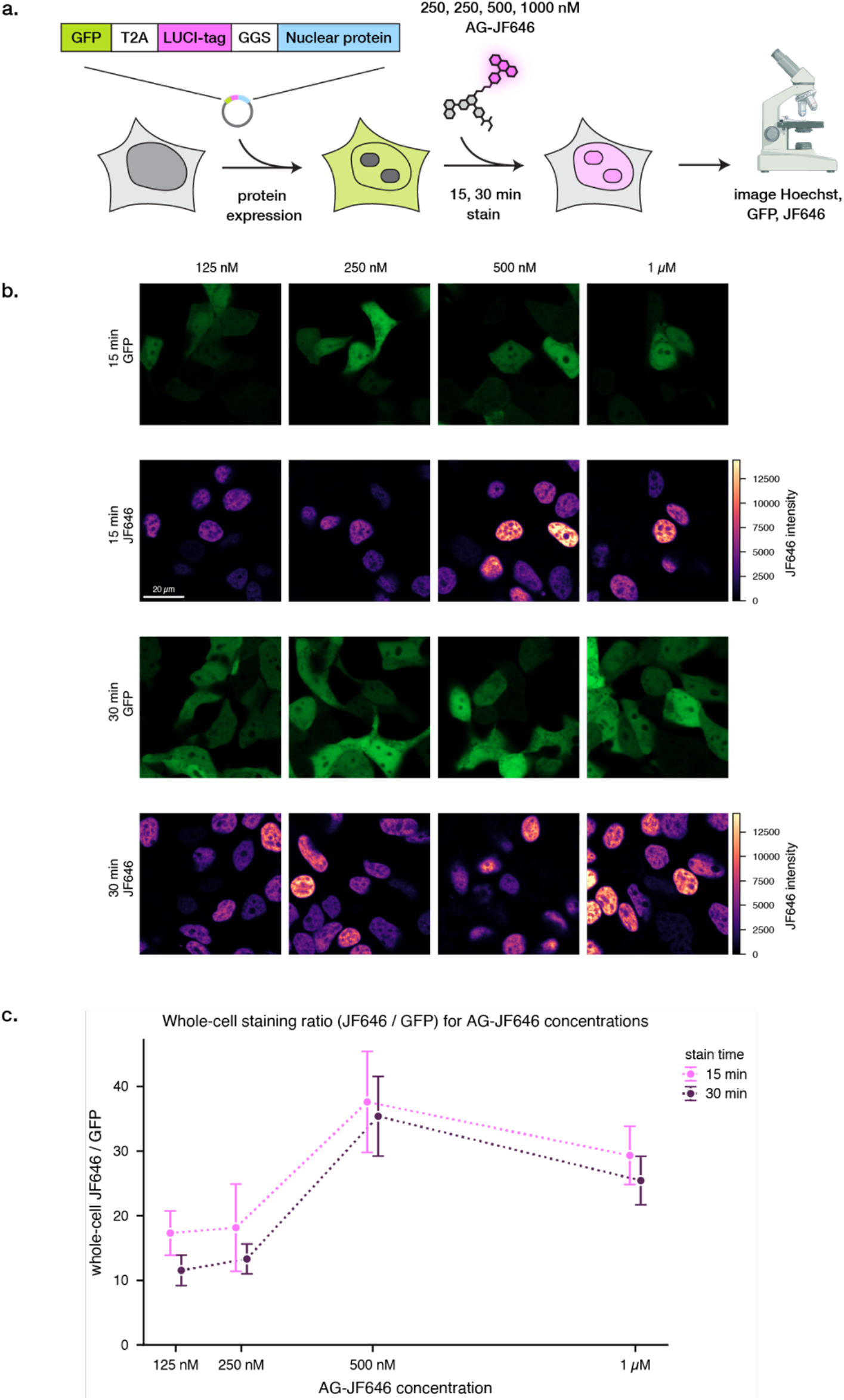
Quantification of LUCI-tag staining intensity at different AG-JF646 concentrations. **a,** HEK293T cells expressing untagged GFP and nuclear-localized LUCI-tag (GFP-T2A-LUCI-Tag–PITX2) were stained with 0.125–1 µM of AG–JF646 for 15 or 30 min and imaged without washing. **b,** Average whole-cell intensity in the GFP and JF646 channels were taken following image segmentation. **c,** The ratio of JF646 / GFP intensities was calculated to normalize for per-well differences in transfection and expression levels. Cells generally showed brighter JF646 staining at higher AG–JF646 concentrations and staining times, with an increase at 500 nM. Points show mean +/- S.E.M for N = 17 randomly selected GFP+ nuclei in each condition.

**Figure S31.**
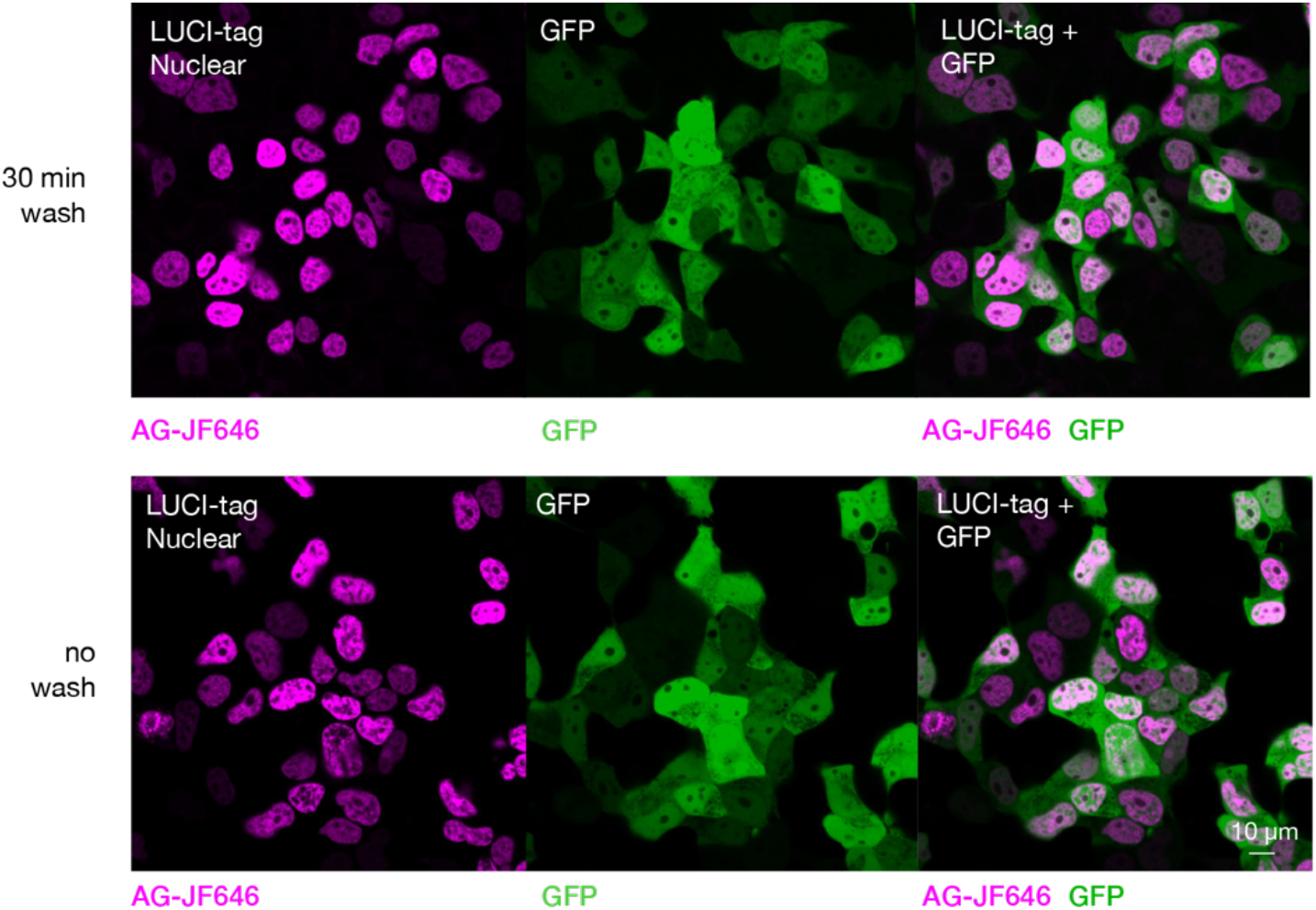
Comparison of imaging quality with and without wash steps with AG–JF646. HEK293T cells expressing nuclear-localized LUCI-tag (GFP-T2A-LUCI-Tag–PITX2) were stained with 1 µM AG– JF646 for 15 min and imaged directly or washed for 30 min in imaging media.

**Figure S32.**
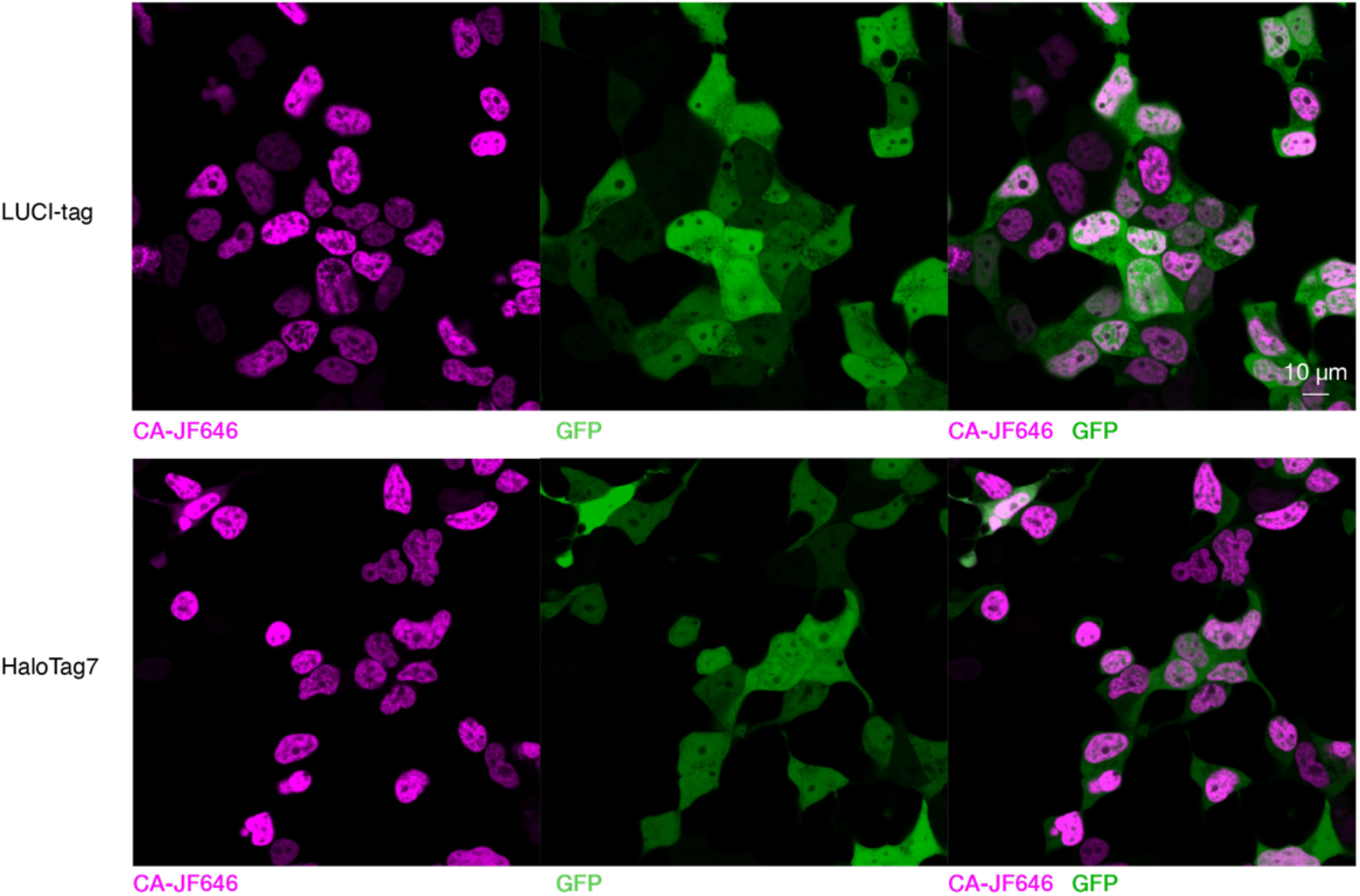
Comparison of nuclear-localized LUCI-tag and HaloTag7 staining in HEK293T cells under identical conditions. HEK293T cells were transfected with GFP-expressing, nuclear-localized LUCI-tag or HaloTag7 constructs and stained with 500 nM of their respective probes with JF646 payload (AG–JF646 for LUCI-tag, CA-JF646 for HaloTag7) for 1 hr. Both tags give nuclear-restricted staining in transfected, GFP-positive cells. Imaging conditions (laser power, exposure time) are kept constant but display range is scaled individually for each image.

**Figure S33.**
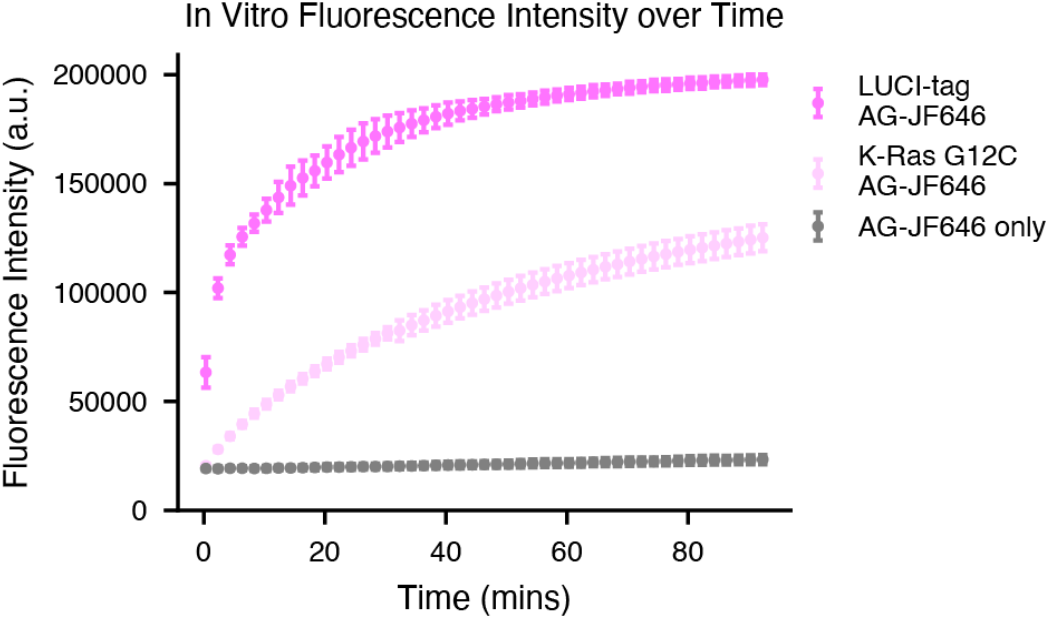
Time-resolved fluorescence intensity of purified LUCI-tag and AG–JF646. LUCI-tag showed modest fluorogenicity that was greater than K-Ras(G12C). Based on the measured k_app_ we expect covalent binding at these concentrations to be complete at sub-min timescales; however fluorescence turn-on appears to have a slower rate. Note that fluorogenicity of dye payloads was not explicitly considered in the design of LUCI-tag. Conditions: [AG–JF646] = 100 nM; [LUCI-tag] = 200 nM; [K-Ras(G12C)] = 200nM; buffer = protein buffer (see Methods). Error bars show standard deviation across 3 technical replicates.

**Figure S34.**
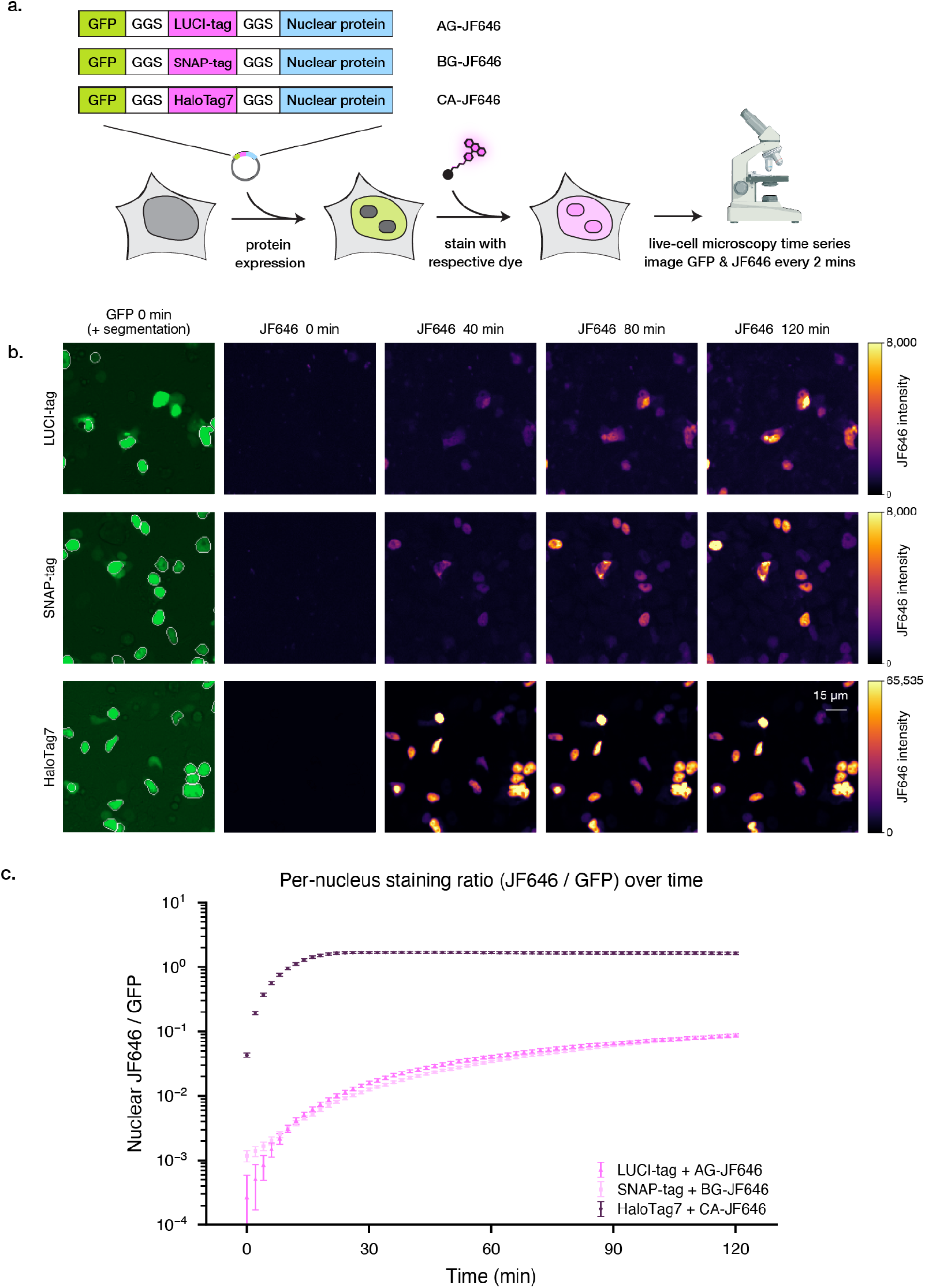
Comparison of LUCI-tag, HaloTag7, and SNAP-tag staining intensity using JF646 in live cells over time. **a,** HEK293T cells were transfected with GFP-SLP-PITX2 where SLP was LUCI-tag, SNAP-tag, or HaloTag7; note the lack of T2A means GFP and SLP were both nuclear-localized. Cells were stained with their respective probes bearing JF646 dye, followed by imaging every 2 min. **b,** Nuclei were separated by image segmentation via the GFP channel and the intensity ratio of JF646 / GFP is reported. Points show mean nuclear intensity per timepoint +/- SEM. **c,** While HaloTag7 shows faster staining kinetics, LUCI-tag with AG–JF646 showed similar kinetics to SNAP-tag with BG-JF646. Because this system is fluorophore matched, differences in speed are due to differences in probe permeability, reaction speed, and fluorogenicity. JF646 intensity color bars are shared for LUCI-tag and SNAP-tag.

**Figure S35.**
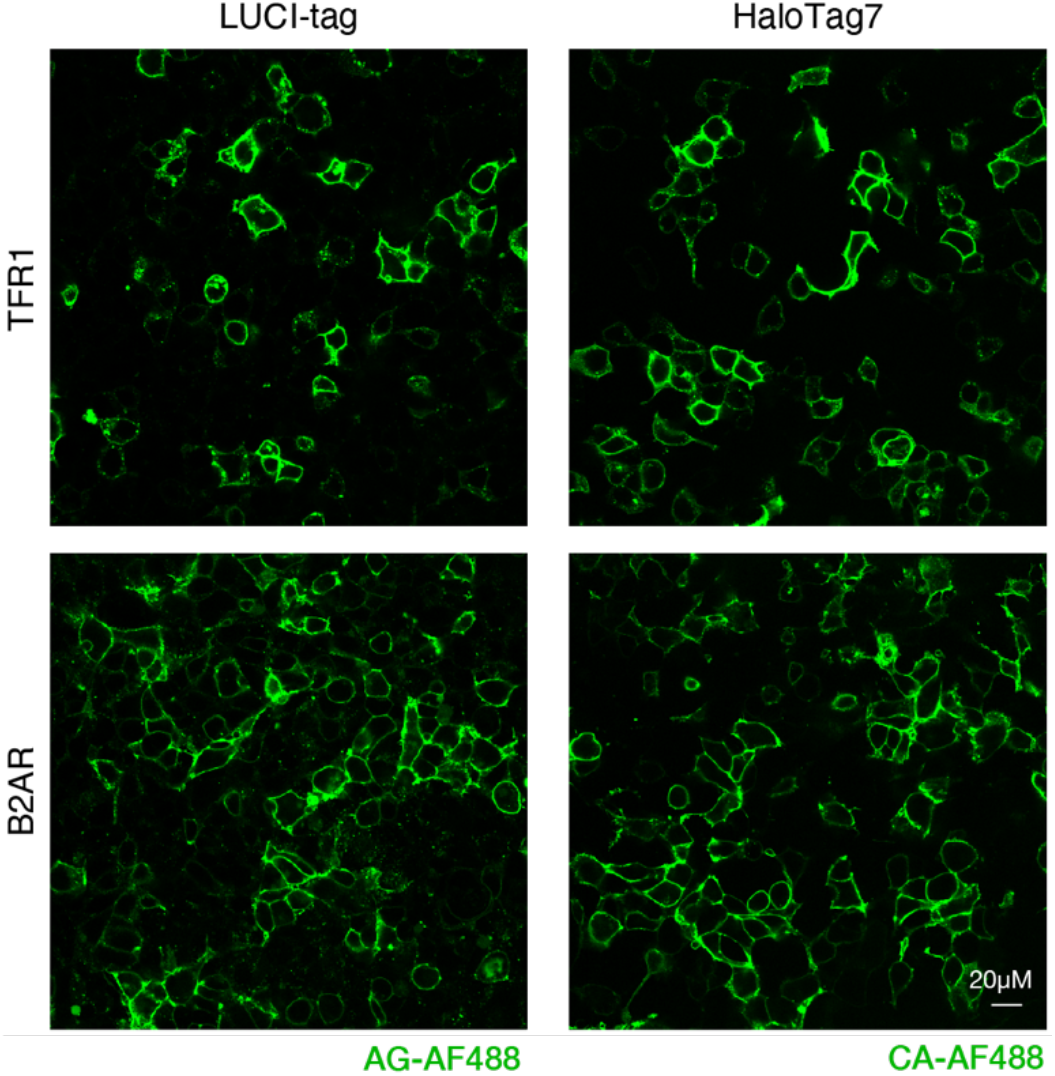
Comparison of staining patterns for membrane-localized LUCI-tag and HaloTag7. HEK293T cells were transfected with SLP constructs fused to β2 adrenergic receptor (B2AR) or transferrin receptor (TFR1) and stained with 500 nM of their respective AF488-bearing probes for 30 min, followed by 3x PBS washes. HaloTag7 and LUCI-tag show similar staining completion and patterns.

**Figure S36.**
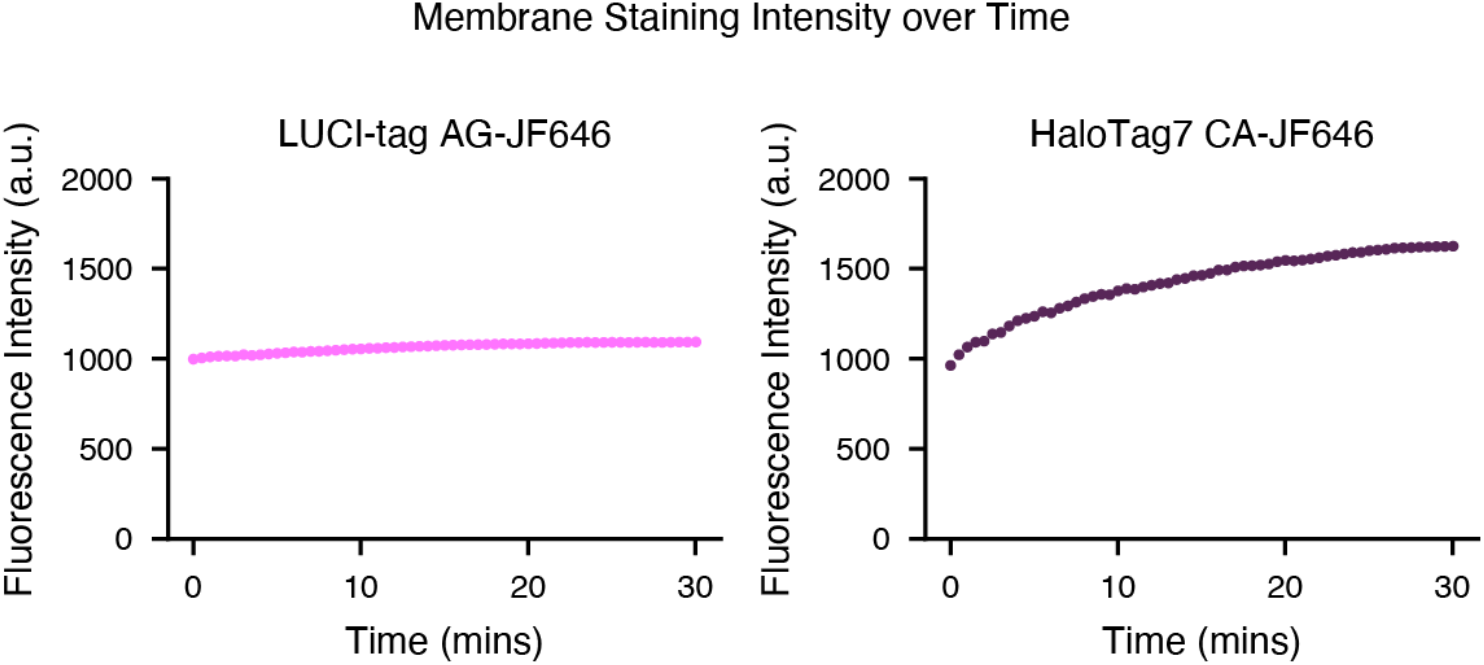
Time-resolved membrane staining in live-cell microscopy. HEK293T cells expressing LUCI-tag–B2AR or HaloTag7–B2AR (see Fig. S35) were stained with their respective probes bearing JF646 dye and imaged every 2 min on the same plate with similar settings. Both LUCI-tag and HaloTag7 show immediate labeling at the earliest possible timepoints. HaloTag7 shows a slow rise in intensity due to focal shift over time. B2AR = β2 adrenergic receptor.

**Figure S37.**
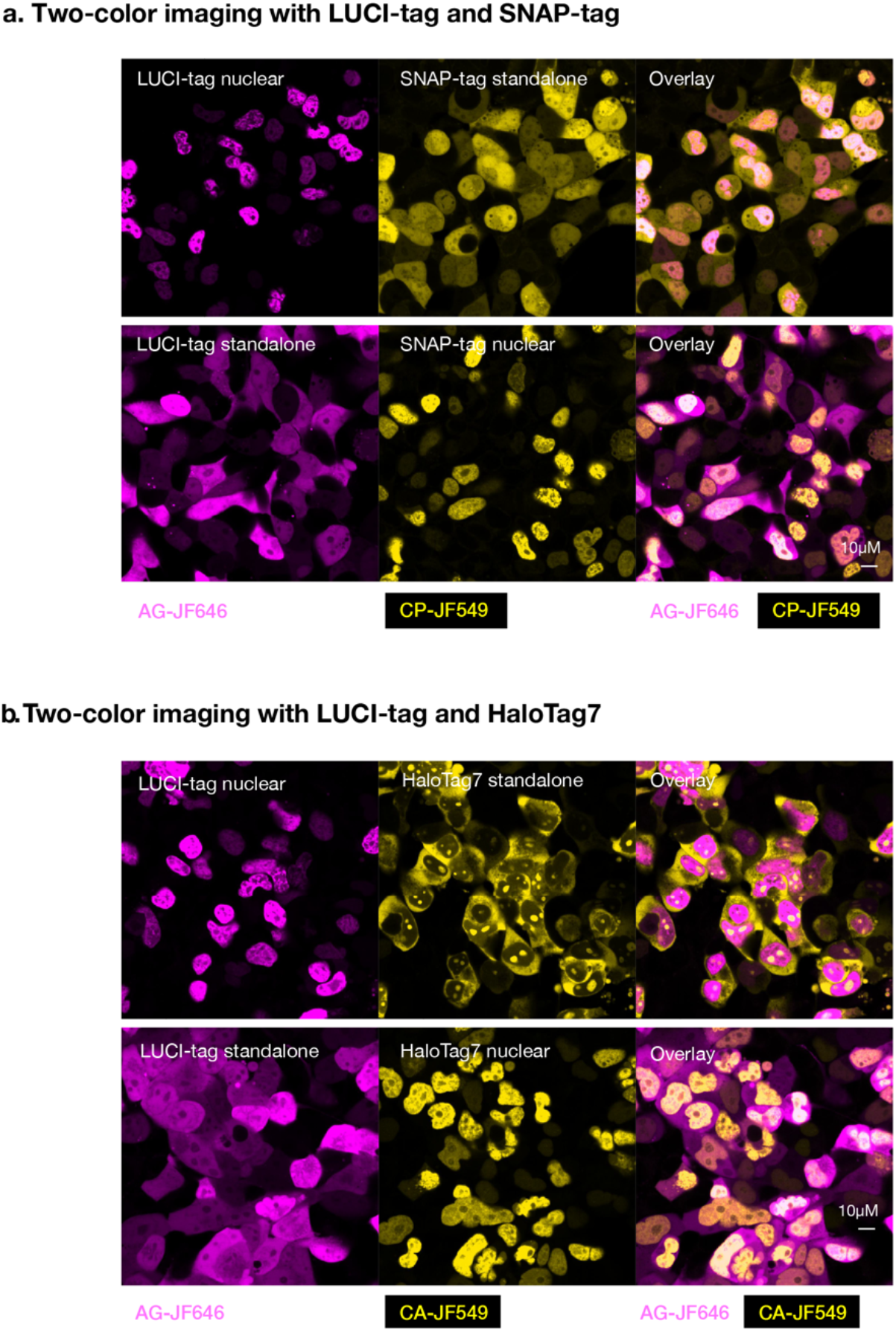
Two-color imaging with LUCI-tag and SNAP-tag or LUCI-tag and HaloTag7. HEK293T cells were co-transfected with two plasmids followed by a 30 min stain at 500 nM dye and imaging. LUCI-tag was paired with SNAP-tag (**a**, top) or HaloTag7 (**b**, bottom) for two-color imaging, where each tag was either nuclear-localized or untagged (diffuse localization between nucleus and cytoplasm). In each instance, the nuclear-localized tags do not have any cytoplasmic background staining, showing that there is little off-target reactivity between protein tags and their probes.

**Figure S38.**
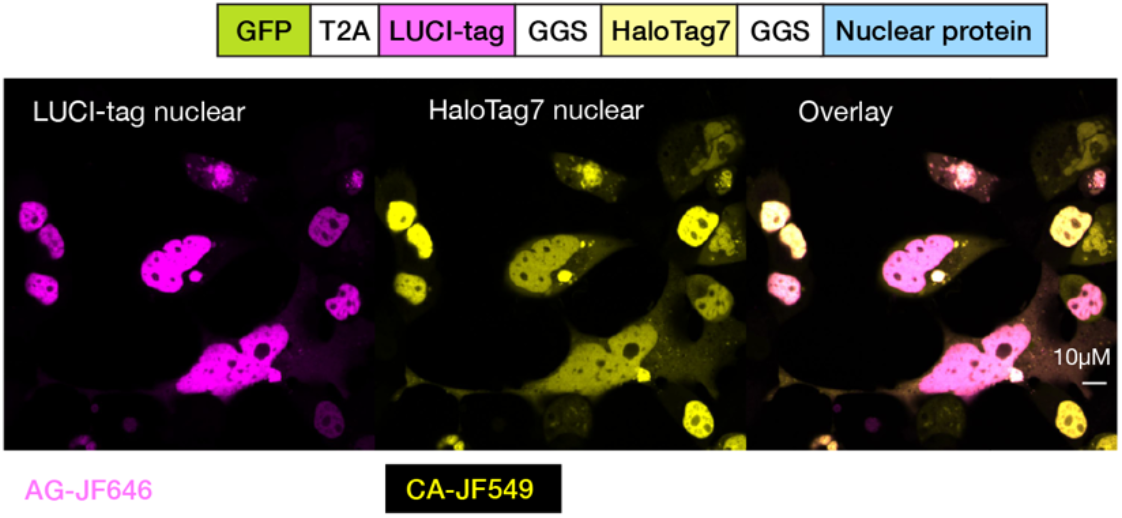
Two-color imaging with tandem fusion of HaloTag7 and LUCI-tag. HEK293T cells were transfected with a single construct containing LUCI-tag fused to HaloTag7 and a nuclear-localized protein. Staining at 500 nM for 30 min showed that LUCI-tag and HaloTag7 had identical staining patterns.

**Figure S39.**
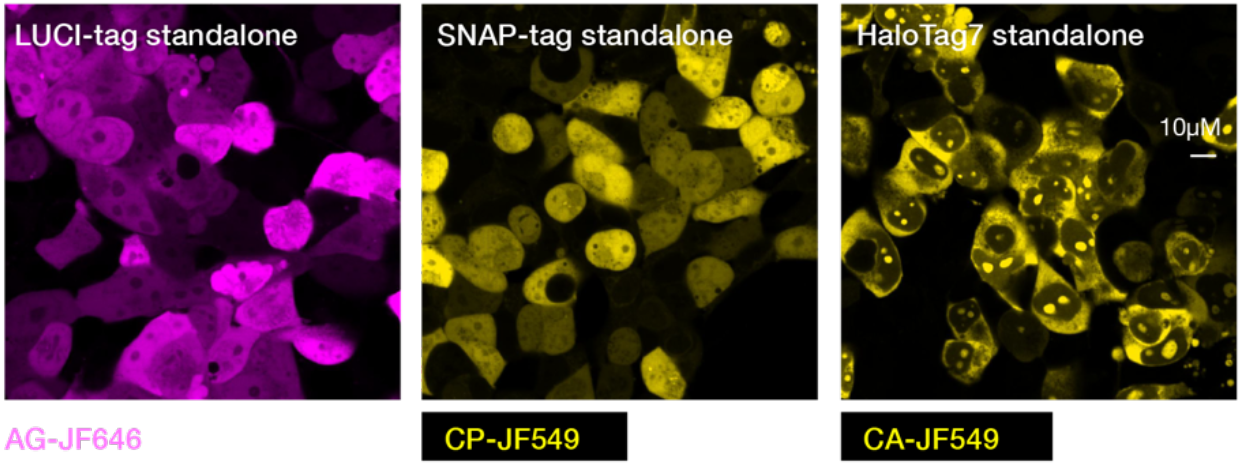
Staining patterns of standalone LUCI-tag, SNAP-tag, and HaloTag7. HEK293T cells transfected with standalone SLP constructs were stained for 500 nM and 30 min with their respective probe–dye conjugates. Unlike LUCI-tag and SNAP-tag, HaloTag7 staining appeared limited to the cytoplasm, with nuclear puncta, perhaps due to HaloTag7’s large size and inability to traverse the nuclear pore complex without attachment to a nuclear-localization signal.

**Figure S40.**
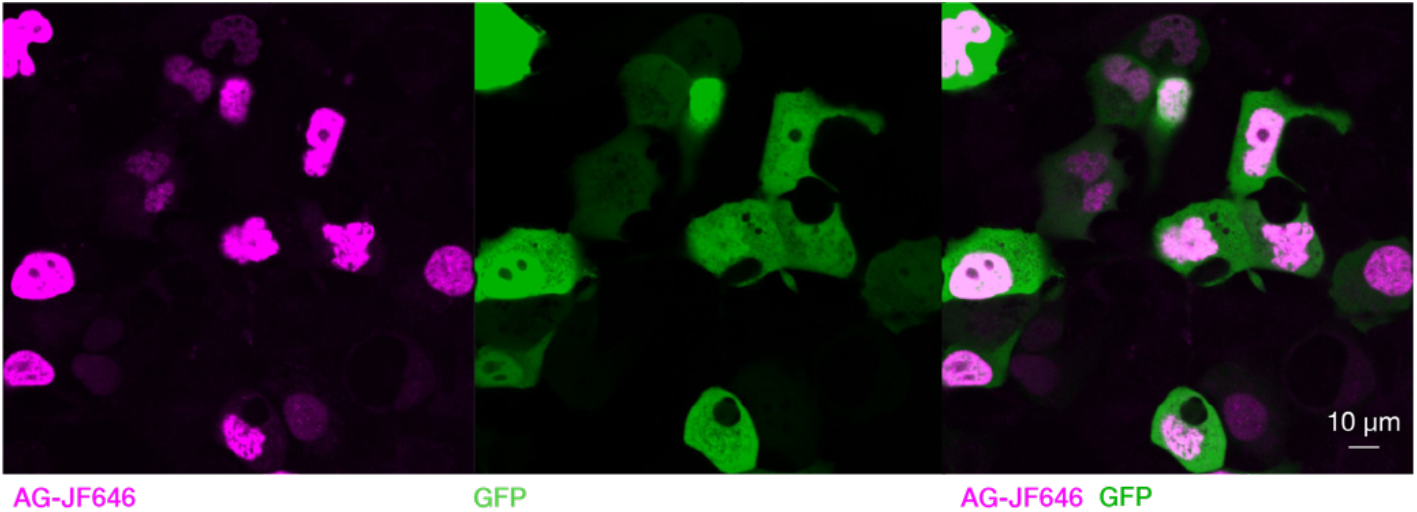
Imaging of nuclear LUCI-tag–CF in HEK293T cells. HEK293T cells expressing standalone GFP and nuclear-localized LUCI-tag–CF (GFP-T2A-PITX2-LUCI-tag–CF) were stained with 500 nM AG–JF646 for 30 min and imaged. LUCI-tag–CF shows the expected staining and localization, indicating that mutations that hinder binding of the nucleotide cofactor did not impair the ability to label the protein intracellularly.

**Figure S41.**
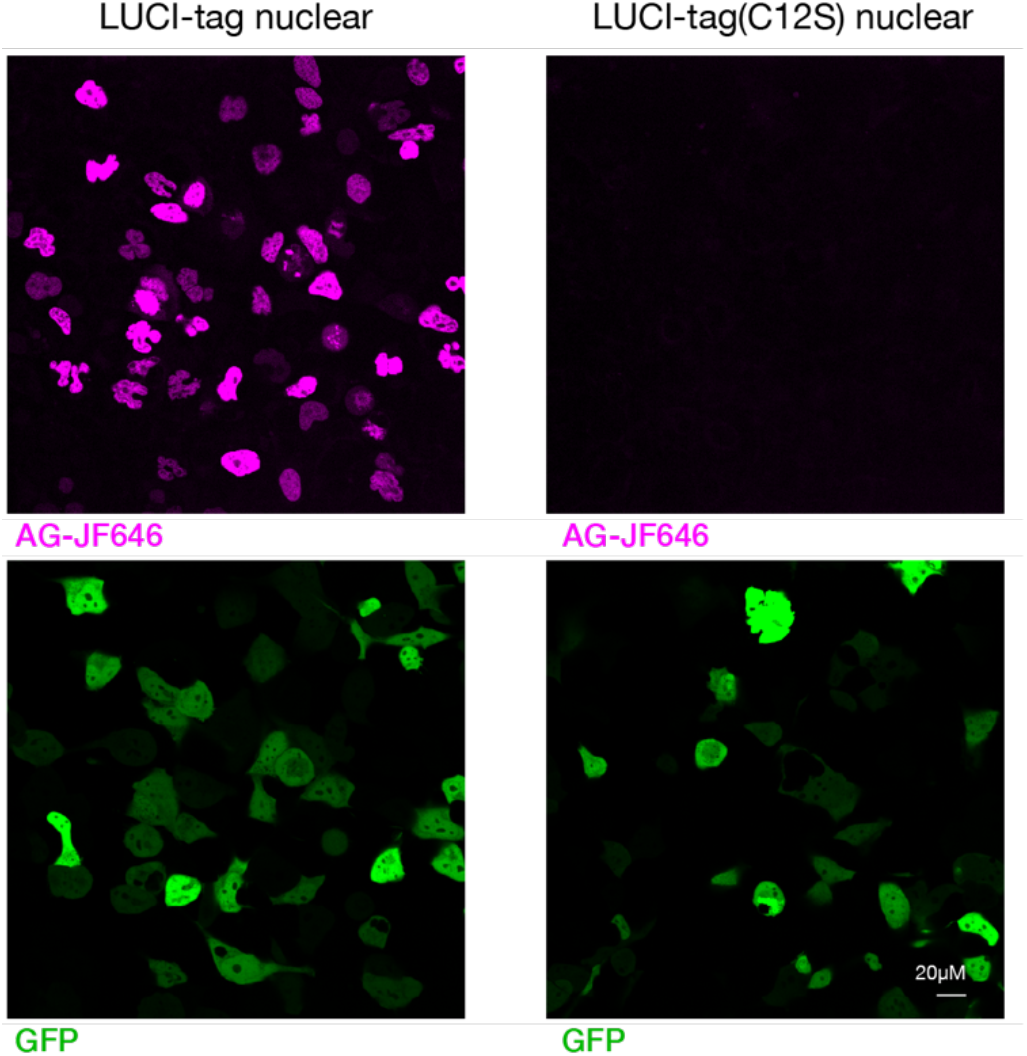
Imaging of LUCI-tag(C12S) in HEK293T cells shows no staining. HEK293T cells expressing nuclear-localized LUCI-tag(C12S) (GFP-T2A-LUCI-tag(C12S)-PITX2) were stained with 500 nM AG–JF646 for 30 min with the same microscope settings and display range as the LUCI-tag control (GFP-T2A-LUCI-tag–PITX2). The Cys to Ser mutation is incapable of covalent probe binding, leading to a lack of nuclear-specific staining in cells.

**Figure S42.**
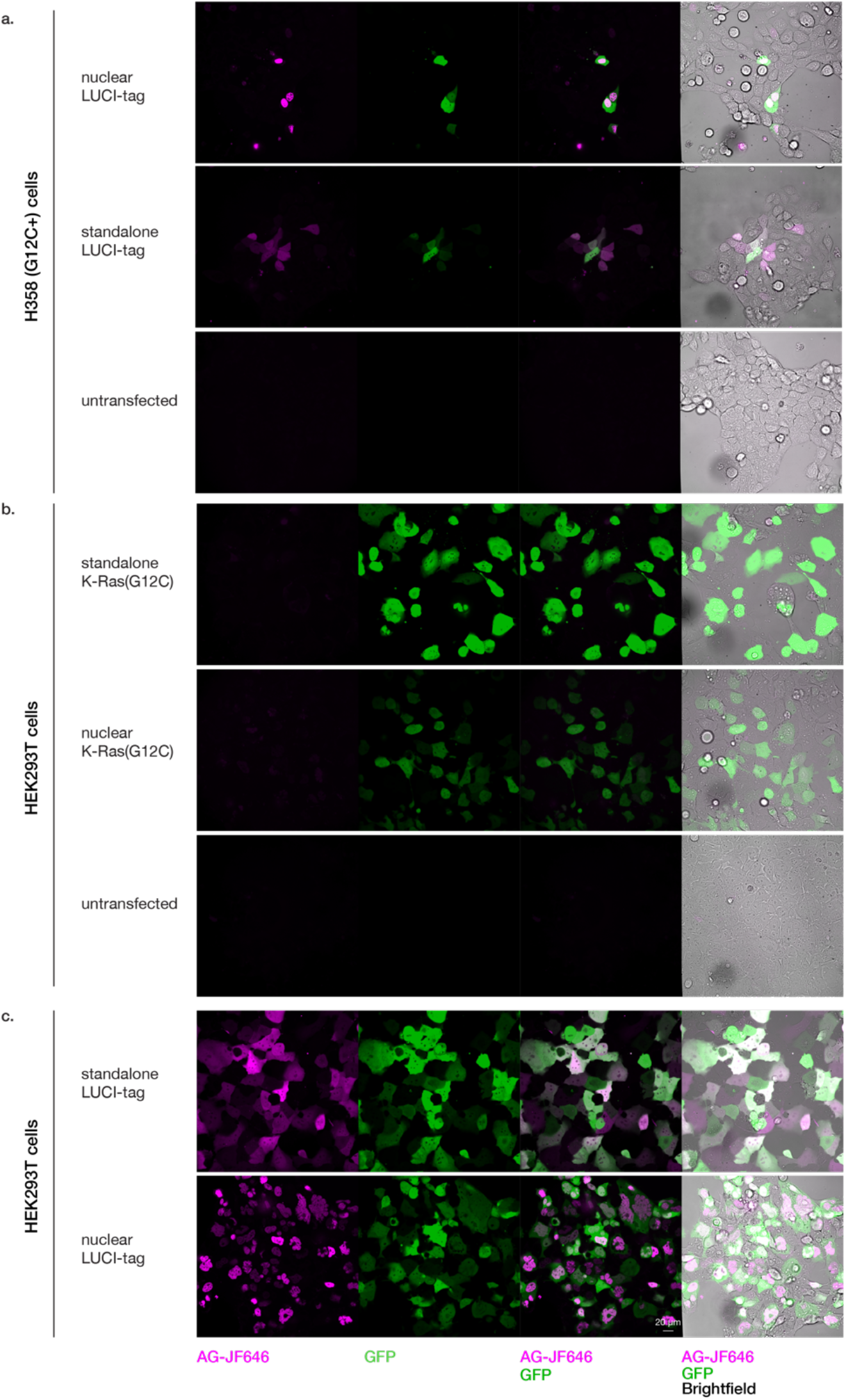
Control imaging in H358 (G12C+) cells and HEK293T cells transfected with K-Ras(G12C). **a,** H358 cells, which endogenously express K-Ras(G12C), were transfected with cytoplasmic LUCI-tag, nuclear-localized LUCI-tag, or left untransfected; although transfection/expression is poor, LUCI-tag staining is still clearly visible, and no background staining is seen in the untransfected cells. **b,** HEK cells were transfected with K-Ras(G12C)ΔHVR constructs, both untagged and nuclear-localized; although expression is robust, there is no visible staining, further confirming that K-Ras(G12C) does not function adequately as an SLP for imaging without sequence redesign. **c,** normal LUCI-tag transfection and staining performed on the same plate for brightness comparison. All staining was performed with 500 nM AG–JF646 for 1 hr with identical imaging and image display range conditions.

**Figure S43.**
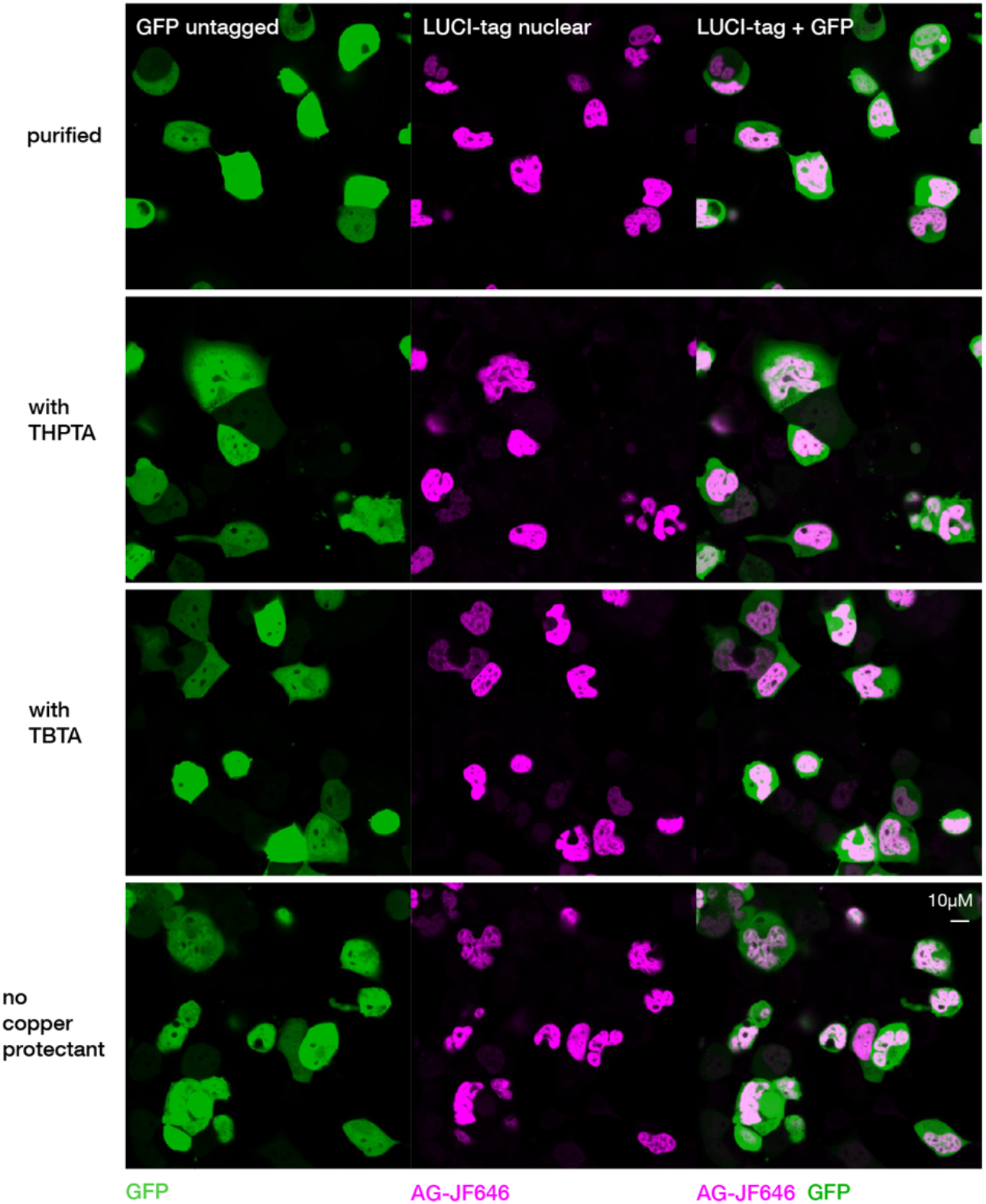
"Direct to Biology" AG–JF646 performs similarly to purified AG–JF646 in microscopy. AG–JF646 was made using the “direct to biology” probe generation protocol (see Methods), and its performance in cell microscopy was compared to AG–JF646 purified using a conventional HPLC purification protocol. Because copper toxicity is a concern for cell viability, 3 different copper protectant conditions were tested: THPTA, TBTA, and no copper protectant. HEK293T cells were transfected with nuclear-localized LUCI-tag (GFP-T2A-LUCI-tag–PITX2) and stained with 500 nM probe–dye conjugate for 30 min; there was no observable difference in imaging quality and cell morphology for all samples. THPTA=Tris(3-hydroxypropyltriazolylmethyl)amine, TBTA= Tris(benzyltriazolylmethyl)amine.

**Figure S44.**
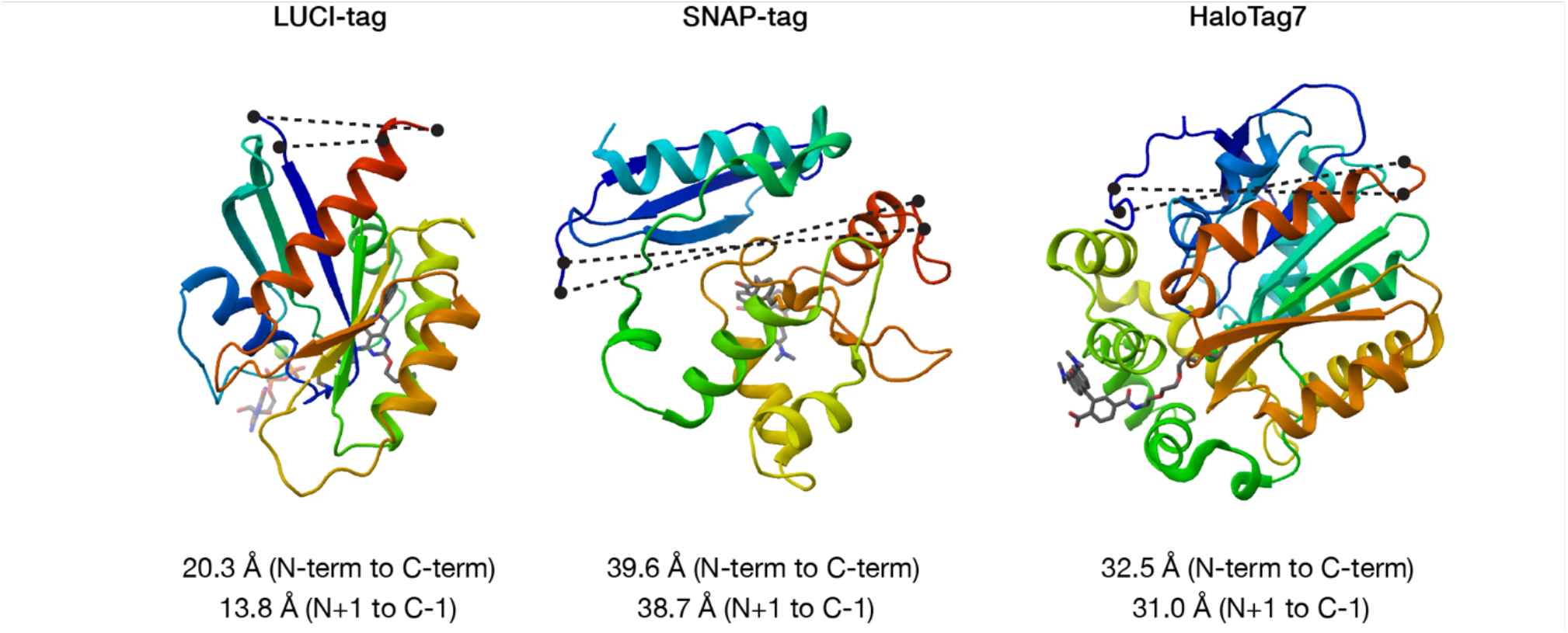
Cα distance between N- and C-termini for LUCI-tag, SNAP-tag (6Y8P), and HaloTag7 (6Y7A).

**Figure S45.**
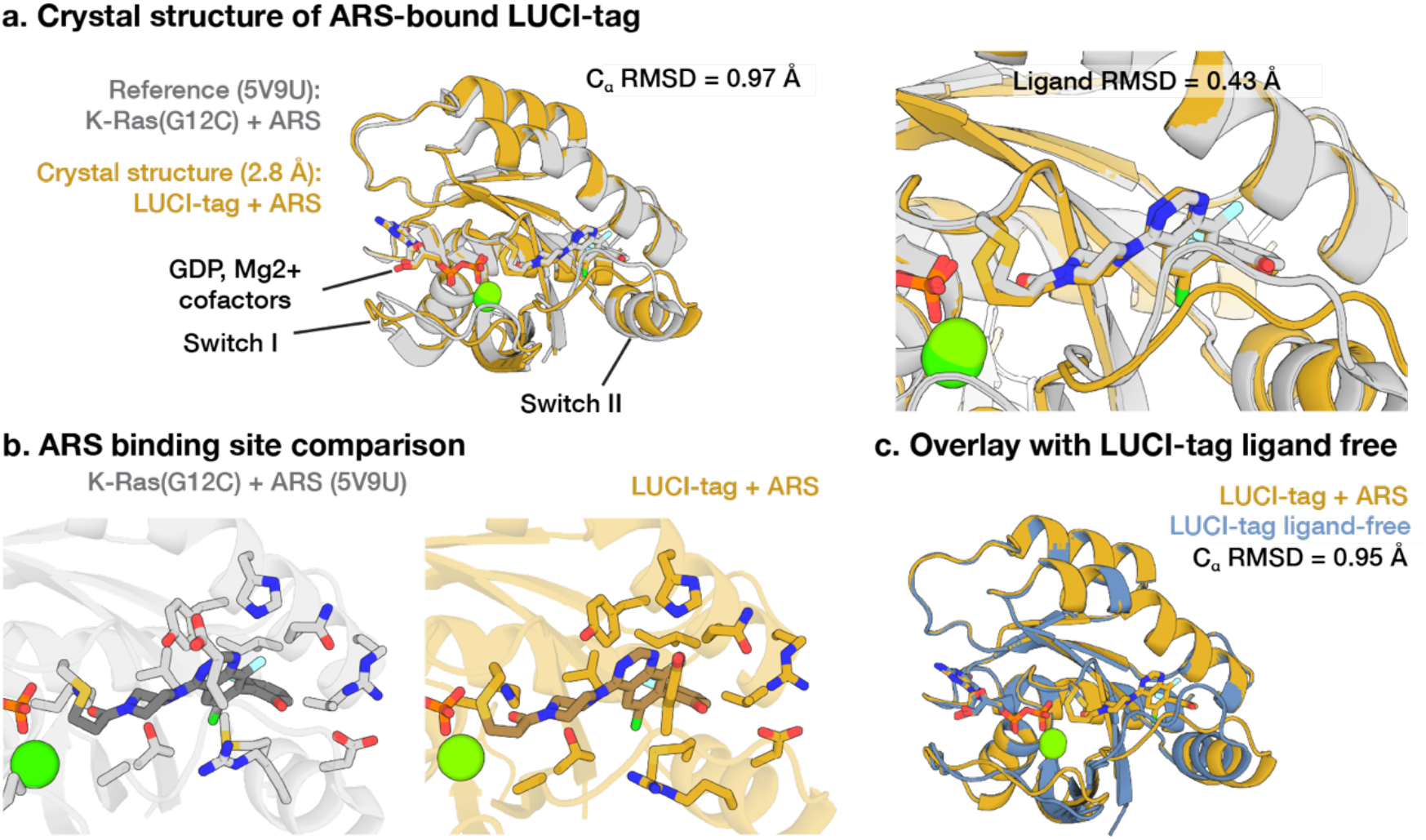
Crystal structure of LUCI-tag bound to ARS-1323. **a,** ARS-bound LUCI-tag (yellow) overlaid with a structure of ARS-bound K-Ras(G12C) showing low backbone and ligand RMSD. **b,** Binding site comparison showing conserved sidechain rotamers. **c,** Ligand free LUCI-tag (blue) overlaid with ARS-bound LUCI-tag (yellow).

**Figure S46.**
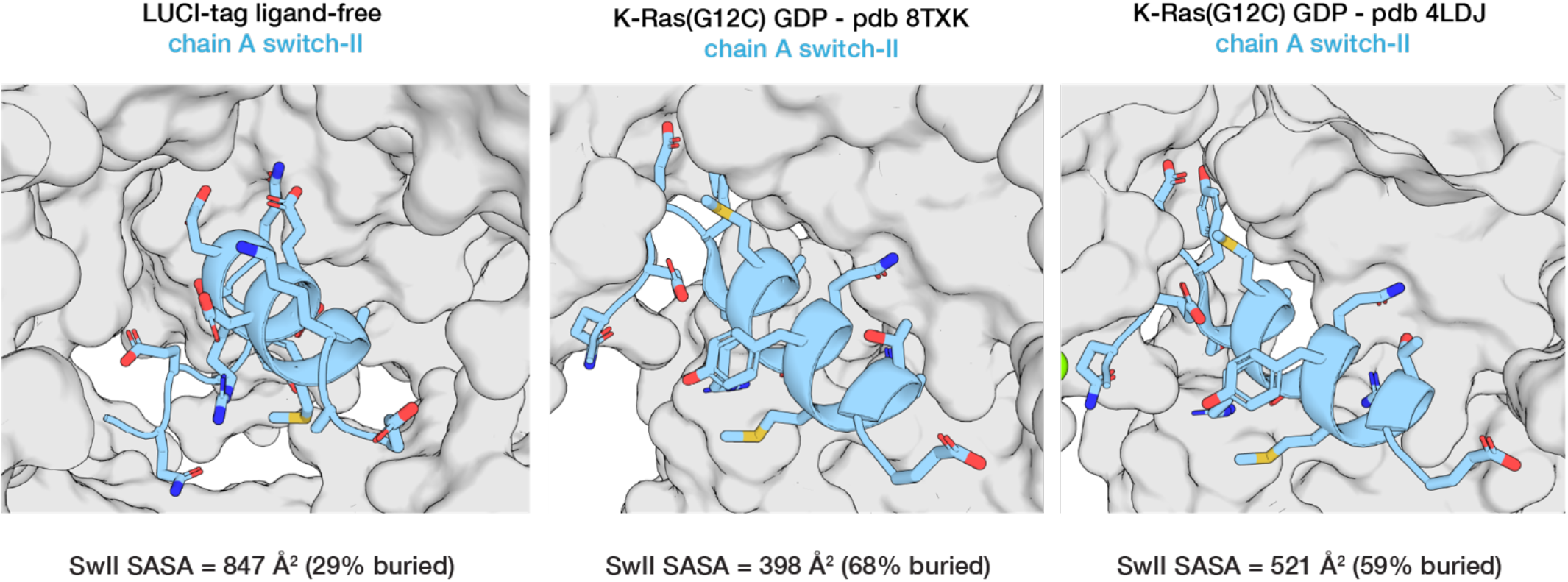
Crystallographic contacts of switch-II region of ligand-free LUCI-tag. Ligand-free (GDP-bound) LUCI-tag (left) has significantly higher solvent-accessible surface area (SASA) and is consequently less packed within neighboring chains in the crystal lattice, compared to crystal structures 8TXK and 4LDJ of K-Ras(G12C) (right).

**Table S1.**
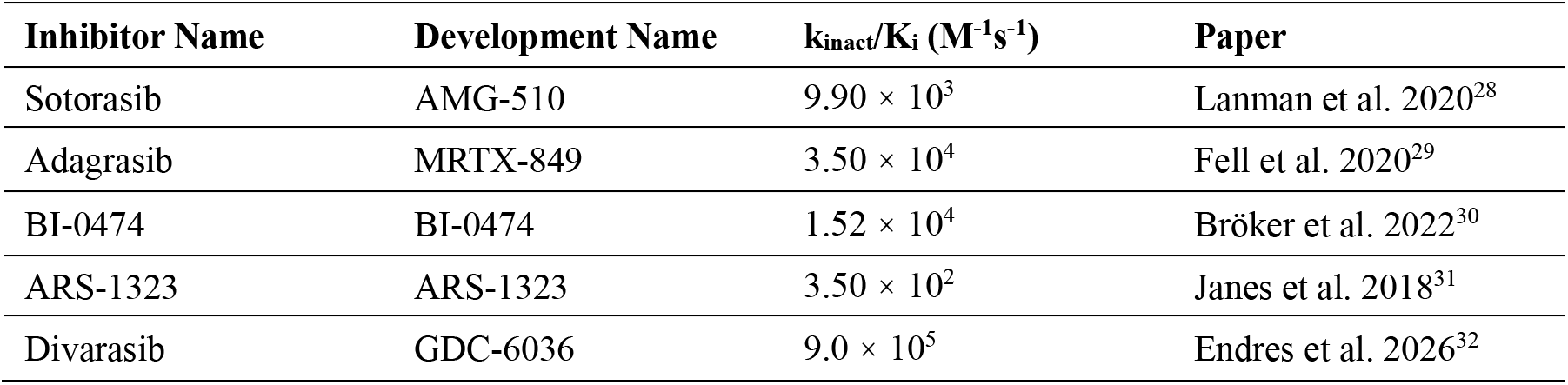
Second-order rate constants of K-Ras(G12C) considered in this study.

**Table S2.**
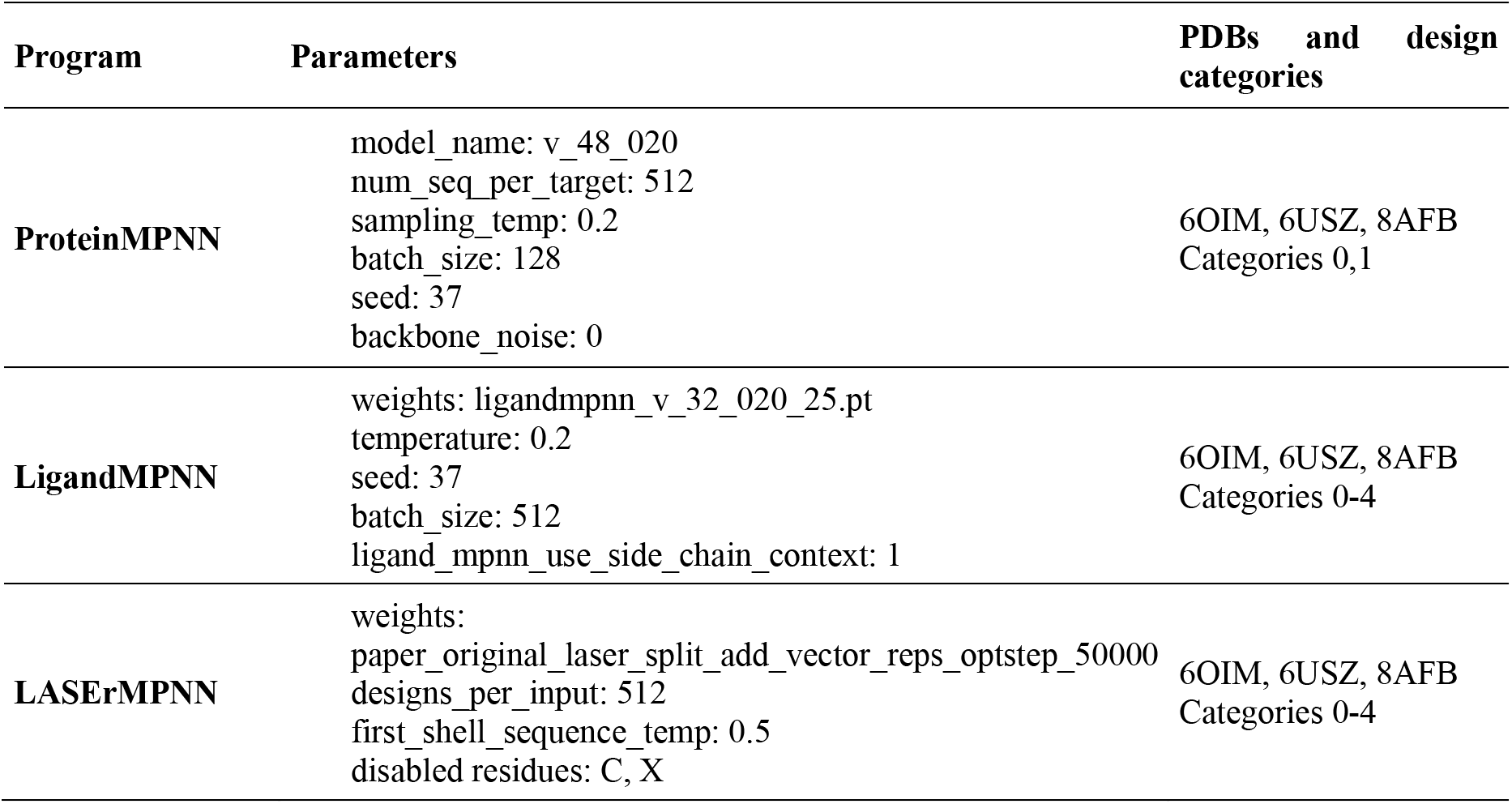
Design parameters for ProteinMPNN, LigandMPNN, and LASErMPNN.

**Table S3.**
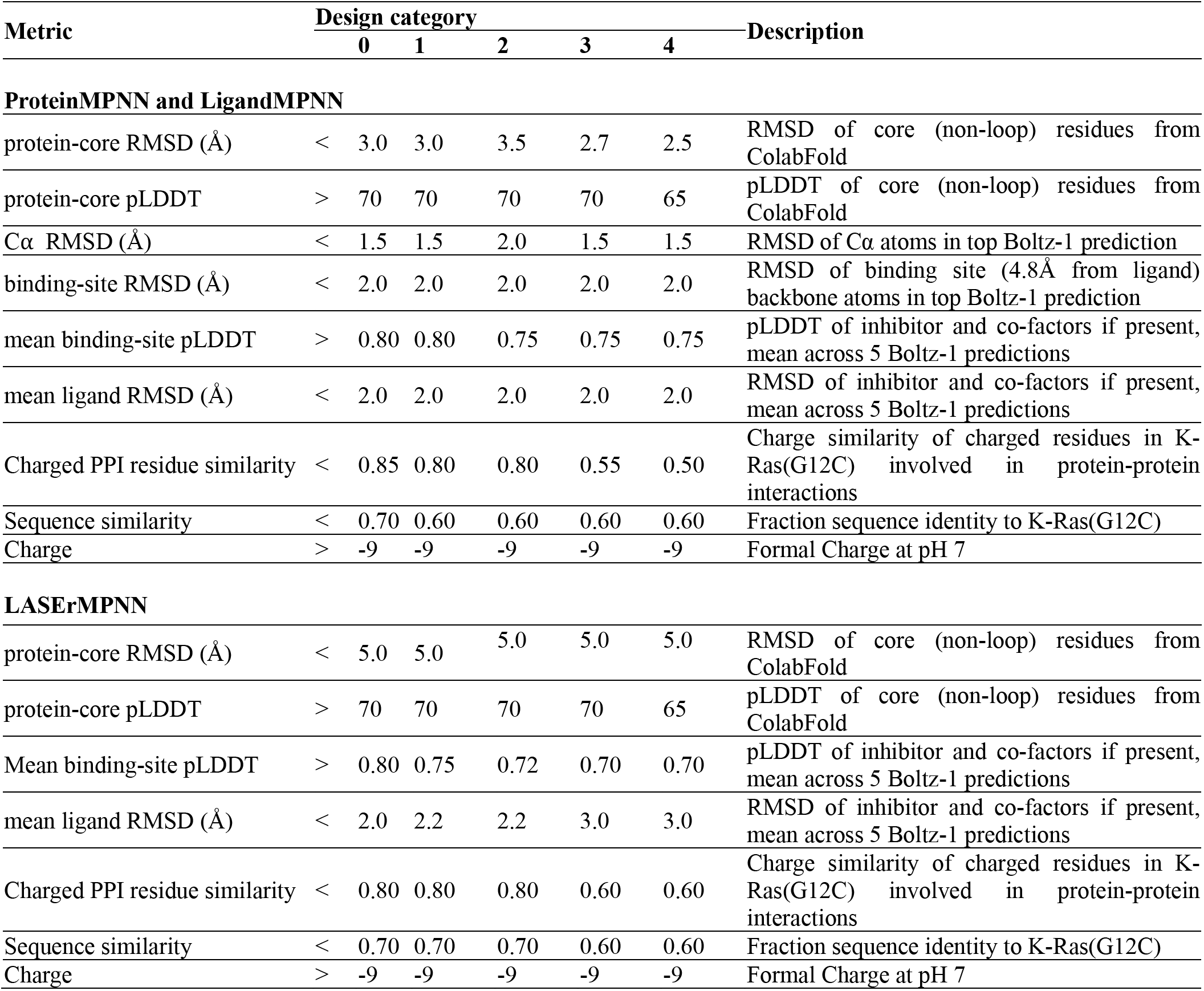
Computational filtering thresholds applied to each sequence design category.

**Table S4.**
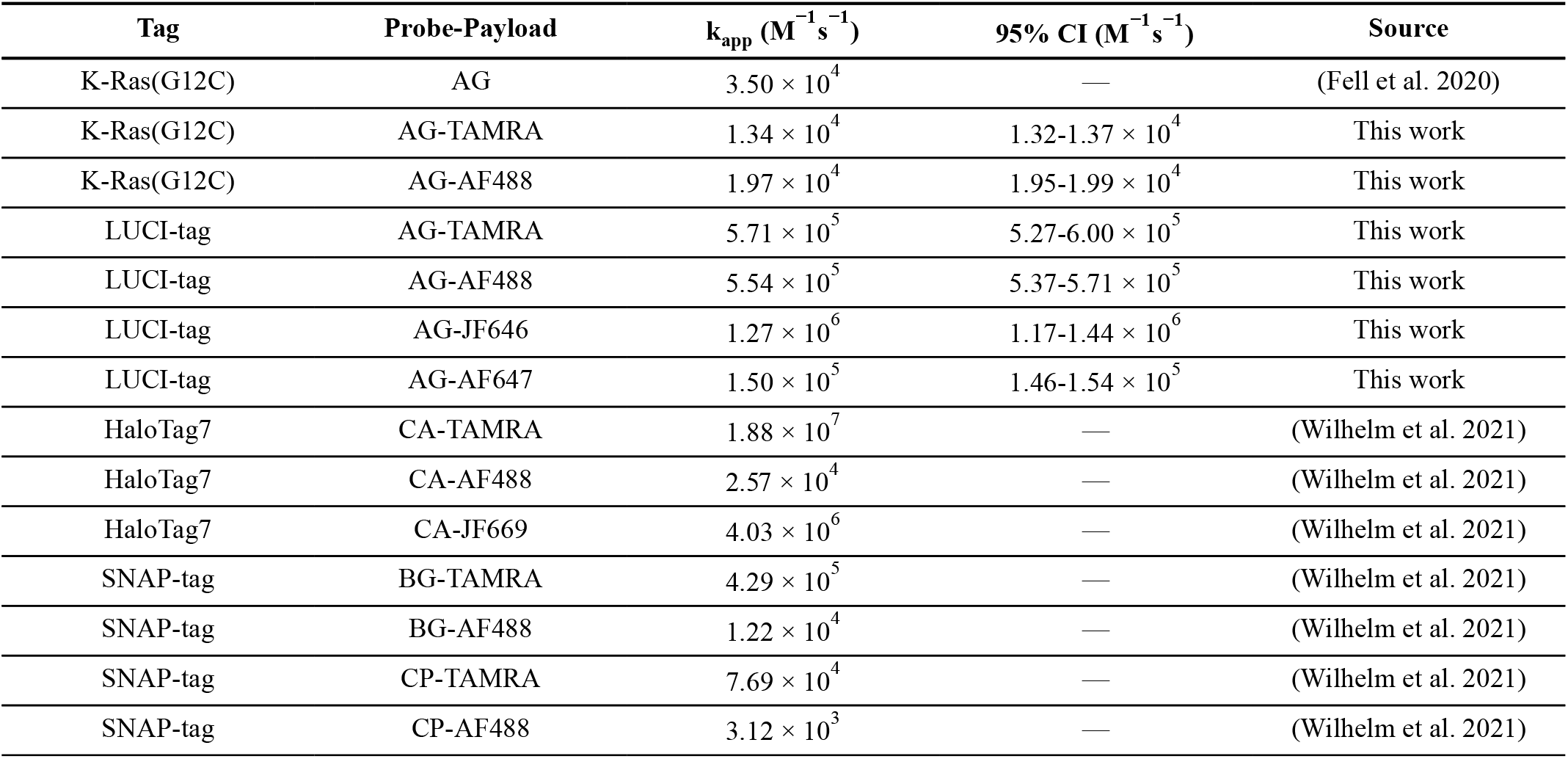
k_app_ of LUCI-tag, K-Ras(G12C), HaloTag7, and SNAP-tag with various payloads.

**Table S5.**
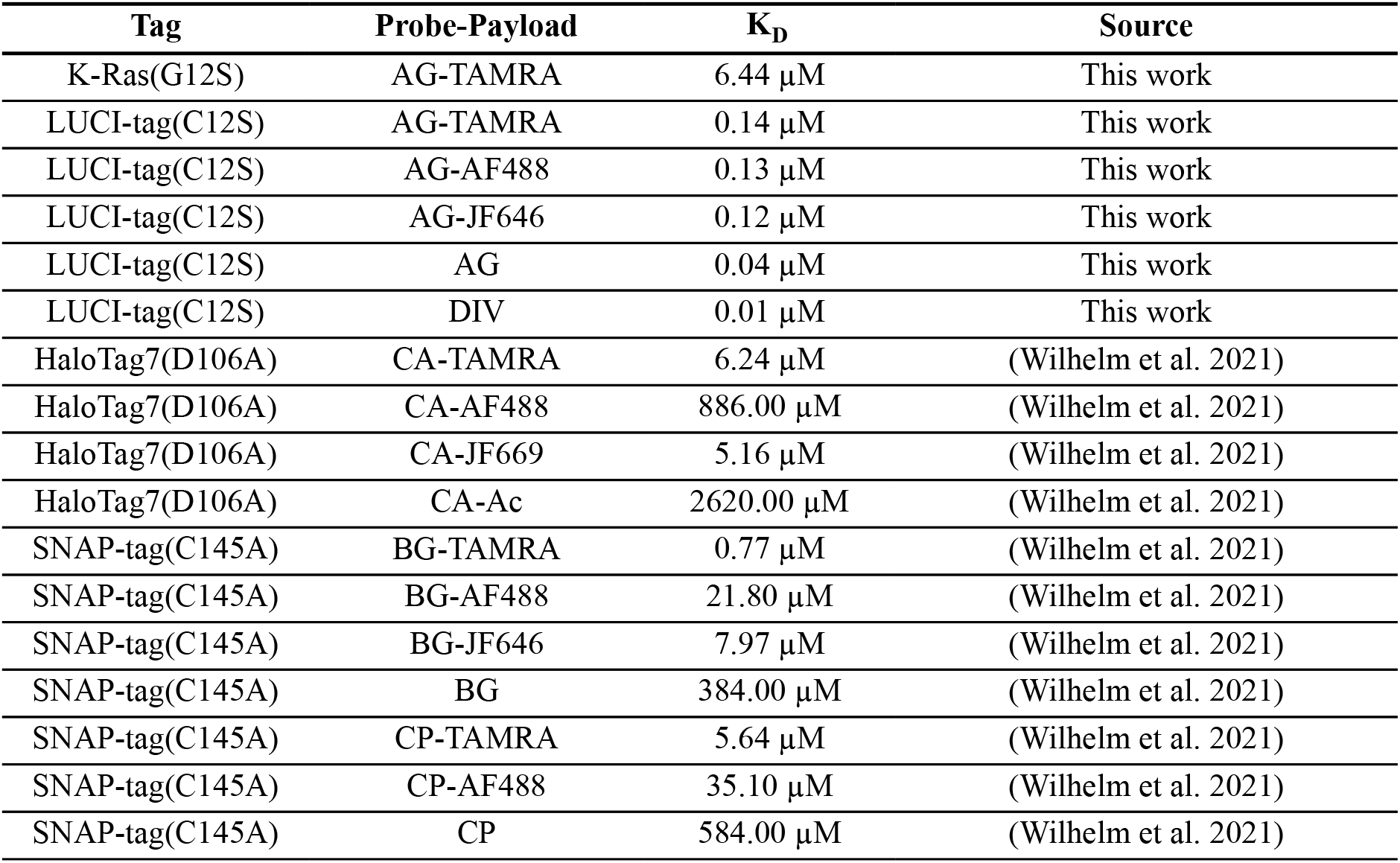
Noncovalent binding affinities of LUCI-tag(C12S), K-Ras(G12S), HaloTag7(D106A), and SNAP-tag(C145A).

**Table S6.**
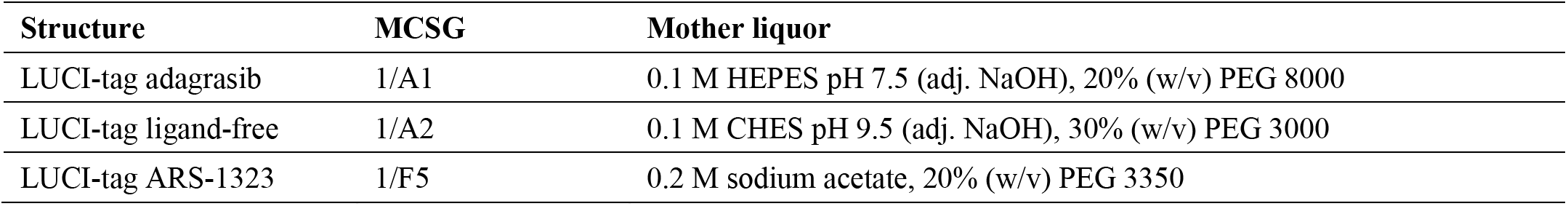
Crystallization conditions.

**Table S7.**
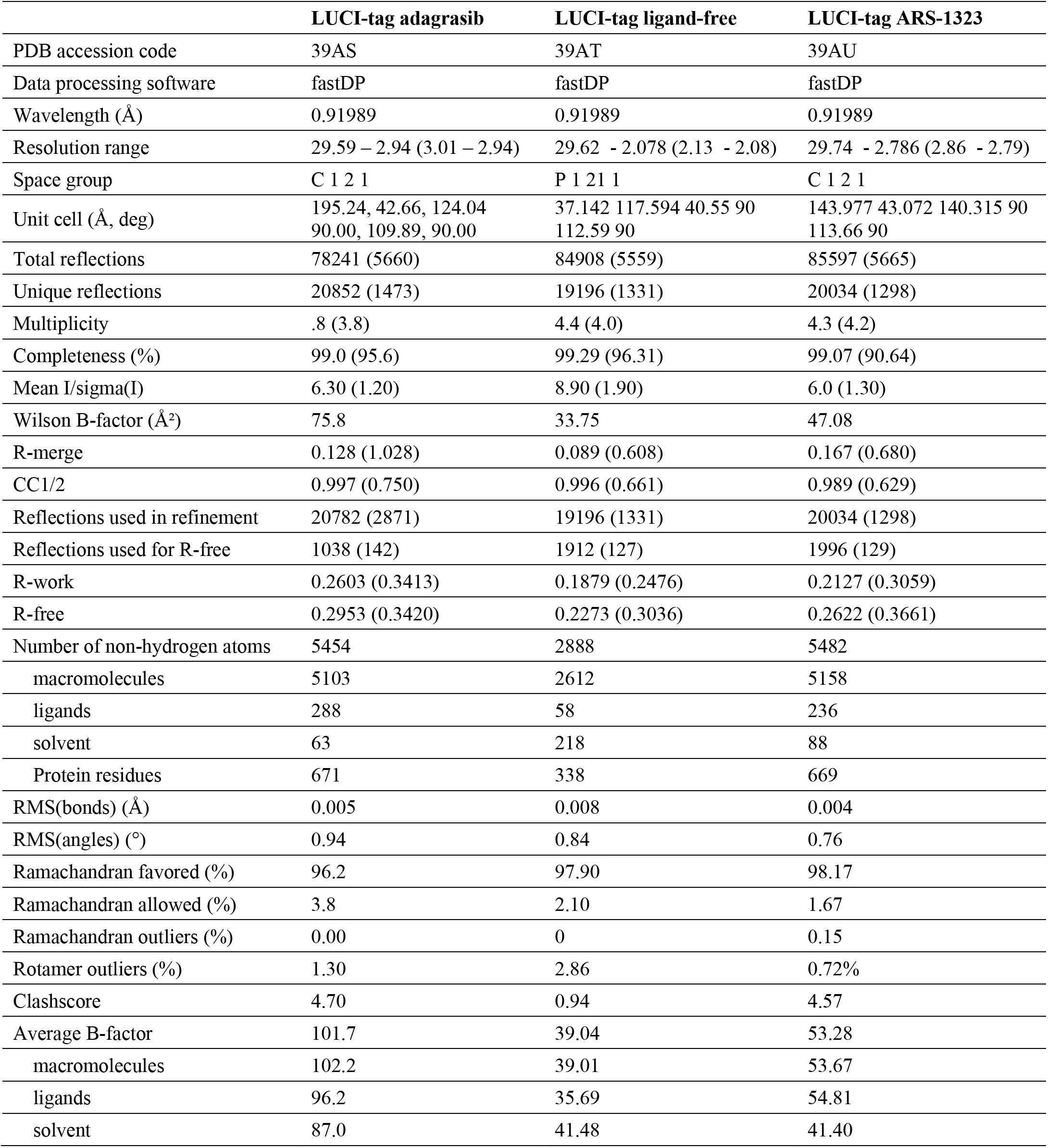
Crystallography statistics for all LUCI-tag structures.

**Table S8.**
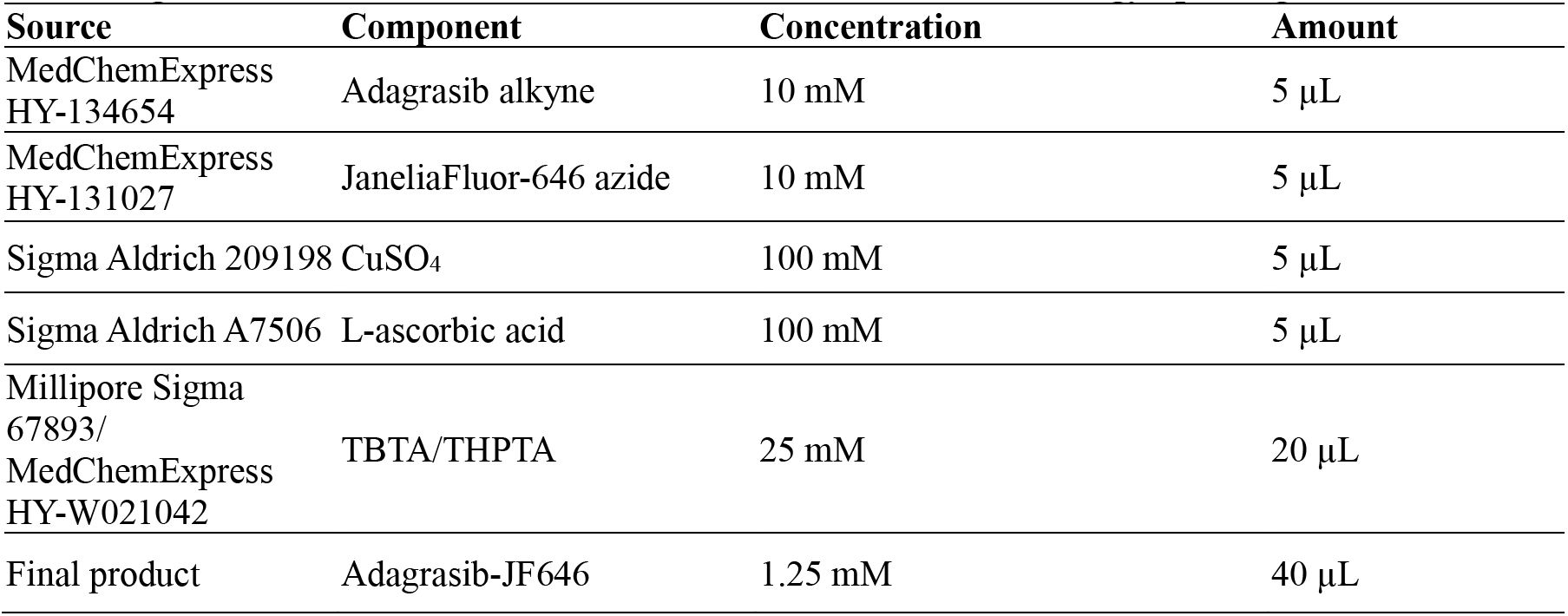
Reagent sources, concentrations, and amounts for "Direct to Biology" probe generation.

**Table S9.**
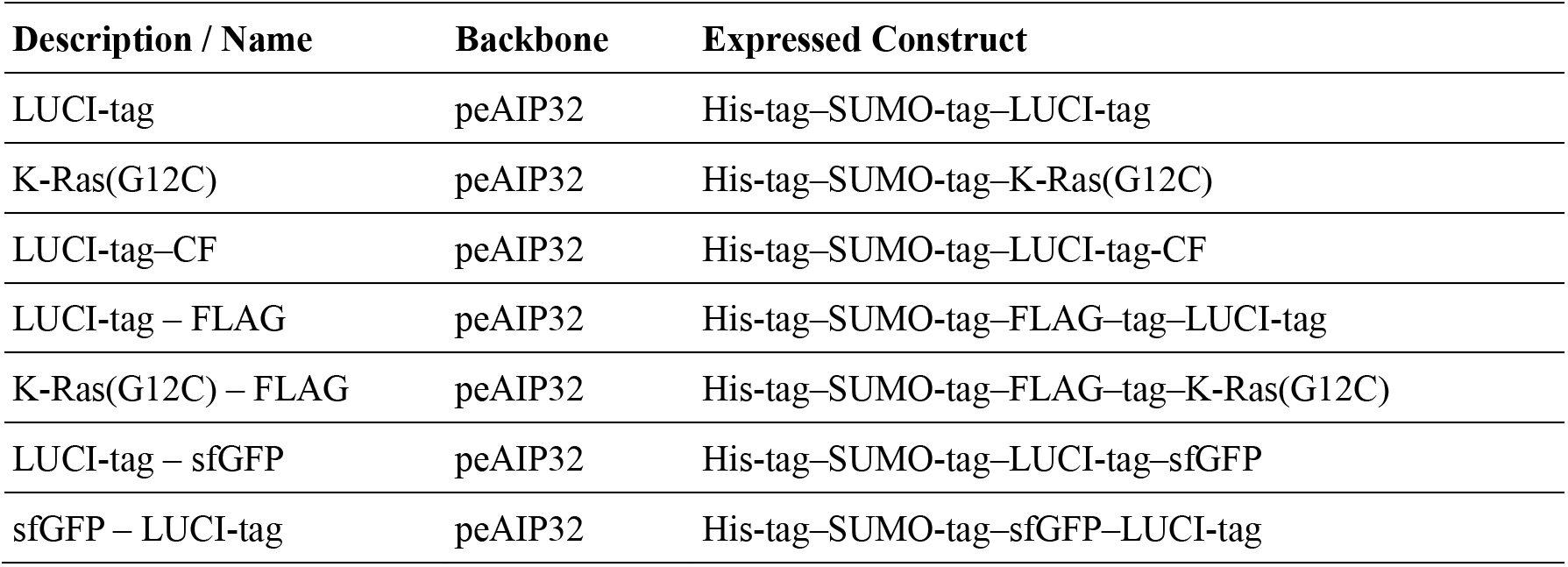
Bacterial Plasmids.

**Table S10.**
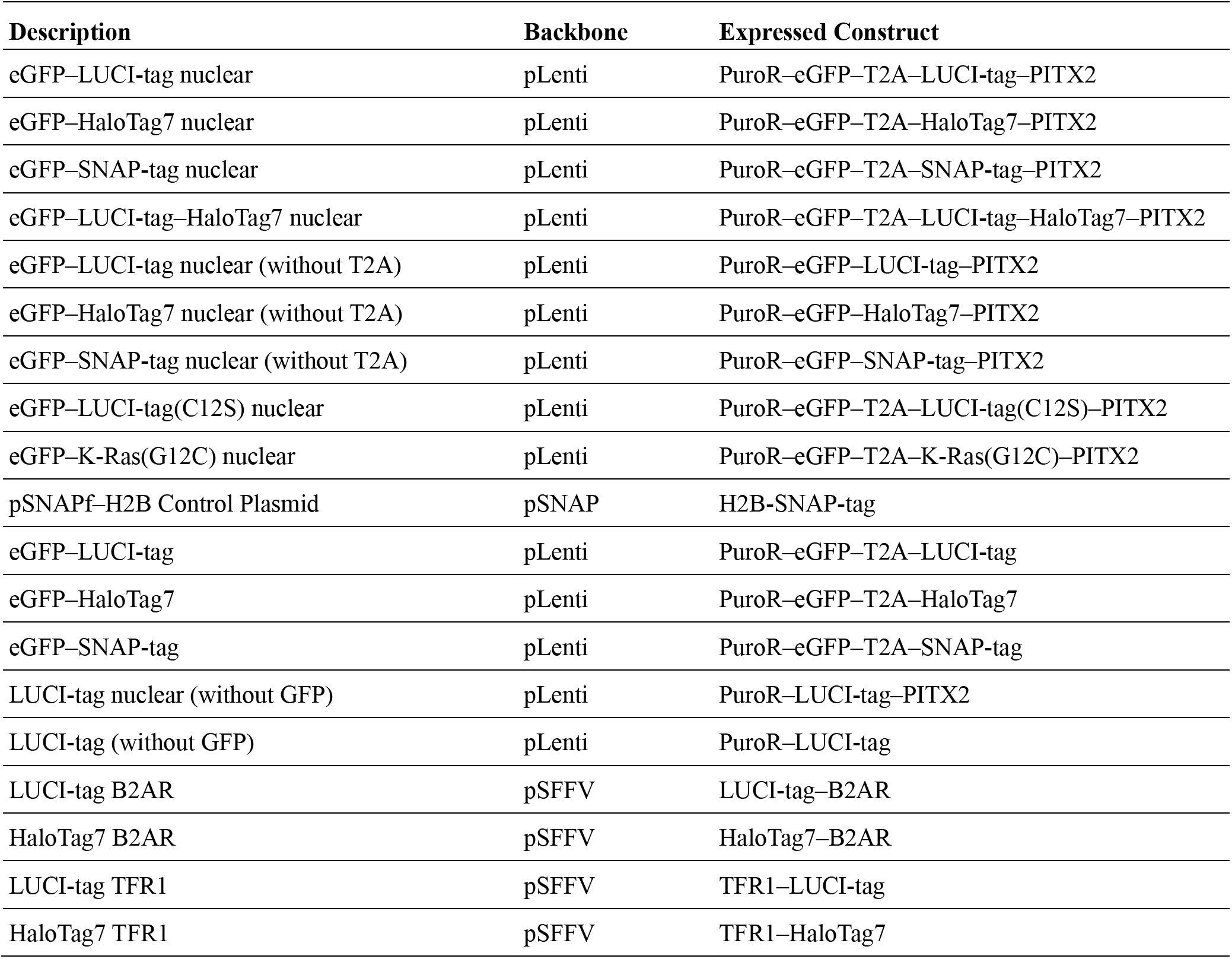
Mammalian Plasmids.

**Table S11.**
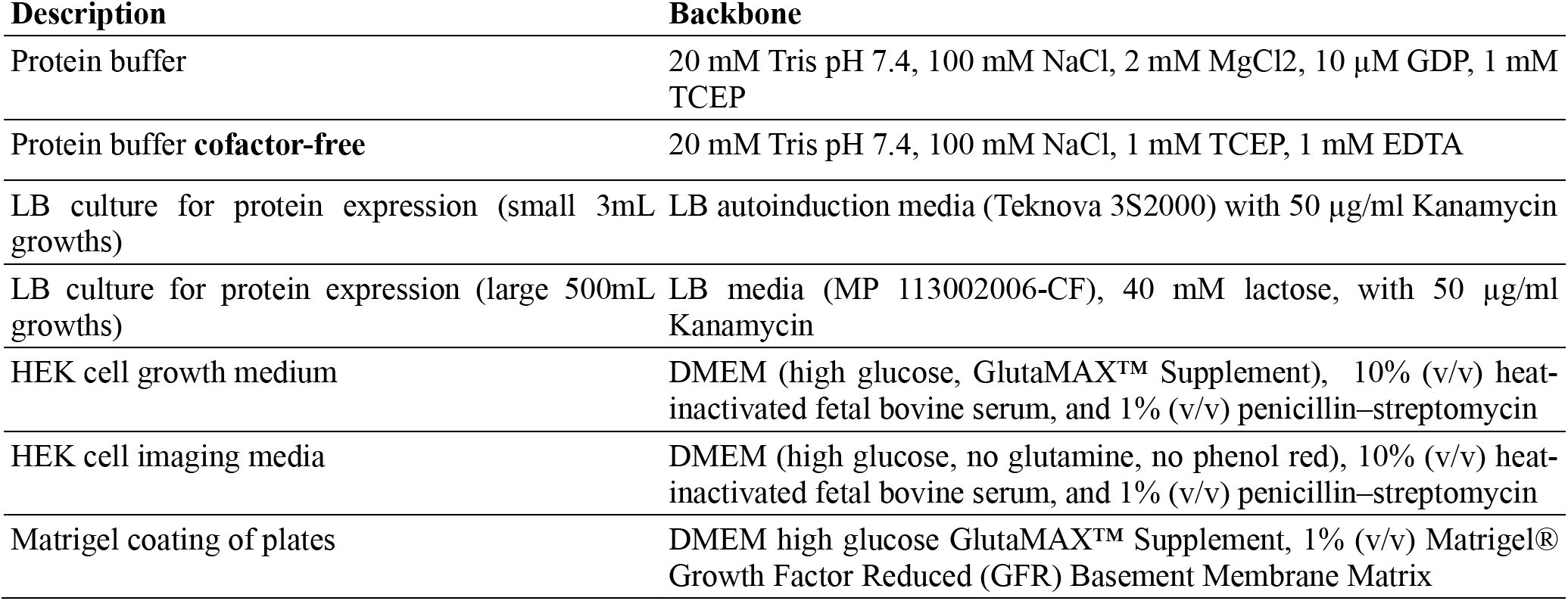
Buffers, media and reagents.

**Table S12.**
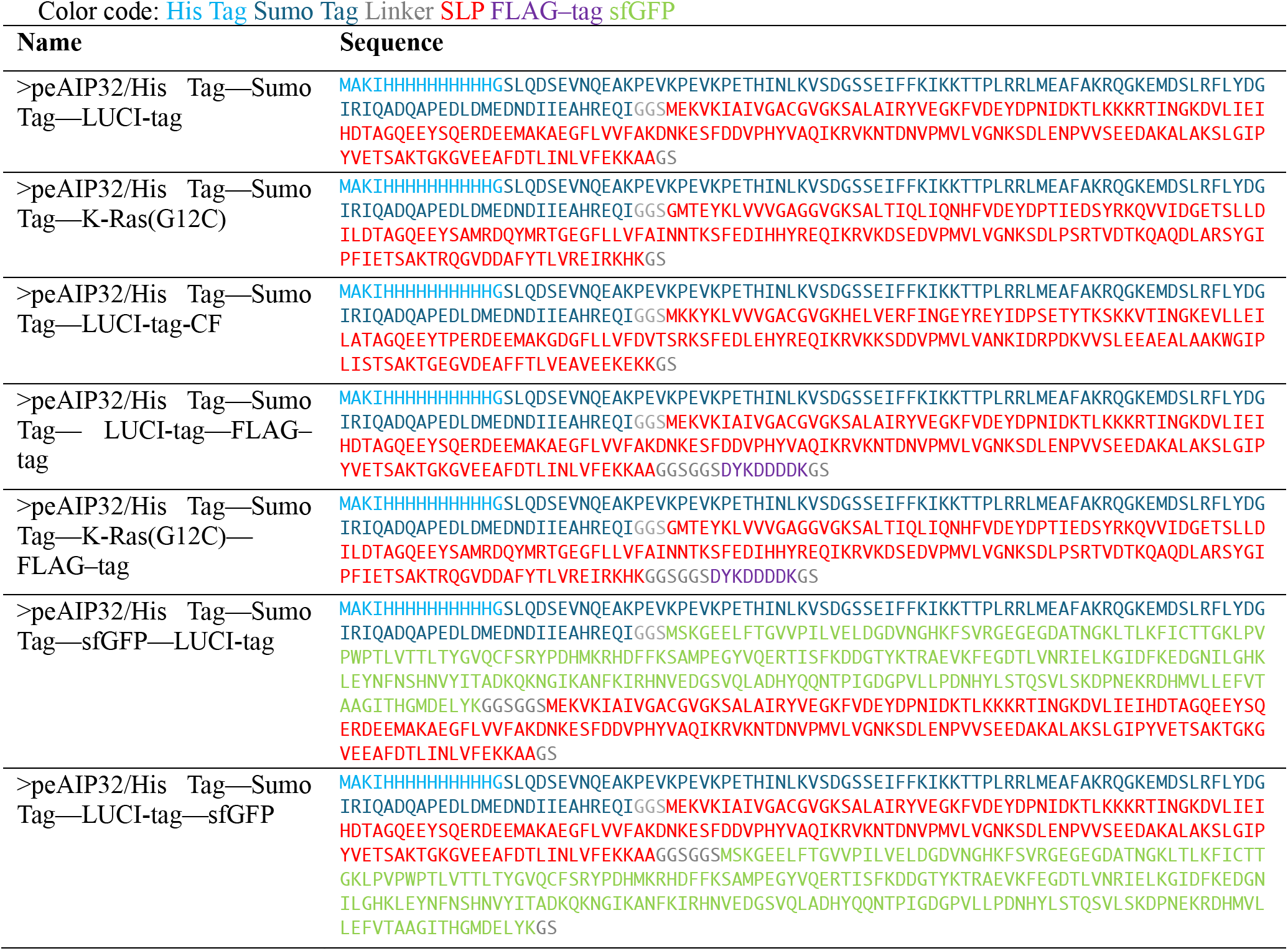
Bacterial Expression Sequences.

**Table S13.**
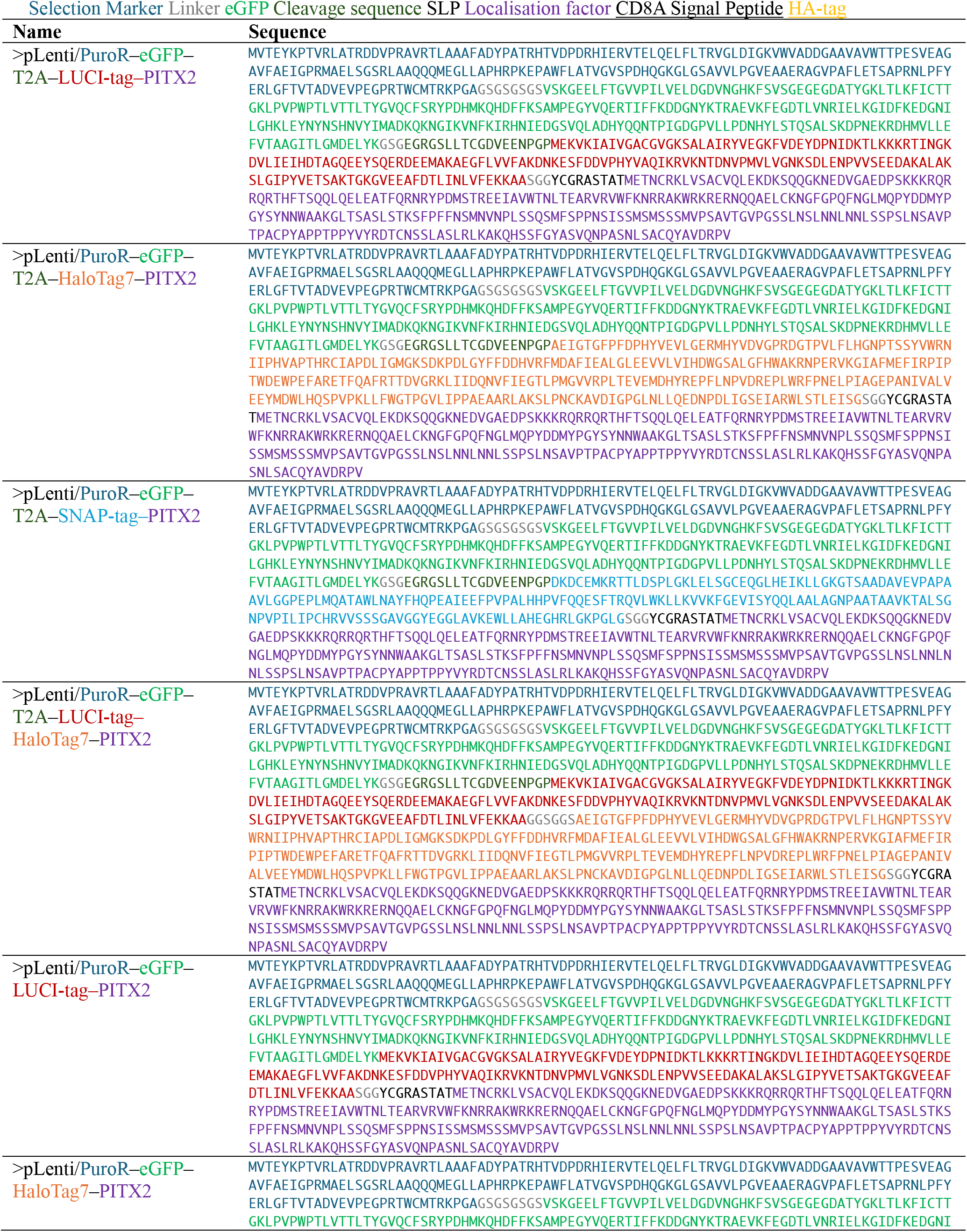
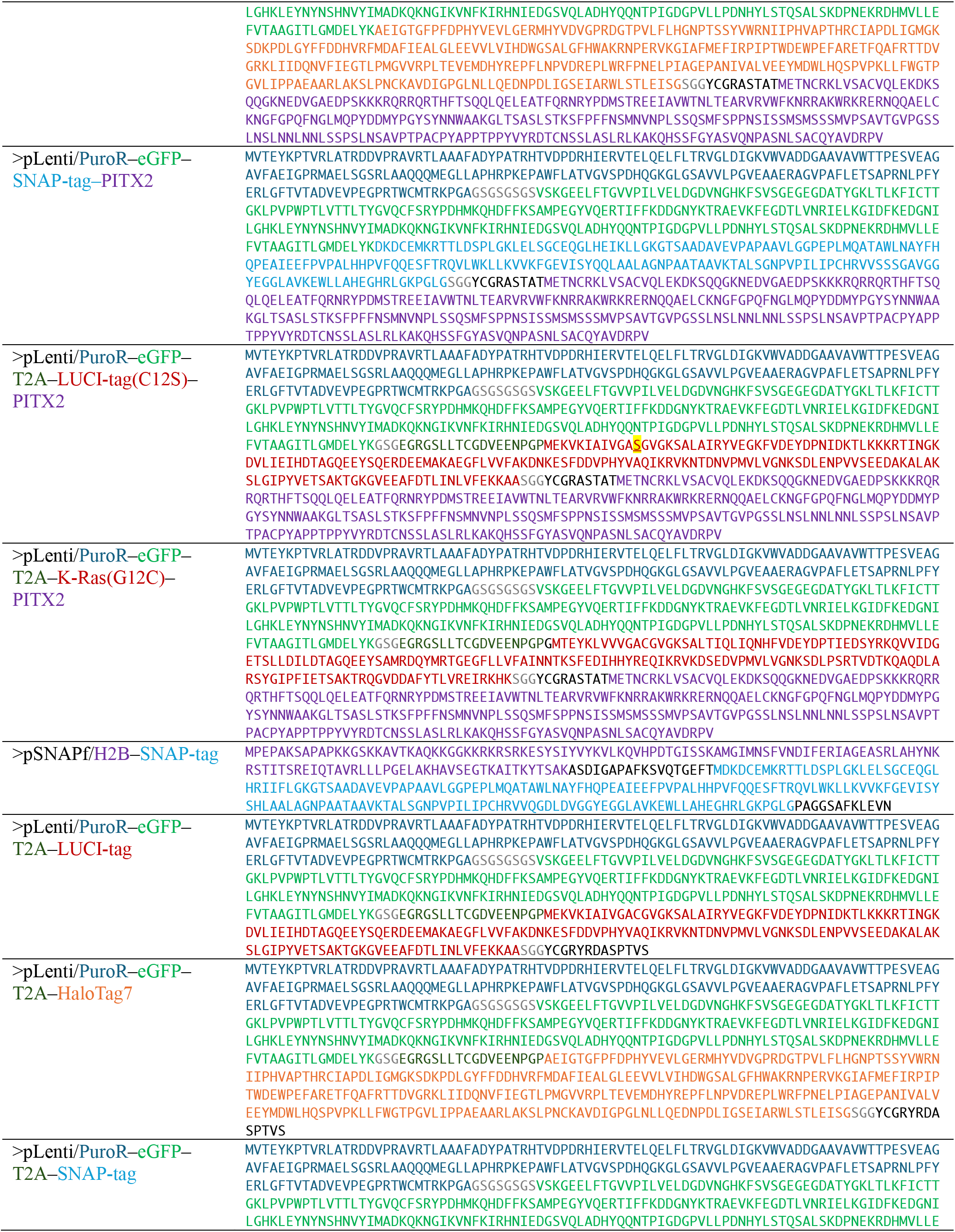
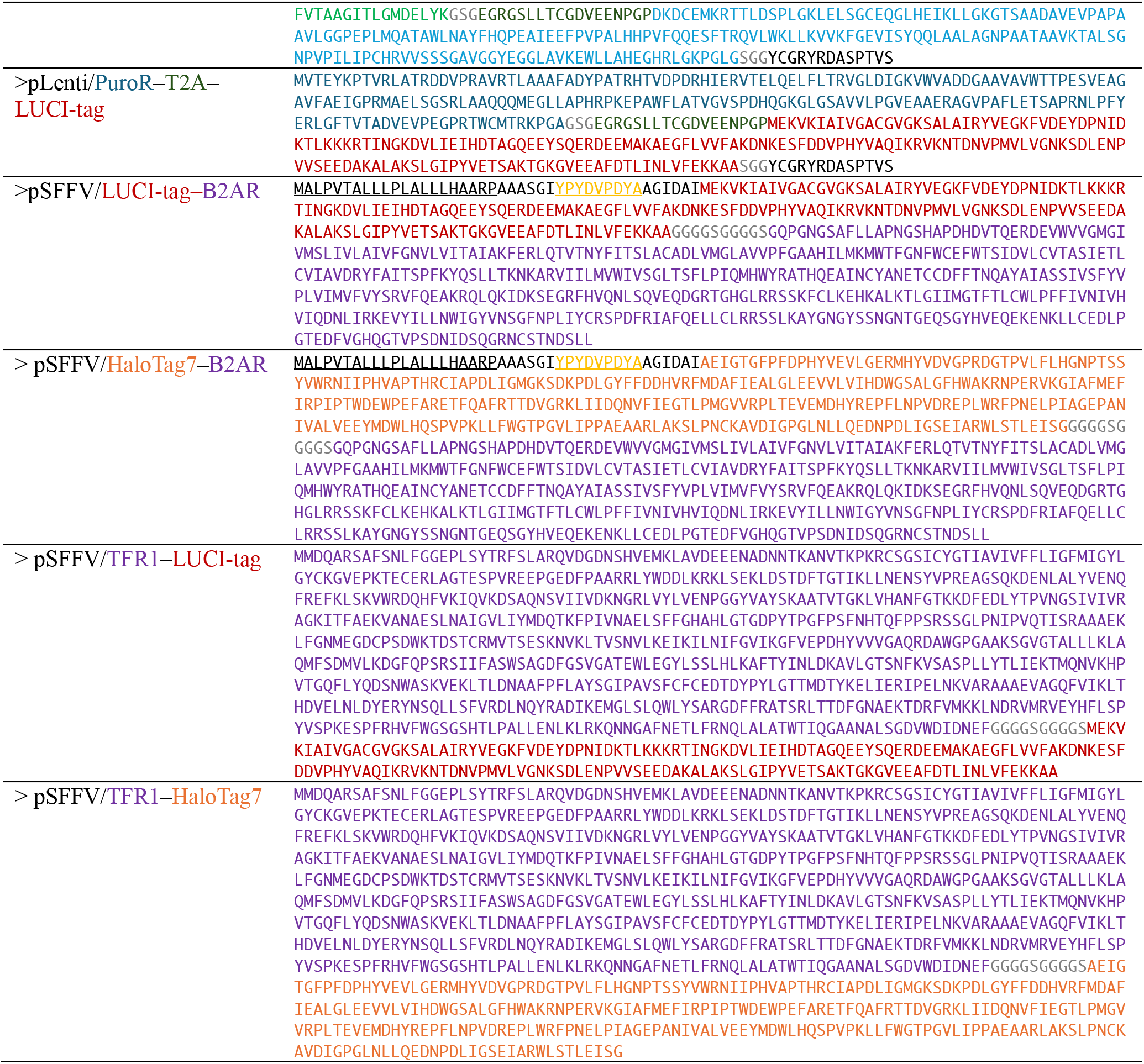
Mammalian Expression Sequences.

## References

1. Porzberg, N., Gries, K. & Johnsson, K. Exploiting covalent chemical labeling with self-labeling proteins. Annu. Rev. Biochem. 94, 29–58 (2025).

2. Ohana, R. F., Encell, L. P., Zhao, K., Simpson, D., Slater, M. R., Urh, M. & Wood, K. V. HaloTag7: a genetically engineered tag that enhances bacterial expression of soluble proteins and improves protein purification. Protein Expr. Purif. 68, 110–120 (2009).

3. Encell, L. P., Friedman Ohana, R., Zimmerman, K., Otto, P., Vidugiris, G., Wood, M. G., Los, G. V., McDougall, M. G., Zimprich, C., Karassina, N., Learish, R. D., Hurst, R., Hartnett, J., Wheeler, S., Stecha, P., English, J., Zhao, K., Mendez, J., Benink, H. A., Murphy, N., Daniels, D. L., Slater, M. R., Urh, M., Darzins, A., Klaubert, D. H., Bulleit, R. F. & Wood, K. V. Development of a dehalogenase-based protein fusion tag capable of rapid, selective and covalent attachment to customizable ligands. Curr. Chem. Genomics 6, 55–71 (2012).

4. Los, G. V., Encell, L. P., McDougall, M. G., Hartzell, D. D., Karassina, N., Zimprich, C., Wood, M. G., Learish, R., Ohana, R. F., Urh, M., Simpson, D., Mendez, J., Zimmerman, K., Otto, P., Vidugiris, G., Zhu, J., Darzins, A., Klaubert, D. H., Bulleit, R. F. & Wood, K. V. HaloTag: a novel protein labeling technology for cell imaging and protein analysis. ACS Chem. Biol. 3, 373–382 (2008).

5. Gautier, A., Juillerat, A., Heinis, C., Corrêa, I. R., Jr, Kindermann, M., Beaufils, F. & Johnsson, K. An engineered protein tag for multiprotein labeling in living cells. Chem. Biol. 15, 128–136 (2008).

6. Kühn, S., Nasufovic, V., Wilhelm, J., Kompa, J., de Lange, E. M. F., Lin, Y.-H., Egoldt, C., Fischer, J., Lennoi, A., Tarnawski, M., Reinstein, J., Vlijm, R., Hiblot, J. & Johnsson, K. SNAP-tag2 for faster and brighter protein labeling. Nat. Chem. Biol. 21, 1754–1761 (2025).

7. Mollwitz, B., Brunk, E., Schmitt, S., Pojer, F., Bannwarth, M., Schiltz, M., Rothlisberger, U. & Johnsson, K. Directed evolution of the suicide protein O^6^-alkylguanine-DNA alkyltransferase for increased reactivity results in an alkylated protein with exceptional stability. Biochemistry 51, 986–994 (2012).

8. Mo, J., Chen, J., Shi, Y., Sun, J., Wu, Y., Liu, T., Zhang, J., Zheng, Y., Li, Y. & Chen, Z. Third-generation covalent TMP-tag for fast labeling and multiplexed imaging of cellular proteins. Angew. Chem. Int. Ed. **61**, e202207905 (2022).

9. Sreekanth, V., Sindi, S. H., Chaudhary, S. K., Pergu, R., Singh, P., Karaj, E., Siriwongsup, S., Fung, J. E., Deb, A., DeCarlo, S. J., Mercer, J. A. M., Yamada, K., Rodriguez, D., Liu, D. R. & Choudhary, A. Ultrasmall chemogenetic tags with group-transfer ligands. Angew. Chem. Int. Ed. **64**, e202506997 (2025).

10. Rodriguez-Rios, M., Craigon, C., Nakasone, M. A., Sathe, G., Bond, A. G., Dorward, M., Edmonds, A. K., Norley, M. C., Arnold, R. E., Wood, P. M., Reynolds, S. J., Cresser-Brown, J. O., Marsh, G. P., Maple, H. J. & Ciulli, A. BromoCatch: a self-labelling tag platform for protein modification and live cell imaging. Nat. Commun. 17, 6406 (2026).

11. Huang, L., Guo, Z., Wang, F. & Fu, L. KRAS mutation: from undruggable to druggable in cancer. Signal Transduct. Target. Ther. 6, 386 (2021).

12. Cregg, J., Edwards, A. V., Chang, S., Lee, B. J., Knox, J. E., Tomlinson, A. C. A., Marquez, A., Liu, Y., Freilich, R., Aay, N., Wang, Y., Jiang, L., Jiang, J., Wang, Z., Flagella, M., Wildes, D., Smith, J. A. M., Singh, M., Wang, Z., Gill, A. L. & Koltun, E. S. Discovery of daraxonrasib (RMC-6236), a potent and orally bioavailable RAS(ON) multi-selective, noncovalent Tri-complex inhibitor for the treatment of patients with multiple RAS-addicted cancers. J. Med. Chem. 68, 6064–6083 (2025).

13. Fell, J. B., Fischer, J. P., Baer, B. R., Blake, J. F., Bouhana, K., Briere, D. M., Brown, K. D., Burgess, L. E., Burns, A. C., Burkard, M. R., Chiang, H., Chicarelli, M. J., Cook, A. W., Gaudino, J. J., Hallin, J., Hanson, L., Hartley, D. P., Hicken, E. J., Hingorani, G. P., Hinklin, R. J., Mejia, M. J., Olson, P., Otten, J. N., Rhodes, S. P., Rodriguez, M. E., Savechenkov, P., Smith, D. J., Sudhakar, N., Sullivan, F. X., Tang, T. P., Vigers, G. P., Wollenberg, L., Christensen, J. G. & Marx, M. A. Identification of the clinical development candidate MRTX849, a covalent KRAS^G12C^ inhibitor for the treatment of cancer. J. Med. Chem. 63, 6679–6693 (2020).

14. Lanman, B. A., Allen, J. R., Allen, J. G., Amegadzie, A. K., Ashton, K. S., Booker, S. K., Chen, J. J., Chen, N., Frohn, M. J., Goodman, G., Kopecky, D. J., Liu, L., Lopez, P., Low, J. D., Ma, V., Minatti, A. E., Nguyen, T. T., Nishimura, N., Pickrell, A. J., Reed, A. B., Shin, Y., Siegmund, A. C., Tamayo, N. A., Tegley, C. M., Walton, M. C., Wang, H.-L., Wurz, R. P., Xue, M., Yang, K. C., Achanta, P., Bartberger, M. D., Canon, J., Hollis, L. S., McCarter, J. D., Mohr, C., Rex, K., Saiki, A. Y., San Miguel, T., Volak, L. P., Wang, K. H., Whittington, D. A., Zech, S. G., Lipford, J. R. & Cee, V. J. Discovery of a covalent inhibitor of KRAS^G12C^ (AMG 510) for the treatment of solid tumors. J. Med. Chem. 63, 52–65 (2020).

15. Ostrem, J. M., Peters, U., Sos, M. L., Wells, J. A. & Shokat, K. M. K-Ras(G12C) inhibitors allosterically control GTP affinity and effector interactions. Nature 503, 548–551 (2013).

16. Bröker, J., Waterson, A. G., Smethurst, C., Kessler, D., Böttcher, J., Mayer, M., Gmaschitz, G., Phan, J., Little, A., Abbott, J. R., Sun, Q., Gmachl, M., Rudolph, D., Arnhof, H., Rumpel, K., Savarese, F., Gerstberger, T., Mischerikow, N., Treu, M., Herdeis, L., Wunberg, T., Gollner, A., Weinstabl, H., Mantoulidis, A., Krämer, O., McConnell, D. B. & W Fesik, S. Fragment optimization of reversible binding to the switch II pocket on KRAS leads to a potent, in vivo active KRAS^G12C^ inhibitor. J. Med. Chem. 65, 14614–14629 (2022).

17. Janes, M. R., Zhang, J., Li, L.-S., Hansen, R., Peters, U., Guo, X., Chen, Y., Babbar, A., Firdaus, S. J., Darjania, L., Feng, J., Chen, J. H., Li, S., Li, S., Long, Y. O., Thach, C., Liu, Y., Zarieh, A., Ely, T., Kucharski, J. M., Kessler, L. V., Wu, T., Yu, K., Wang, Y., Yao, Y., Deng, X., Zarrinkar, P. P., Brehmer, D., Dhanak, D., Lorenzi, M. V., Hu-Lowe, D., Patricelli, M. P., Ren, P. & Liu, Y. Targeting KRAS mutant cancers with a covalent G12C-specific inhibitor. Cell 172, 578–589.e17 (2018).

18. Endres, N. F. et al. Discovery and characterization of divarasib (GDC-6036), a potent covalent inhibitor of KRAS G12C. J. Med. Chem. 69, 5147–5165 (2026).

19. Huynh, M. V., Parsonage, D., Forshaw, T. E., Chirasani, V. R., Hobbs, G. A., Wu, H., Lee, J., Furdui, C. M., Poole, L. B. & Campbell, S. L. Oncogenic KRAS G12C: Kinetic and redox characterization of covalent inhibition. J. Biol. Chem. 298, 102186 (2022).

20. Fry, B., Slaw, K. & Polizzi, N. F. Zero-shot design of drug-binding proteins via neural iterative selection-expansion. Nature 656, 237–249 (2026).

21. Dauparas, J., Lee, G. R., Pecoraro, R., An, L., Anishchenko, I., Glasscock, C. & Baker, D. Atomic context-conditioned protein sequence design using LigandMPNN. Nat. Methods 22, 717–723 (2025).

22. Dauparas, J., Anishchenko, I., Bennett, N., Bai, H., Ragotte, R. J., Milles, L. F., Wicky, B. I. M., Courbet, A., de Haas, R. J., Bethel, N., Leung, P. J. Y., Huddy, T. F., Pellock, S., Tischer, D., Chan, F., Koepnick, B., Nguyen, H., Kang, A., Sankaran, B., Bera, A. K., King, N. P. & Baker, D. Robust deep learning-based protein sequence design using ProteinMPNN. Science 378, 49–56 (2022).

23. Mirdita, M., Schütze, K., Moriwaki, Y., Heo, L., Ovchinnikov, S. & Steinegger, M. ColabFold: making protein folding accessible to all. Nat. Methods 19, 679–682 (2022).

24. Wohlwend, J., Corso, G., Passaro, S., Getz, N., Reveiz, M., Leidal, K., Swiderski, W., Atkinson, L., Portnoi, T., Chinn, I., Silterra, J., Jaakkola, T. & Barzilay, R. Boltz-1 democratizing biomolecular interaction modeling. *bioRxiv* (2025) doi:10.1101/2024.11.19.624167.

25. Wilhelm, J., Kühn, S., Tarnawski, M., Gotthard, G., Tünnermann, J., Tänzer, T., Karpenko, J., Mertes, N., Xue, L., Uhrig, U., Reinstein, J., Hiblot, J. & Johnsson, K. Kinetic and structural characterization of the self-labeling protein tags HaloTag7, SNAP-tag, and CLIP-tag. Biochemistry 60, 2560–2575 (2021).

26. Grimm, J. B., English, B. P., Chen, J., Slaughter, J. P., Zhang, Z., Revyakin, A., Patel, R., Macklin, J. J., Normanno, D., Singer, R. H., Lionnet, T. & Lavis, L. D. A general method to improve fluorophores for live-cell and single-molecule microscopy. Nat. Methods 12, 244– 50, (2015).

27. Condakes, M. L., Zhang, Z., Danahy, D. B., Moore, R. R., Lakkaraju, S. K., Zhuo, X., Amako, Y., Borzilleri, R. M., Balachander, S. B., Chourb, L., Civiello, R. L., Dongre, A. R., Downes, D. P., Drexler, D. M., Dudiak, B. M., Dzhekieva, L., El-Samin, M., Fink, B. E., Frederick, K., Huang, C., Khan, J., Lees, E., Levins, C. G., McCarthy, C., Mintier, G. A., Mosure, K., Parker, M. F., Powles, R., Qi, J., Ruzanov, M., Sharma, S., Sheriff, S., Singh, A. K., Stedman, J., Szapiel, N., Thompson, R. L., Vaccaro, W., Wang, T., Yang, T., You, D., Meyer, M. J., Bronson, J. J. & Stewart, M. L. Covalent inhibitor design confers activity against both GDP- and GTP-bound forms of KRAS G12C. Nat. Commun. 17, 2233 (2026).

28. Vish, K. J., Paul, M. E., Rollins, A. P. & Boggon, T. J. Optimized conditions for GTP loading of Ras. J. Biol. Chem. 301, 110923 (2025).

29. Alexander, P., Chan, A. H., Rabara, D., Swain, M., Larsen, E. K., Dyba, M., Chertov, O., Ashraf, M., Champagne, A., Lin, K., Maciag, A., Gillette, W. K., Nissley, D. V., McCormick, F., Simanshu, D. K. & Stephen, A. G. Biophysical and structural analysis of KRAS switch-II pocket inhibitors reveals allele-specific binding constraints. J. Biol. Chem. 301, 110331 (2025).

30. Spoerner, M., Herrmann, C., Vetter, I. R., Kalbitzer, H. R. & Wittinghofer, A. Dynamic properties of the Ras switch I region and its importance for binding to effectors. Proc. Natl. Acad. Sci. U. S. A. 98, 4944–4949 (2001).

31. Mondal, K., Posa, M. K., Shenoy, R. P. & Roychoudhury, S. KRAS mutation subtypes and their association with other driver mutations in oncogenic pathways. Cells 13, 1221 (2024).

32. Fernando, M. C., Craven, G. B. & Shokat, K. M. The structure of KRAS^G12C^ bound to divarasib highlights features of potent switch-II pocket engagement. Small GTPases 15, 1–7 (2024).

33. Zeng, M., Xiong, Y., Safaee, N., Nowak, R. P., Donovan, K. A., Yuan, C. J., Nabet, B., Gero, T. ., Feru, F., Li, L., Gondi, S., Ombelets, L. J., Quan, C., Jänne, P. A., Kostic, M., Scott, D. A., Westover, K. D., Fischer, E. S. & Gray, N. S. Exploring targeted degradation strategy for oncogenic KRAS^G12C^. Cell Chem. Biol. 27, 19–31.e6 (2020).

34. Tran, L., Klein, S., Juergens, D., Sharma, S., Decarreau, J., Lee, G. R., Wang, Y., Chen, W., Bera, A. K., Kang, A., Woods, J., Joyce, E., Vafeados, D. K., Roullier, N., Li, X., Liu, B., Bo, Y., Muratspahić, E., Brown, T. A., Grimm, J. B., Patel, R., Lavis, L. D., Mahamid, J., An, L. & Baker, D. De novo design of orthogonal far-red, orange, and green fluorophore-binding proteins for multiplexed imaging. Science 393, 813–819 (2026).

35. Chen, Y., Yserentant, K., Hong, K., Chen, K., Kuang, Y., Picardo, R. S., Bhowmick, A., Charles-Orszag, A., Lord, S. J., Lu, L., Hou, K., Mann, S. I., Bhattacharya, S., Horst, M., Grimm, J. B., Lavis, L. D., Mullins, R. D., DeGrado, W. F. & Huang, B. De novo pan-rhodamine binders for fluorescence microscopy from mammalian cells to extremophiles. Cell 0, (2026).

36. Mou, J. LUCI-tag design pipeline. Zenodo (2026) doi:10.5281/ZENODO.22803458.

## References

1. Ahern, W., Yim, J., Tischer, D., Salike, S., Woodbury, S. M., Kim, D., Kalvet, I., Kipnis, Y., Coventry, B., Altae-Tran, H. R., Bauer, M. S., Barzilay, R., Jaakkola, T. S., Krishna, R. & Baker, D. Atom-level enzyme active site scaffolding using RFdiffusion2. Nat. Methods 23, 96–105 (2026).

2. Huynh, M. V., Parsonage, D., Forshaw, T. E., Chirasani, V. R., Hobbs, G. A., Wu, H., Lee, J., Furdui, C. M., Poole, L. B. & Campbell, S. L. Oncogenic KRAS G12C: Kinetic and redox characterization of covalent inhibition. J. Biol. Chem. 298, 102186 (2022).

3. Kwon, J. J., Dilly, J., Liu, S., Kim, E., Bian, Y., Dharmaiah, S., Tran, T. H., Kapner, K. S., Ly, S. H., Yang, X., Rabara, D., Waybright, T. J., Giacomelli, A. O., Hong, A. L., Misek, S., Wang, B., Ravi, A., Doench, J. G., Beroukhim, R., Lemke, C. T., Haigis, K. M., Esposito, D., Root, D. E., Nissley, D. V., Stephen, A. G., McCormick, F., Simanshu, D. K., Hahn, W. C. & Aguirre, A. J. Comprehensive structure-function analysis reveals gain- and loss-of-function mechanisms impacting oncogenic KRAS activity. bioRxiv (2024) doi:10.1101/2024.10.22.618529.

4. Dauparas, J., Anishchenko, I., Bennett, N., Bai, H., Ragotte, R. J., Milles, L. F., Wicky, B. I. M., Courbet, A., de Haas, R. J., Bethel, N., Leung, P. J. Y., Huddy, T. F., Pellock, S., Tischer, D., Chan, F., Koepnick, B., Nguyen, H., Kang, A., Sankaran, B., Bera, A. K., King, N. P. & Baker, D. Robust deep learning-based protein sequence design using ProteinMPNN. Science 378, 49–56 (2022).

5. Dauparas, J., Lee, G. R., Pecoraro, R., An, L., Anishchenko, I., Glasscock, C. & Baker, D. Atomic context-conditioned protein sequence design using LigandMPNN. Nat. Methods 22, 717–723 (2025).

6. Fry, B., Slaw, K. & Polizzi, N. F. Zero-shot design of drug-binding proteins via neural iterative selection-expansion. Nature 656, 237–249 (2026).

7. Mirdita, M., Schütze, K., Moriwaki, Y., Heo, L., Ovchinnikov, S. & Steinegger, M. ColabFold: making protein folding accessible to all. Nat. Methods 19, 679–682 (2022).

8. Jumper, J., Evans, R., Pritzel, A., Green, T., Figurnov, M., Ronneberger, O., Tunyasuvunakool, K., Bates, R., Žídek, A., Potapenko, A., Bridgland, A., Meyer, C., Kohl, S. A. A., Ballard, A. J., Cowie, A., Romera-Paredes, B., Nikolov, S., Jain, R., Adler, J., Back, T., Petersen, S., Reiman, D., Clancy, E., Zielinski, M., Steinegger, M., Pacholska, M., Berghammer, T., Bodenstein, S., Silver, D., Vinyals, O., Senior, A. W., Kavukcuoglu, K., Kohli, P. & Hassabis, D. Highly accurate protein structure prediction with AlphaFold. Nature 596, 583–589 (2021).

9. Wohlwend, J., Corso, G., Passaro, S., Getz, N., Reveiz, M., Leidal, K., Swiderski, W., Atkinson, L., Portnoi, T., Chinn, I., Silterra, J., Jaakkola, T. & Barzilay, R. Boltz-1 democratizing biomolecular interaction modeling. *bioRxiv* (2025) doi:10.1101/2024.11.19.624167.

10. Waybright, T. & Stephen, A. G. Nucleotide exchange on RAS proteins using hydrolysable and non-hydrolysable nucleotides. Methods Mol. Biol. 2797, 35–46 (2024).

11. Mou, J. LUCI-tag design pipeline. Zenodo (2026) doi:10.5281/ZENODO.22803458.

12. Wilhelm, J., Kühn, S., Tarnawski, M., Gotthard, G., Tünnermann, J., Tänzer, T., Karpenko, J., Mertes, N., Xue, L., Uhrig, U., Reinstein, J., Hiblot, J. & Johnsson, K. Kinetic and structural characterization of the self-labeling protein tags HaloTag7, SNAP-tag, and CLIP-tag. Biochemistry 60, 2560–2575 (2021).

13. Winter, G. & McAuley, K. E. Automated data collection for macromolecular crystallography. Methods 55, 81– 93 (2011).

14. Adams, P. D., Afonine, P. V., Bunkóczi, G., Chen, V. B., Echols, N., Headd, J. J., Hung, L.-W., Jain, S., Kapral, G. J., Grosse Kunstleve, R. W., McCoy, A. J., Moriarty, N. W., Oeffner, R. D., Read, R. J., Richardson, D. C., Richardson, J. S., Terwilliger, T. C. & Zwart, P. H. The Phenix software for automated determination of macromolecular structures. Methods 55, 94–106 (2011).

15. Emsley, P., Lohkamp, B., Scott, W. G. & Cowtan, K. Features and development of coot. Acta Crystallogr. D Biol. Crystallogr. 66, 486–501 (2010).

16. Long, F., Nicholls, R. A., Emsley, P., Graǽulis, S., Merkys, A., Vaitkus, A. & Murshudov, G. N. AceDRG: a stereochemical description generator for ligands. Acta Crystallogr. D Struct. Biol. 73, 112–122 (2017).

17. Skowronek, P., Thielert, M., Voytik, E., Tanzer, M. C., Hansen, F. M., Willems, S., Karayel, O., Brunner, A.-D., Meier, F. & Mann, M. Rapid and in-depth coverage of the (phospho-)proteome with deep libraries and optimal window design for dia-PASEF. Mol. Cell. Proteomics 21, 100279 (2022).

18. Demichev, V., Messner, C. B., Vernardis, S. I., Lilley, K. S. & Ralser, M. DIA-NN: neural networks and interference correction enable deep proteome coverage in high throughput. Nat. Methods 17, 41–44 (2020).

19. Baek, K., Metivier, R. J., Roy Burman, S. S., Bushman, J. W., Yoon, H., Lumpkin, R. J., Ryan, J. K., Abeja, D. M., Lakshminarayan, M., Yue, H., Ojeda, S., Xiong, Y., Che, J., Verano, A. L., Schmoker, A. M., Gray, N. S., Donovan, K. A. & Fischer, E. S. Unveiling the hidden interactome of CRBN molecular glues. Nat. Commun. 16, 6831 (2025).

20. Ritchie, M. E., Phipson, B., Wu, D., Hu, Y., Law, C. W., Shi, W. & Smyth, G. K. limma powers differential expression analyses for RNA-sequencing and microarray studies. Nucleic Acids Res. 43, e47 (2015).

21. Mondal, K., Posa, M. K., Shenoy, R. P. & Roychoudhury, S. KRAS mutation subtypes and their association with other driver mutations in oncogenic pathways. Cells 13, 1221 (2024).

22. Ostrem, J. M., Peters, U., Sos, M. L., Wells, J. A. & Shokat, K. M. K-Ras(G12C) inhibitors allosterically control GTP affinity and effector interactions. Nature 503, 548–551 (2013).

23. Maciag, A. E., Stice, J. P., Wang, B., Sharma, A. K., Chan, A. H., Lin, K., Singh, D., Dyba, M., Yang, Y., Setoodeh, S., Smith, B. P., Ju, J. H., Jeknic, S., Rabara, D., Zhang, Z., Larsen, E. K., Esposito, D., Denson, J.-P., Ranieri, M., Meynardie, M., Mehdizadeh, S., Alexander, P. A., Abreu Blanco, M., Turner, D. M., Xu, R., Lightstone, F. C., Wong, K.-K., Stephen, A. G., Wang, K., Simanshu, D. K., Sinkevicius, K. W., Nissley, D. V., Wallace, E., McCormick, F. & Beltran, P. J. Discovery of BBO-8520, a first-in-class direct and covalent dual inhibitor of GTP-bound (ON) and GDP-bound (OFF) KRAS^G12C^. Cancer Discov. 15, 578–594 (2025).

24. Condakes, M. L., Zhang, Z., Danahy, D. B., Moore, R. R., Lakkaraju, S. K., Zhuo, X., Amako, Y., Borzilleri, R. M., Balachander, S. B., Chourb, L., Civiello, R. L., Dongre, A. R., Downes, D. P., Drexler, D. M., Dudiak, B. M., Dzhekieva, L., El-Samin, M., Fink, B. E., Frederick, K., Huang, C., Khan, J., Lees, E., Levins, C. G., McCarthy, C., Mintier, G. A., Mosure, K., Parker, M. F., Powles, R., Qi, J., Ruzanov, M., Sharma, S., Sheriff, S., Singh, A. K., Stedman, J., Szapiel, N., Thompson, R. L., Vaccaro, W., Wang, T., Yang, T., You, D., Meyer, M. J., Bronson, J. J. & Stewart, M. L. Covalent inhibitor design confers activity against both GDP- and GTP-bound forms of KRAS G12C. Nat. Commun. 17, 2233 (2026).

25. Ostrem, J. M. L. & Shokat, K. M. Direct small-molecule inhibitors of KRAS: from structural insights to mechanism-based design. Nat. Rev. Drug Discov. 15, 771–785 (2016).

26. Srinivasu, B. Y., Damerla, T. S., Stec, A., Zhou, Z., Engen, J. R., Westover, K. D. & Wales, T. E. Magnesium as a conformational gatekeeper of KRAS: Structural dynamics and therapeutic implications. Protein Sci. 35, e70681 (2026).

27. Hu, Z. & Marti, J. Discovering and targeting dynamic drugging pockets of oncogenic proteins: The role of magnesium in conformational changes of the G12D mutated Kirsten rat sarcoma-guanosine diphosphate complex. Int. J. Mol. Sci. 23, 13865 (2022).

28. Lanman, B. A., Allen, J. R., Allen, J. G., Amegadzie, A. K., Ashton, K. S., Booker, S. K., Chen, J. J., Chen, N., Frohn, M. J., Goodman, G., Kopecky, D. J., Liu, L., Lopez, P., Low, J. D., Ma, V., Minatti, A. E., Nguyen, T. T., Nishimura, N., Pickrell, A. J., Reed, A. B., Shin, Y., Siegmund, A. C., Tamayo, N. A., Tegley, C. M., Walton, M. C., Wang, H.-L., Wurz, R. P., Xue, M., Yang, K. C., Achanta, P., Bartberger, M. D., Canon, J., Hollis, L. S., McCarter, J. D., Mohr, C., Rex, K., Saiki, A. Y., San Miguel, T., Volak, L. P., Wang, K. H., Whittington, D. A., Zech, S. G., Lipford, J. R. & Cee, V. J. Discovery of a covalent inhibitor of KRAS^G12C^ (AMG 510) for the treatment of solid tumors. J. Med. Chem. 63, 52–65 (2020).

29. Fell, J. B., Fischer, J. P., Baer, B. R., Blake, J. F., Bouhana, K., Briere, D. M., Brown, K. D., Burgess, L. E., Burns, A. C., Burkard, M. R., Chiang, H., Chicarelli, M. J., Cook, A. W., Gaudino, J. J., Hallin, J., Hanson, L., Hartley, D. P., Hicken, E. J., Hingorani, G. P., Hinklin, R. J., Mejia, M. J., Olson, P., Otten, J. N., Rhodes, S. P., Rodriguez, M. E., Savechenkov, P., Smith, D. J., Sudhakar, N., Sullivan, F. X., Tang, T. P., Vigers, G. P., Wollenberg, L., Christensen, J. G. & Marx, M. A. Identification of the clinical development candidate MRTX849, a covalent KRAS^G12C^ inhibitor for the treatment of cancer. J. Med. Chem. 63, 6679–6693 (2020).

30. Bröker, J., Waterson, A. G., Smethurst, C., Kessler, D., Böttcher, J., Mayer, M., Gmaschitz, G., Phan, J., Little, A., Abbott, J. R., Sun, Q., Gmachl, M., Rudolph, D., Arnhof, H., Rumpel, K., Savarese, F., Gerstberger, T., Mischerikow, N., Treu, M., Herdeis, L., Wunberg, T., Gollner, A., Weinstabl, H., Mantoulidis, A., Krämer, O., McConnell, D. B. & W Fesik, S. Fragment optimization of reversible binding to the switch II pocket on KRAS leads to a potent, in vivo active KRAS^G12C^ inhibitor. J. Med. Chem. 65, 14614–14629 (2022).

31. Janes, M. R., Zhang, J., Li, L.-S., Hansen, R., Peters, U., Guo, X., Chen, Y., Babbar, A., Firdaus, S. J., Darjania, L., Feng, J., Chen, J. H., Li, S., Li, S., Long, Y. O., Thach, C., Liu, Y., Zarieh, A., Ely, T., Kucharski, J. M., Kessler, L. V., Wu, T., Yu, K., Wang, Y., Yao, Y., Deng, X., Zarrinkar, P. P., Brehmer, D., Dhanak, D., Lorenzi, M. V., Hu-Lowe, D., Patricelli, M. P., Ren, P. & Liu, Y. Targeting KRAS mutant cancers with a covalent G12C-specific inhibitor. Cell 172, 578–589.e17 (2018).

32. Endres, N. F. et al. Discovery and characterization of divarasib (GDC-6036), a potent covalent inhibitor of KRAS G12C. J. Med. Chem. 69, 5147–5165 (2026).

